# Frequent lineage-specific substitution rate changes support an episodic model for protein evolution

**DOI:** 10.1101/2020.08.25.266486

**Authors:** Neel Prabh, Diethard Tautz

## Abstract

Since the inception of the molecular clock model for sequence evolution, the investigation of protein divergence has revolved around the question of a more or less constant rate of overall sequence information change. Although anomalies in clock-like divergence are described for some proteins, nowadays, the assumption of a constant decay rate for a given protein family is taken as the null model for protein evolution. Still, so far, a systematic test of this null model has not been done at a genome-wide scale despite the databases’ enormous growth. We focus here on divergence rate comparisons between closely related lineages, since this allows clear orthology assignments by synteny and unequivocal alignments, which are crucial for the determination of substitution rate changes. Thus, we generated a high-confidence dataset of syntenic orthologs from four ape species, including humans. Further analysis revealed that despite the appearance of an overall clock-like substitution pattern, a substantial number of proteins show lineage-specific acceleration and deceleration in divergence rates, or combinations of both in different lineages. Interestingly, when aggregated, even the families showing large lineage-specific rate perturbations can show overall rate equality. Our analysis uncovers a much more dynamic history of substitution rate changes in protein families. Which invalidates a pan-genome null model of constant decay, on the one hand, but remains compatible with the existing notion that aggregated data can be reliably used to estimate species splitting time. Ultimately, our data shows that a null model of constant change is not suitable to predict the evolutionary trajectories of individual proteins.

## Introduction

The idea of a constant divergence of proteins over time has existed since the initial investigations into protein divergence, which started with examining serological evidence followed by the analysis of haemoglobin homologs [1,2]. Refinement of this idea then led to the formulation of the molecular clock hypothesis of a more or less constant decay of sequence information in genes over evolutionary time [3,4]. However, examples that violated the molecular clock pattern were also identified early on, initially in haemoglobin itself [5]. Based on an extended sampling, Goodman et al. noted “.. in contradistinction to conclusions on the constancy of evolutionary rates, the haemoglobin genes evolved at markedly non-constant rates’’, pointing out that phases of adaptation can lead to a lineage-specific change of substitution rates. Still, in cumulative studies across many genes, the molecular clock pattern is often supported and is systematically used to compile divergence times for the tree of life [6]. Hence, the question of the role of rate fluctuations has not been systematically revisited, despite the tremendous growth of the databases in the past decades. In fact, it is now usually assumed that rate constancy is the norm. This question is of special relevance for the detection of novelty in protein evolution sequence space. Novel proteins have either diverged *de novo* from non-coding sequences or through fast adaptive divergence from duplicated genes [7,8]. Since the assignment of novelty is based on non-detectability by similarity search algorithms, Weisman *et al*. (2020) have recently proposed to use the assumption of constant family-specific decay rates as a null hypothesis for judging whether a given protein family diverges simply by constant decay into non-detectability, or whether it could be a candidate for protein novelty [9]. However, the application of such a procedure would be problematic if many protein families do not adhere to a constant decay rate over time.

Due to the fast increase of genomic datasets, one could expect that systematic estimates of protein decay rates to resolve this question should be straight forward. However, it remains a non-trivial problem due to three main reasons. First, separating orthologs from paralogs is not straight forward, and it gets further complicated as one moves deeper into the phylogeny. Alignment of paralogs can create a systematic problem in divergence rate estimation [10,11]. Second, insertion and deletion within genes make alignment and recognition of substitution events less reliable [12–17]. Third, one cannot automatically scale model-based evolutionary rate estimation methods such as dN/dS analysis to the genome level, mainly because their underlying parameters are independently calculated for each gene family. Also, these methods assume that the dS evolves under neutral rates, but this assumption has been challenged [18–23]. To avoid the confounding problems around non-coding substitution rates, we focus our study on the original approach of estimating decay rates, i.e. on direct amino acid sequence comparisons.

With the availability of large datasets, alignments of protein sequences became automatised to handle such comparisons efficiently while accepting that this creates noise in the case of suboptimal alignments around indels or highly diverged regions [12–17]. Hence, getting reliable data for divergence rates requires alignment optimisation. Further, whole-genome data have shown that misalignments between duplicated copies of the genes can be a major impediment and need to be systematically addressed [10,11].

We have produced a highly curated dataset of four species from the ape phylogeny, including humans, to revisit the decay rate constancy question for a large part of these genomes’ known coding sequences. We identify lineage-specific slow and fast diverging proteins, ancestrally rapidly evolving proteins, and other proteins with complex evolutionary trajectories. We conclude that there is a high probability of acceleration and deceleration of substitution rates for many genes, even at short evolutionary time scales. Such fluctuations may for a given protein result in bursts of rapid acceleration followed by periods of strong conservation that may cancel each other. Although this can yield a long-term constant rate pattern, the actual history of protein sequence evolution can be much more complex. Hence, we conclude that the classic alternative to a null model of constant decay, namely episodic evolution [24,25], is the more appropriate model for understanding protein family evolution.

## Results

### Synteny guided ortholog identification

To study lineage-specific divergence at the amino acid residues level, we started with an identification of orthologous proteins in the extant species. To ascertain this, we chose four ape species: human, chimpanzee, gorilla and gibbon (Fig 1a). They have a well-documented evolutionary history, and their overall genome divergence is sufficiently small to ensure unambiguous alignments of proteins. Furthermore, the human genome is among the best-curated genomes and serves as a reliable reference for comparisons. We identified the orthologs of the genes annotated in humans by combining reciprocal best BLAST hits and the pairwise analysis of gene order (Fig 1b). Among the 19,976 annotated human genes in the study, we found 14,645 syntenic with chimpanzee, 14,136 with gorilla, and 13,566 with white-cheeked gibbon (Fig 1c). Note that there are large scale chromosomal rearrangements in the gibbon genome, but at the smaller-scale, it is largely comparable with the other apes [26]. In total, we found 12,621 ortholog gene families shared between the four species, which represents about two-thirds of the annotated human genes. Note that the failure to identify definite orthologs for the remainder of the genes is mostly due to duplication patterns that could not be fully resolved based on our strict filtering criteria (see Methods). Still, this constitutes the most abundant gene set comparison analysed for these species so far.

**Fig 1:**
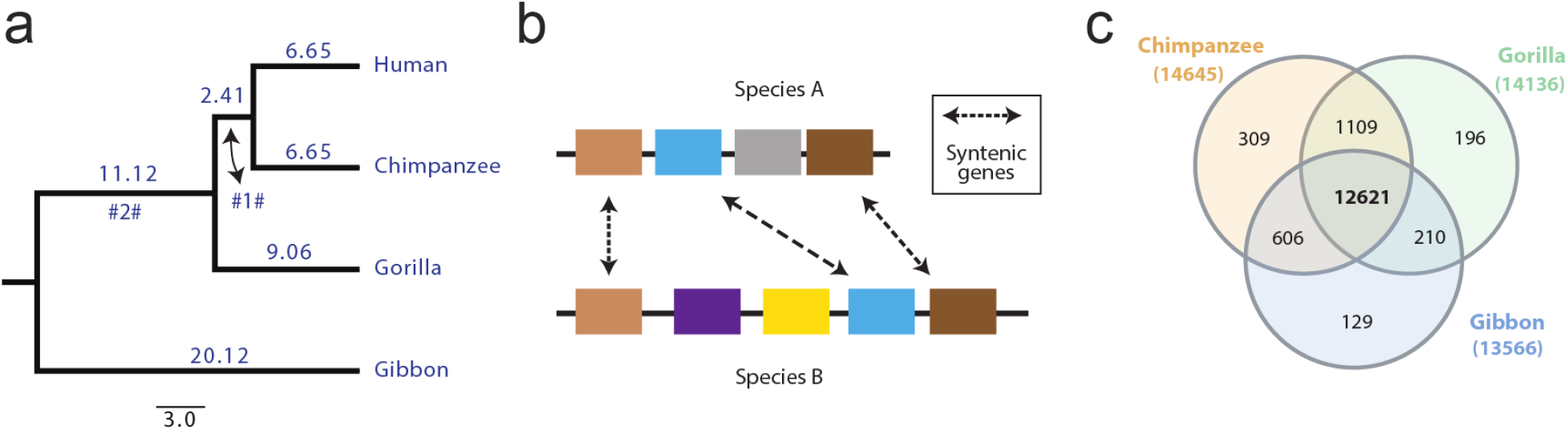
Syntenic orthologs. a) Phylogeny of apes adapted from www.timetree.org [6], branch length is divergence time in mya. #1# and #2# are the two internal branches. b) Same colour boxes represent the collinear or syntenic orthologs between the two species A and B. c) Venn diagram depicting the overlap of human genes syntenic with the other three apes.

### Optimised alignment

Proteins can diverge due to amino acid substitutions and changes in the reading frames’ length, either due to new start/stop codons or inclusion/exclusion of exons. Therefore, we have analysed how far these latter factors influence our gene set by examining the length variation between the longest and shortest orthologs from each family (Fig 2a). One-third of the ortholog families (N=4,238) had no length variation, with each ortholog having the same length, another one-third (N=4,289) had the longest orthologs that were less than 5% longer than the shortest orthologs (Fig 2b). Thus, confirming that most ortholog families in our analysis were made of proteins that do not show considerable variation in their lengths.

**Fig 2:**
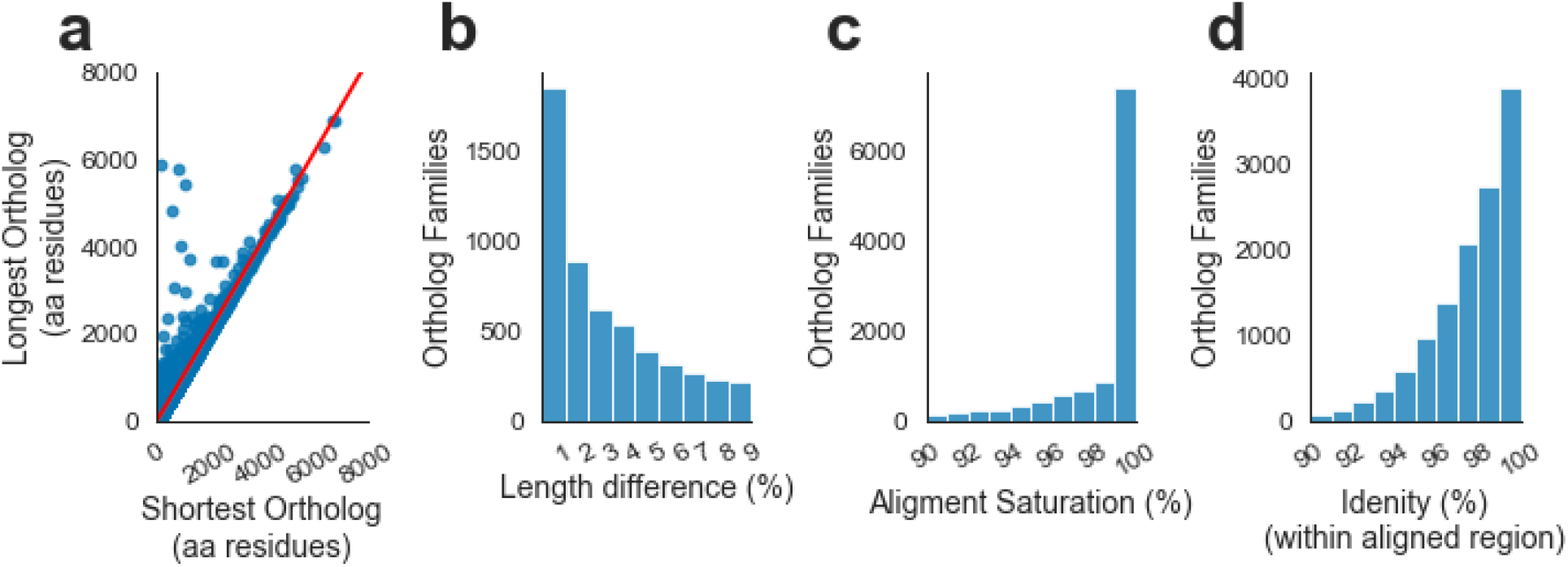
Ortholog families. a) Scatter plot of the shortest and the longest orthologs of each ortholog family. The regression line is drawn in red. b) Histogram showing ortholog family distribution for maximum length difference per 100 residues of the shortest ortholog. c) Histogram showing ortholog family distribution for alignment saturation. d) Histogram showing ortholog family distribution for identical sites per 100 aligned sites. Lower limits were excluded from the bins in panels b, c, and d.

The orthologs’ overall length similarity allowed us to employ stringent criteria (zero gap tolerance and low contiguous substitution threshold) for creating multiple sequence alignments from these families. Only three ortholog families did not overlap in the final alignment due to truncation (see S1 File); we removed these from further analysis. Thirty-five ortholog families shared less than 50 residue overlap, but we retained these. The presence of non-overlapping families suggested that our alignment protocol’s rigour could have led to the filtering of a large number of sites. So, we checked whether or not the alignments stretched across the entire length of the shortest ortholog. We estimated our alignments’ completeness by calculating the alignment saturation level, representing the fraction of the smallest ortholog retained in the final alignment. Our results show that nearly half of the ortholog families (N=5,805) had 100% alignment saturation, and only 11% of all ortholog families had less than 90% alignment saturation (Fig 2c). The mean saturation level stood at 96.5%. The observation that further bolstered the confidence in our alignments’ quality was that 88% of all ortholog families shared over 95% identity with all four species within the aligned region (Fig 2d).

### Substitution patterns

Of the aligned 7,313,620 amino acid residues, 97.5% were identical in all four species. Among the 175,284 sites with at least one substitution, 92.7% were species-specific substitutions, i.e. they were substituted in only one of the four species. Moreover, 55.8% of the substituted sites were only substituted in gibbons. Note that because no outgroup was used, the internal branch #2# (shown in Fig 1a) was added to the Gibbon branch.

12,795 sites in 5,407 protein families were substituted in more than one species, and are therefore potentially phylogenetically informative. We divided them into six categories (Fig 3a-f) based on the phylogeny. Interestingly, only about half of these sites with two states (N=6,279) were consistent with the phylogeny (Fig 3a,g), while 4,987 sites with two or three states were phylogenetically inconsistent in different combinations (Fig 3b,c). Since convergent substitutions are unlikely in this dataset, this testifies the influence of incomplete lineage sorting or introgression effects in this shallow phylogeny. As these can potentially lead to errors in rate estimation [27], and since they constitute only less than 3% of all substitute sites, we removed these sites from further analysis. The remaining sites constitute three or more states (Fig 3d-f), including the category ‘No identity’, which covered 52 sites with a different amino acid in each branch (Fig 3f), and each site was in a different ortholog family. We conjecture that these are hypermutable sites, since the estimated expected number of sites with substitution on all four species branches was only less than two (see Methods). We also removed these remaining multiple state sites from further analysis.

**Fig 3:**
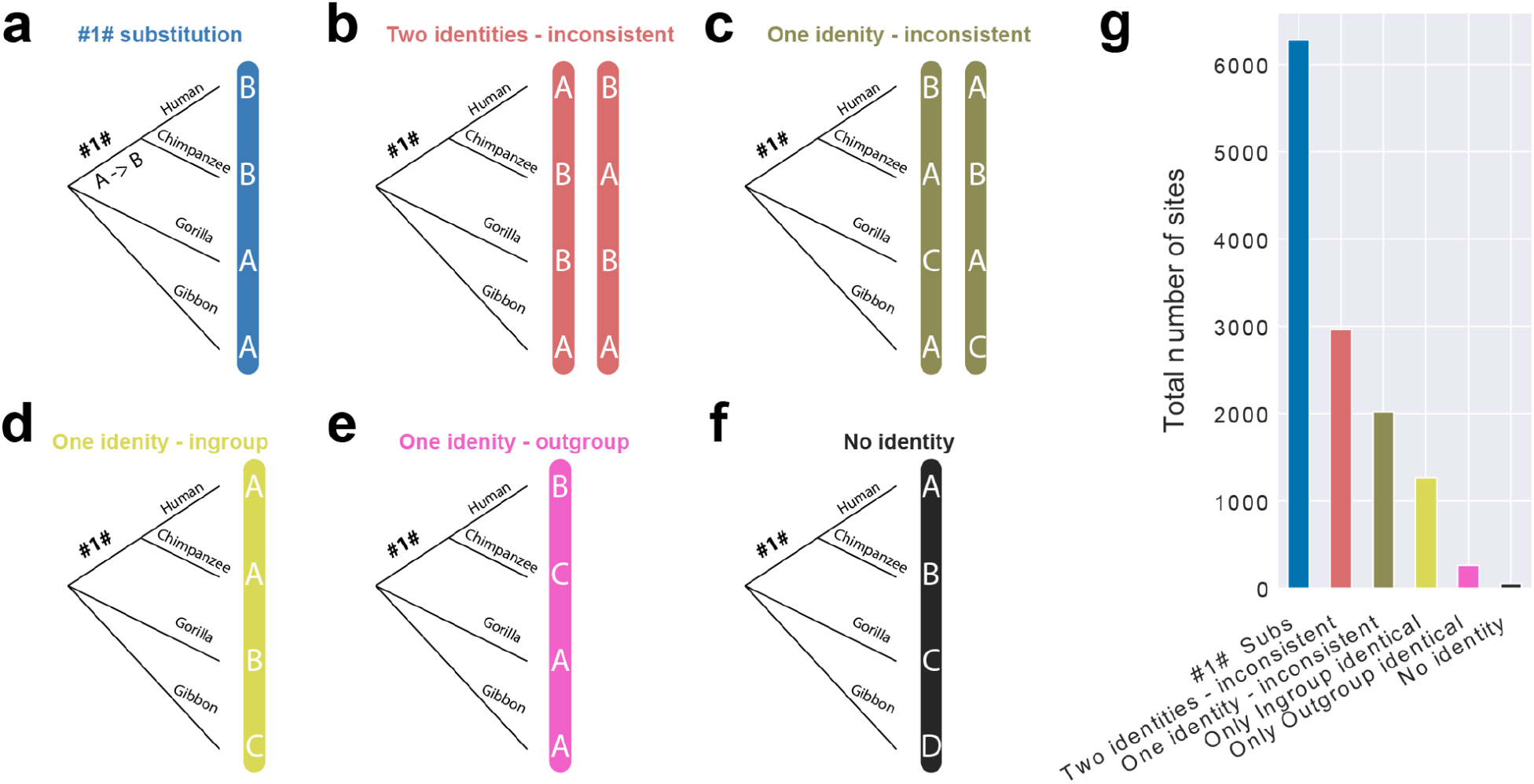
Sites substituted in more than one species. a-f) Different types of substitution patterns identified from the alignments. Letters in vertical blue bars represent residues on an aligned site. The outgroup Gibbon branch includes the #2# branch from Fig 1a. g) Bar plot representing the total number of sites based on substitution types.

### A range of decay rates among ortholog families

The branch-specific substitutions, including the #1# branch, added up to 168,768 (96.3% of all) substitutions, and we relate the further analysis only to these sites. They were used to obtain overall branch-specific substitution rates, and when scaled to the branch length from timetree, that conform to a close range of 0.036-0.043 substitutions per site per Mya (Table 1). Of course, given that the timetree branches were also calculated from molecular data, this rate consistency is not surprising per se [6,28,29]. Still, we confirm that the cumulative analysis of substitutions supports a clock-like divergence hypothesis.

**Table 1:** Branch-specific substitution rates

|  | branch timetree<br>(Mya) | total subs | % subs per site | % subs per site<br>per Mya from<br>average | Mean tree PC<br>length |
| --- | --- | --- | --- | --- | --- |
| Human | 6.65 | 19815 | 0.27 | 0.041 | 0.003 |
| Chimpanzee | 6.65 | 18818 | 0.26 | 0.039 | 0.003 |
| #1# | 2.41 | 6279 | 0.09 | 0.036 | 0.001 |
| Gorilla | 9.06 | 26026 | 0.36 | 0.039 | 0.004 |
| Gibbon | 31.24 | 97830 | 1.34 | 0.043 | 0.015 |

Moving to the individual gene families, we assessed the among family divergence rate variation in our dataset. We first removed the 1,053 families with null substitution on all five branches and retained 11,565 substituted families for further analysis (S1 Fig). We calculated Poisson corrected branch lengths for every substituted family. These were then compared to the average branch length across all families, which we call the “mean tree” (Table 1). We then used the branch length aware RF metric to compare all constituent gene family tree lengths with the mean tree length and used a Z-test for testing significant differences [30] (details in Methods). We found that 73% of families showed a significant departure from the average tree, testifying that overall rates depicted in Table 1 constitute a mixture of family-specific rates. The RF score sums absolute differences across branches and does not distinguish whether they are shorter or longer than the average. The tree length distribution is plotted in Fig 4, showing that the trees significantly different from the mean tree tend to be biased towards overall shorter trees.

**Fig 4:**
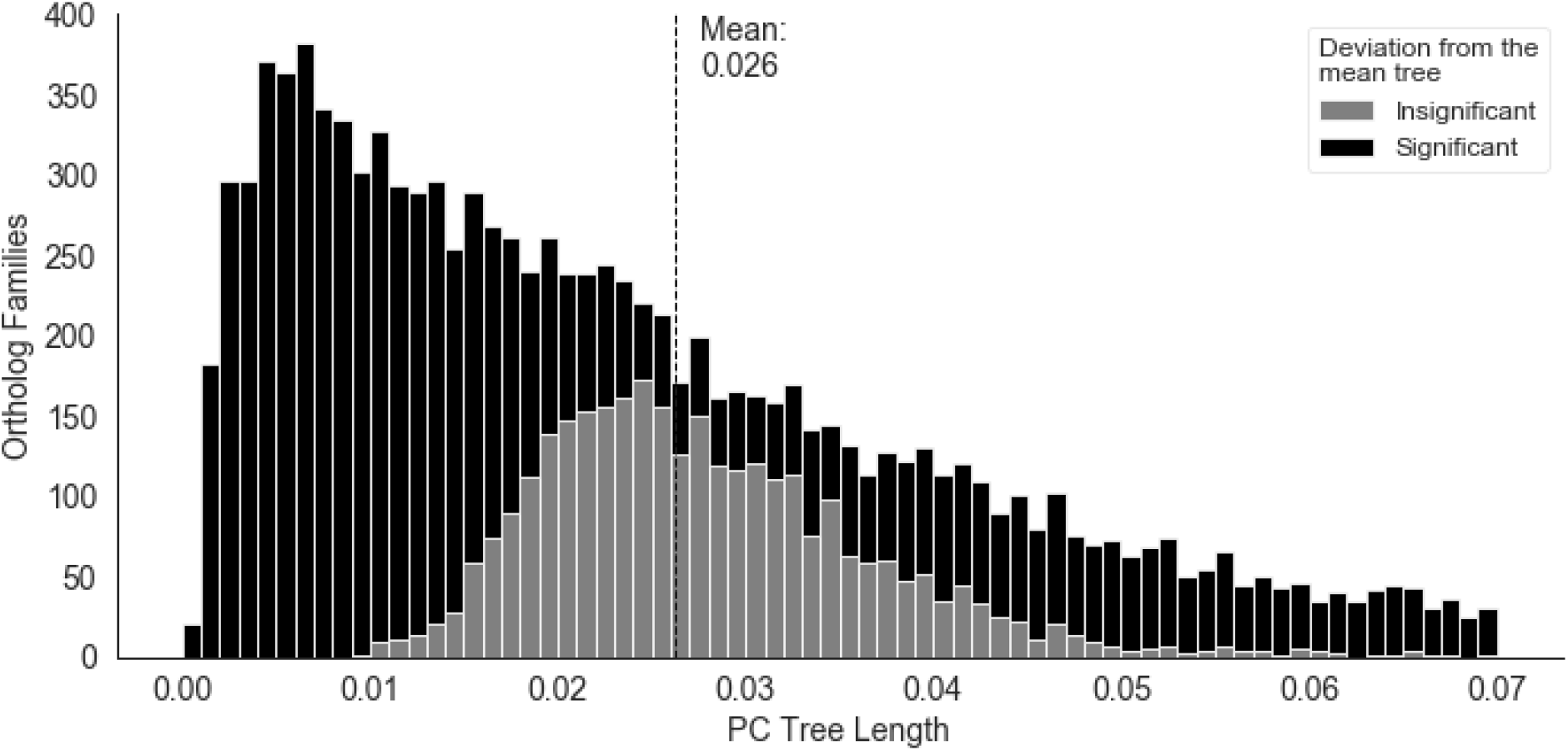
PC tree length distribution (stacked histogram), mean tree total length is marked by the dashed line. Shading represents significance of departure from the mean tree based on the Z-statistic for the branch length aware RF score [30].

### Lineage-specific rate fluctuations

The RF score does not distinguish between an average rate difference versus strong branch-specific deviations. To detect significant deviations in the lineage-specific divergence rate of proteins, we needed to take a different approach. We recognised three main issues in reliable identification of branch-specific substitution rate fluctuation at the family level. First, Z-statistics for departure from mean rates could not be calculated for null substitution branches as they have zero variance. To compensate for this, we eliminated null substitutions by artificially adding one substitution on each branch in every substituted family and recalculated the Poisson corrected branch lengths (Fig 5a). Second, to account for the family divergence rate, we normalised the Poisson corrected length on all branches with each family’s total tree length (Fig 5b, see Methods). Third, the median number of (uncorrected) substitutions on all three hominid species branches (human, chimpanzee, and gorilla) and the #1# branch was one and zero, respectively (S2 Fig). Thus, indicating that the presence or absence of a couple of substitutions may appear as a relatively large fluctuation in the substitution rate. Here, to conservatively test lineage-specific rate fluctuations, we only selected the 3,219 families with at least four substitutions (not including the artificially added substitution) on either human, chimpanzee, #1#, or gorilla branch. This arbitrary cutoff ensured that a minimum of four substitutions was required to qualify any hominid branch as a candidate for a higher than expected number of substitutions, while none of the branches would be considered as a candidate for lower than expected number of substitution unless one of the hominid branches had accumulated four or more substitutions.

**Fig 5:**
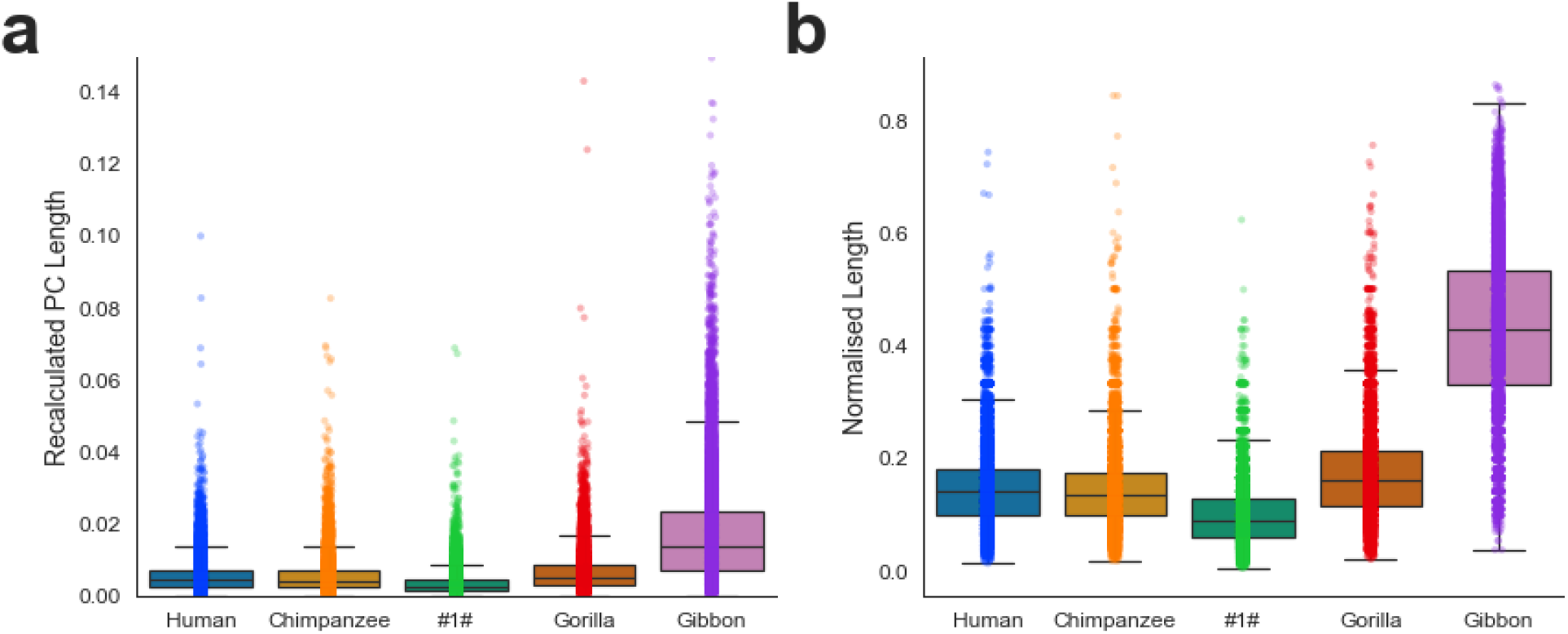
Branch-specific divergence rates after adding an extra substitution on each branch. (a) Recalculated Poisson corrected branch length distribution. (b) Within family normalised branch length distribution.

Among the selected ortholog families, we classified a branch as slow-or fast-evolving only if both its Poisson corrected and its normalised lengths were either significantly lower or higher than their respective mean lengths. We found that 677 proteins were slow-evolving in just one species, and 601 proteins were fast-evolving in just one species (Table 2, S1-S8 Tables). 121 of these were slow-evolving in one species and fast-evolving in another species; among these, the gibbon branch was fast-evolving in 78 cases. Conversely, 26 families had lower than expected substitution on the gibbon branch but higher than expected substitution in one of the other three species. Hence, more than a third of all analysed families (N=1,157) showed significant species-specific fluctuation in substitution rate.

**Table 2:** Species branches with different than expected substitutions

| Branches | Ortholog families |  |
| --- | --- | --- |
|  | Lower than expected | Higher than expected |
| Human | 208 | 42 |
| Chimpanzee | 185 | 50 |
| Gorilla | 137 | 90 |
| Gibbon | 147 | 419 |

Altogether, the fluctuating families had 5,241 and 5,353 substitutions on the human and chimpanzee branches, respectively. Under the molecular clock hypothesis, the number of unique substitutions on both human and chimpanzee branches should not differ significantly. A chi-square test indeed did not find a significant difference between the unique substitutions on the human and chimpanzee branches of the species-specific fluctuating families (see Methods), thus holding the assumption of rate equality of sequence decay. Hence, even when genes show considerable lineage-specific variations in their divergence rate, their aggregate can still follow a clock model.

### Genes evolving at a slower than expected rate

We identified over 100 lineage-specific slow-evolving proteins in every species. Among the 147 families with a lower than expected substitution on the gibbon branch, 120 families did not have higher than expected substitution on any other branch (S1 Table). For example, NELL2 (neural EGFL like 2) protein lacked a single substitution in the gibbon lineage over 826 aligned residues but had one, three, and four substitutions in human, chimpanzee, and gibbon lineage, respectively (S3 Fig). BMP8B (bone morphogenetic protein 8b), with a 402 residue alignment, had five substitutions on the gorilla branch and three substitutions on the otherwise short #1# branch but only one substitution on the gibbon branch (S4 Fig). NDUFAF6 (NADH:ubiquinone oxidoreductase complex assembly factor 6) had a complex evolutionary trajectory; it was the only protein with a slower than the expected rate on the gibbon branch and higher than the expected rate on the #1# branch (Table 3,4). All 333 residues of NDUFAF6 were retained in the final alignment, yet, the gibbon lineage had only one substitution while the #1# branch had five substitutions (S5 Fig).

**Table 3:** Genes with a complex evolutionary history

| Ortholog family /<br>Human transcript | Gene | Branch-specific substitutions |  |  |  |  | Align<br>overlap | Align sat |
| --- | --- | --- | --- | --- | --- | --- | --- | --- |
|  |  | Human | Chimp | Gorilla | Gibbon | #1# |  |  |
| ENST00000396124 | NDUFAF6 | 0 | 1 | 2 | 1 | 5 | 333 | 100 |
| ENST00000382082 | KCNV2 | 12 | 12 | 0 | 3 | 2 | 545 | 100 |
| ENST00000359741 | SLC39A14 | 2 | 0 | 1 | 4 | 17 | 485 | 99 |
| ENST00000274520 | IL9 | 0 | 0 | 0 | 6 | 4 | 144 | 100 |

**Table 4:** Genes diverging rapidly on a particular branch

| Branch | Ortholog family / Human transcript | Gene | Branch-specific % subs per site | Norm branch length | Align overlap | Align Sat | Most recent duplication in the last common ancestors of |
| --- | --- | --- | --- | --- | --- | --- | --- |
| Human | ENST00000450565 | ADCYAP1 | 7.60 | 0.74 | 171 | 97.16 | Jawed vertebrates |
| Human | ENST00000637878 | PVALEF | 3.76 | 0.40 | 133 | 99.25 | - |
| Human | ENST00000008938 | PGLYRP1 | 3.57 | 0.33 | 196 | 100.00 | Bilateral animals |
| Human | ENST00000259845 | PSORS1C2 | 2.94 | 0.42 | 136 | 100.00 | - |
| Human | ENST00000454136 | BTNL2 | 2.88 | 0.37 | 243 | 89.67 | Jawed vertebrate |
| Chimpanzee | ENST00000542996 | CALU | 6.25 | 0.84 | 320 | 99.07 | Bilateral animals |
| Chimpanzee | ENST00000296280 | MASP1 | 6.18 | 0.77 | 518 | 74.11 | Jawed vertebrates |
| Chimpanzee | ENST00000380041 | SCML1 | 5.21 | 0.36 | 326 | 99.09 | Bilateral animals |
| Chimpanzee | ENST00000342995 | CXorf67 | 3.11 | 0.29 | 386 | 97.97 | Chimpanzee |
| Chimpanzee | ENST00000343470 | LYAR | 2.65 | 0.38 | 378 | 100 | - |
| #1# | ENST00000359741 | SLC39A14 | 3.51 | 0.62 | 485 | 98.98 | Bilateral animals |
| #1# | ENST00000274520 | IL9 | 2.78 | 0.33 | 144 | 100.00 | - |
| #1# | ENST00000299191 | C16orf78/EZHIP | 2.27 | 0.22 | 264 | 100.00 | - |
| #1# | ENST00000396124 | NDUFAF6 | 1.50 | 0.43 | 333 | 100.00 | Bilateral animals |
| #1# | ENST00000433976 | ZNF778 | 1.34 | 0.20 | 598 | 98.36 | Primates |
| Gorilla | ENST00000255499 | RNF128 | 7.26 | 0.76 | 317 | 78.86 | Bilateral animals |
| Gorilla | ENST00000432056 | PRAC2 | 5.15 | 0.36 | 136 | 97.14 | - |
| Gorilla | ENST00000254976 | SNAP25 | 4.37 | 0.72 | 206 | 100.00 | Vertebrates |
| Gorilla | ENST00000613760 | WDR38 | 3.30 | 0.41 | 303 | 98.70 | Animals and Fungi |
| Gorilla | ENST00000255977 | MKRN1 | 3.17 | 0.60 | 473 | 98.13 | Simians |
| Gibbon | ENST00000513010 | S100Z | 15.22 | 0.76 | 92 | 93.88 | Vertebrates |
| Gibbon | ENST00000345088 | DPPA3 | 14.97 | 0.83 | 147 | 92.45 | Placental mammals |
| Gibbon | ENST00000398462 | SPAG11B | 13.43 | 0.68 | 67 | 62.04 | Placental mammals |
| Gibbon | ENST00000393330 | TSPAN8 | 12.66 | 0.75 | 237 | 100.00 | Bilateral animals |
| Gibbon | ENST00000523047 | SMIM23 | 12.14 | 0.79 | 140 | 96.55 | - |

The highest number of lineage-specific slow-evolving genes were found in humans, and their gene ontology analysis revealed an enrichment of genes involved in ‘otolith development’ (i.e. otoconia in mammals). The PANTHER database has eight genes associated with the biological process of otolith development, and four of them, namely LRIG1, NOX3, OTOL1, and LRIG3, had lower than expected substitutions in humans, which represented a 47.9 fold enrichment at an FDR adjusted p-value of 0.02 [31].

NOS2 (nitric oxide synthase 2) was the slowest evolving protein in our analysis with a lineage-specific perturbation restricted to humans. Although it was ranked ninth based on the normalised human branch length, the other eight proteins with lower branch lengths were fast-evolving along the gibbon lineage (S2 Table). Across the 1,153 aligned residues, NOS2 did not have a single substitution in the human lineage, while the chimpanzee lineage had four substitutions and the gorilla lineage had eight. NOS2 was associated with malaria resistance and inflammatory response [32–34].

### The fastest evolving proteins

One of the initial goals of our investigation was to identify proteins that diverge rapidly in particular lineages, since they may be indicative of a functional change of a protein that leads to an evolutionary novelty over time [35]. Genes with a significantly higher than expected number of substitutions are listed in Table 2 and S5-9 Tables. The bulk of lineage-specific fast-evolving proteins rapidly diverged in the gibbon lineage (Table 2, S8 Table). Nonetheless, even the #1# branch had eight proteins with a higher than expected number of substitutions (S9 Table).

KCNV2 was the only protein with a more than expected number of substitutions on both human and chimpanzee branches, but it was not fast diverging on the ancestral #1# branch (Table 3). We manually curated a list of the five most divergent proteins on each branch (Table 4). Three human genes, ADCYAP1, PSORS1C2 and BTNL2, are associated with neuronal phenotypes such as schizophrenia (S10 Table) [36–38]. The most closely related paralogs of these genes were traced to the common ancestor of jawed vertebrates. Thus the genes appear to be fast diverging even in the absence of recent duplication [39]. This also stands true for the top genes on the other branches. Calmuenin, RNF128, SLC39A14, and S100Z were the most divergent genes on the chimpanzee, gorilla, #1#, and gibbon branch, respectively. Also, IL9, the second-fastest diverging protein on the #1# branch, did not have a single human-, chimpanzee, or gorilla-specific substitution (Table 3), indicating that the rapid divergence on the ancestral branch was followed by absolute conservation along both descendent lineages.

ADCYAP1 is the most divergent human gene in our analysis. A previous study has shown that the gene went through accelerated adaptive evolution [40]. However, in the absence of genome sequence from other species, they did not compare its evolutionary rate with other genes. The gene encodes a neuropeptide Pituitary adenylate cyclase-activating polypeptide (PACAP). PACAP, along with its receptor PAC1 (ADCYAP1R1), plays a crucial role in regulating behavioural and cellular stress response [41]. PACAP is known to stimulate adenylate cyclase in pituitary cells and promote neuron projection [42]. ADCYAP1 has biased expression in appendix, brain, gall bladder, testis, and nine other tissues [43]. Within the sites retained in the final alignment, there were 13 substitutions along the human branch and one substitution on the gibbon branch; at the nucleotide level, there were 20 human-specific and seven other substitutions within the same region (Fig 6). It is important to note here that all human-specific substitutions are A/T to G/C, implicating biased gene conversion as a possible mutational mechanism [44,45]. A comparison of both amino acid and nucleotide alignment revealed that the five amino acid residues stretch not included in the final alignment included only five nucleotide substitutions. However, they resulted in five contiguous substitutions at the protein level, causing these residues’ exclusion from the final alignment. This stretch had four human-specific amino acid residues emanating from four nucleotide substitutions, and including these five residues raised the human-specific % substitutions per site to 9.7.

**Fig 6:**
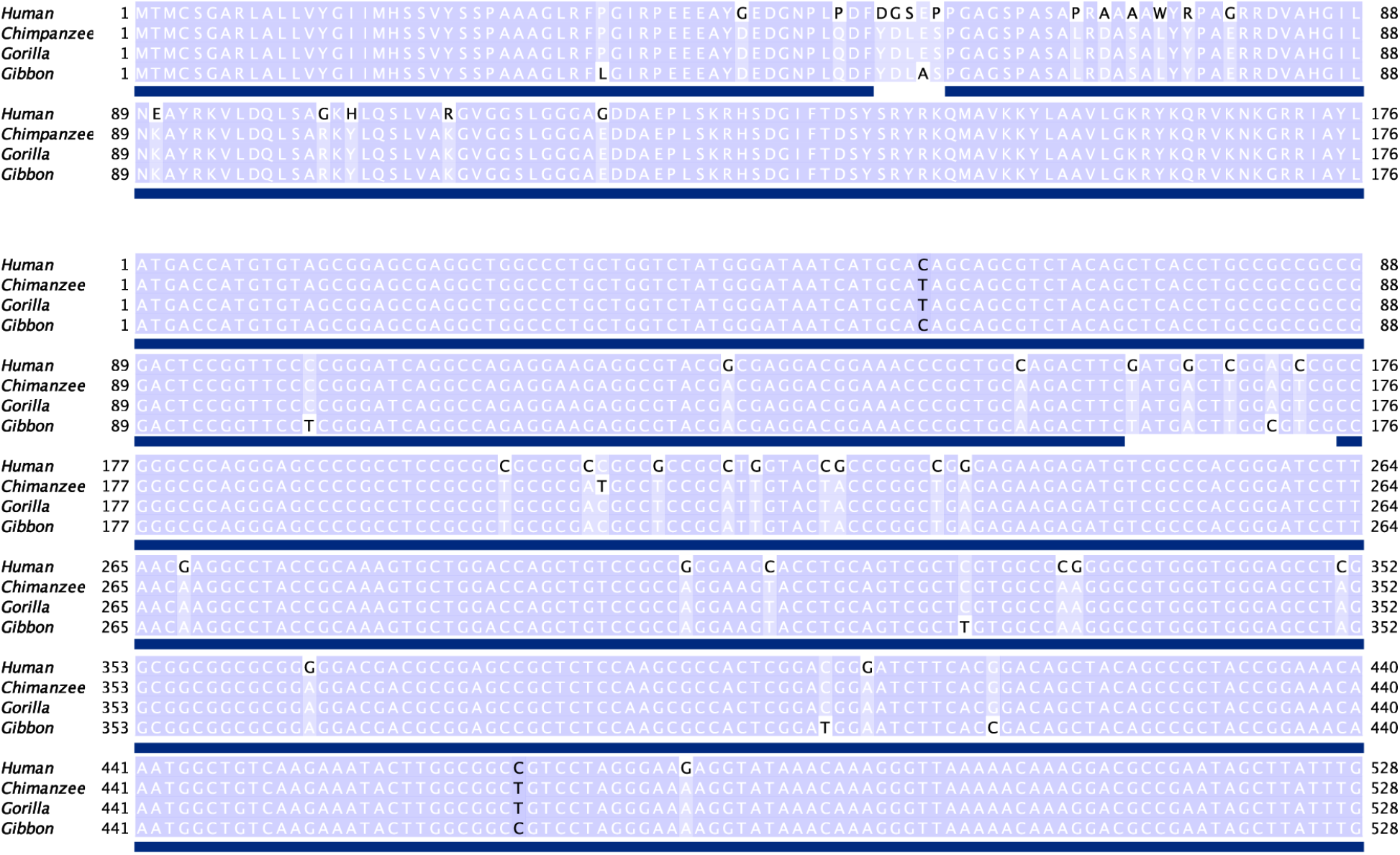
Protein and CDS multiple sequence alignments of the ADCYAP1 orthologs. Sites retained in the final alignment are underlined by the blue blocks.

## Discussion

Using the classic approach to study protein-sequence divergence rates across time, we find numerous departures from the mean substitution rates at the family-level and individual branches. Interestingly, we observed that even genes showing lineage-specific rate fluctuations can show a signature of apparent rate equality once aggregated. Given that the fluctuations include both higher divergence and higher conservation, it appears that these effects tend to cancel each other out in the aggregate data, leading to the emergence of an overall clock-like pattern. This interpretation reconciles the molecular clock departure patterns in individual protein families with the generally accepted notion that molecular data can be reliably used to derive splitting time estimates of taxa.

### Technical considerations

The tempo of protein evolution through point mutation is measured by detecting divergent sites. However, accurate identification of such sites relies heavily on a proper alignment achieved by juxtaposing conserved sites. A precise alignment of ancestrally derived sites is vital for a comparative genomic analysis involving multiple species. Thus, to conduct a thorough investigation, we carefully created a curated dataset that goes beyond the reciprocal Blast hit approach that is used to create ortholog databases across large phylogenetic distances [46]. We needed to identify true orthologs derived from the same ancestral gene of the extant species’ last common ancestor. Identification of definite orthologs even among closely related species is complicated by evolutionary processes such as deletion, duplication, and gene conversion. Our reliance on synteny to identify a set of collinear genes verified that these positional orthologs were homologous and situated at loci with conserved gene order.

Three findings validated our confidence in the chosen approach. First, most ortholog families did not show considerable variation in their protein lengths. Second, we obtained a high mean alignment saturation level even after removing all gaps. Third, there was an overwhelming abundance of identical sites within the aligned columns, and the substituted sites were heavily enriched with branch-specific substitutions. Another indication of our final alignments’ reliability was that all 52 hypermutable sites were in different families. Even a partial misalignment can easily lead to erroneous detection of multiple hypermutable sites. Hence, the lack of more than one hypermutable site in any family should be considered an additional testament to the alignment quality. Here, we posit that our rigorous approach, necessary to create a high-confidence set of ortholog families, provided an opportunity for comprehensive analysis of a large dataset.

### Overall clock-like patterns

Before analysing the individual gene families, we calibrated our dataset against a given scale. For this, we normalised our branch-specific substitution rates with their respective evolutionary time estimates from the timetree [6]. A constant rate of sequence divergence predicted by the molecular clock hypothesis should result in similar substitution rates on all branches. When scaled to the timetree, which was itself estimated from molecular data, our estimated branch-specific substitution rates confirmed that the substitutions per site per Mya falls within a close range on all branches, but it is known that some fluctuation exists in the hominid tree that leads to overall branch length changes [47,48]. However, our focus was not on these overall effects but on the lineage-specific fluctuation at the family level. Comparison of the individual family trees with the mean tree by the branch-length aware Robinson-Foulds (RF) metric revealed that nearly three-fourth of substituted families showed statistically significant departure from the mean tree [30]. Moreover, upon testing the direction of departure from the mean tree, we found that two out of three families departing from the mean tree lengths were shallower than the mean tree. This suggests that each family has its own specific rate due to evolutionary constraints.

### Lineage-specific effects in protein families

While differences in family-specific rates were expected, the key question of our analysis was the frequency of lineage-specific rate deviations in individual families. We have limited this analysis to a subset of 3,219 families that showed considerable hominid substitutions. Among these, we found approximately a third (N=1,157) that showed significant lineage-specific deceleration or acceleration of rates. For the accelerated ones, we find roughly 10 families per million years of divergence for each of the branches. Hence, if one would analyse a much deeper phylogeny, e.g. of a divergence time of 500 million years, one should conservatively expect several thousand families with phases of acceleration. If deceleration phases compensate these acceleration phases, one could still end up with an overall clock pattern for most families, but departures could also be frequent. In fact, this possibility of episodic evolution was intensively studied early on, based on mathematical considerations and simulations [24,25]. Hudson (1983) suggested, “It is concluded that the constant-rate neutral model is highly improbable”, and Gillespie (1984) record, “… our statistical analysis suggests that the course of molecular evolution is episodic…”. Interestingly, while the databases were growing, this issue had not been systematically revisited so far. Our analysis here fully supports these statements. While our analysis will still need to be confirmed in other datasets and ideally also in deeper phylogenies, it appears that the current evidence does not permit an unimpeachable assumption of a constant rate decay model for protein evolution as the null hypothesis, as recently proposed by Weisman *et al*. (2020). Their analysis was mostly guided by asking which fraction of proteins would decay with a sufficiently high rate over time to let it escape homology detection algorithms (such as BLAST). This is of particular relevance for rejecting candidate genes for de novo evolution. They concluded that a large number of genes would not have to be considered as de novo evolved when a rate calibrated from a shallow phylogeny is projected to a deeper phylogeny. However, if this shallow phylogeny included acceleration phases for the protein in question, it would yield a wrong conclusion for the long-term evolution. We conclude, therefore, that their approach is not as clear as it might seem.

### Extreme genes

Genes with extreme changes in substitutions rates are candidates for having a specific adaptive relevance for the respective taxon in whose branch they occur. In this analysis, we focused on each branch’s five fastest diverging genes, but we emphasise that the list could easily be extended (S5-9 Tables). Among the 25 highly divergent genes, five at every branch, only one was recently duplicated, confirming that rapid sequence divergence can occur even in the absence of duplication. In humans, we found that the rapidly evolving genes involved essential biological processes such as cognition. These genes were associated with disease phenotypes such as schizophrenia, autism, and blood pressure. We identified ADCYAP1 as the most divergent human protein-coding gene. It encodes a 176 amino acid residue protein that contains 17 human-specific substitutions, which estimates to a substitution frequency of 10%. The high substitution rate could have been fostered by a biased gene conversion process, as all nucleotide substitutions in humans were A/T to G/C. Despite the high divergence, PACAP, the product of ADCYAP1, remains a key mediator of the behavioural and cellular stress response, and the gene shows biased expression in appendix, brain, gall bladder, testis, and nine other tissues. This gene’s biological relevance, coupled with the lack of recent duplicates, affirms that the accelerated divergence did not result from functional redundancy. To our knowledge, no other protein has been shown to have such a high rate of human-specific divergence. It is necessary to emphasise that each lineage includes such lineage-specific highly accelerated genes, i.e. it is not special for humans to find such cases, but it is a general pattern that accompanies species formation.

### Conclusion

Our analysis reveals a substantially dynamic history of substitution rate changes in surprisingly many protein families. It appears that the episodic acceleration and deceleration cancel each other out in the aggregated data over extended times. While this could give the impression of a long-term constant rate, which is often assumed as a null model for protein evolution, the actual history of the evolution of a given protein sequence turns out to be much more complex.

## Methods

### One to one orthologs

For each species, we downloaded the CDS fasta file and gff file from the Ensembl ftp server (release-98) [39]. We extracted the fasta sequence for the CDS of each gene’s longest isoform categorised as “biotype:protein-coding” in the gff file for further analysis. We translated the extracted CDS fasta sequences to obtain their corresponding protein sequence. To detect homologous genes for each pair of species in our analysis, we ran all vs all BLASTP [49]. The BLASTP result file and gff files of each species pair were provided as an input to MCScanX for synteny ascertainment [50]. MCScanX was allowed to call a collinear block if a minimum of three collinear genes were found for the species pair with a maximum gap of two genes in between (Fig 2a). Several recent studies also relied on collinearity to establish orthologous relationships [51–57].

We parsed the collinear gene pairs obtained from the MCScanX using the following method:

1. Both protein sequences from each collinear gene pair were aligned with Stretcher [58].
2. If both proteins had 95% or more sequence identity, then this syntenic gene pair was retained.
  a. Else, we checked if either gene has a better BLASTP match, based on the BLASTP bit score, with another gene from the other species. If so, we removed the gene pair.
3. If a gene was present in more than one syntenic pair, we retained the pair within the larger syntenic block (based on the number of genes within each block).
4. Gene pairs with either gene identified as a tandem duplicate by MCScanX were removed.

Thus, in the end, we were left with a list of 1:1 orthologs for the given species pair (Fig 1c).

We overlapped the list of 1:1 orthologs of human genes with chimpanzee, gorilla, and gibbon genes to determine the ortholog gene families. We retained only those gene families that have orthologs of the human genes in all three lists (Fig 1b). The length variation within the ortholog families was calculated by subtracting the shortest ortholog’s length from the longest ortholog.

### Multiple sequence alignment of orthologous gene families

To investigate protein sequence divergence caused by single nucleotide substitution, we need to align amino acid residues that are derived from the same site of their last common ancestor. Given that most gene families are of comparable length, we set out to create alignments with fewer gaps. Hence, MAFFT was run with a gap opening penalty of 3 [59]. Then to remove unreliable columns from the alignment, we used Gblocks with the following parameters [13]:

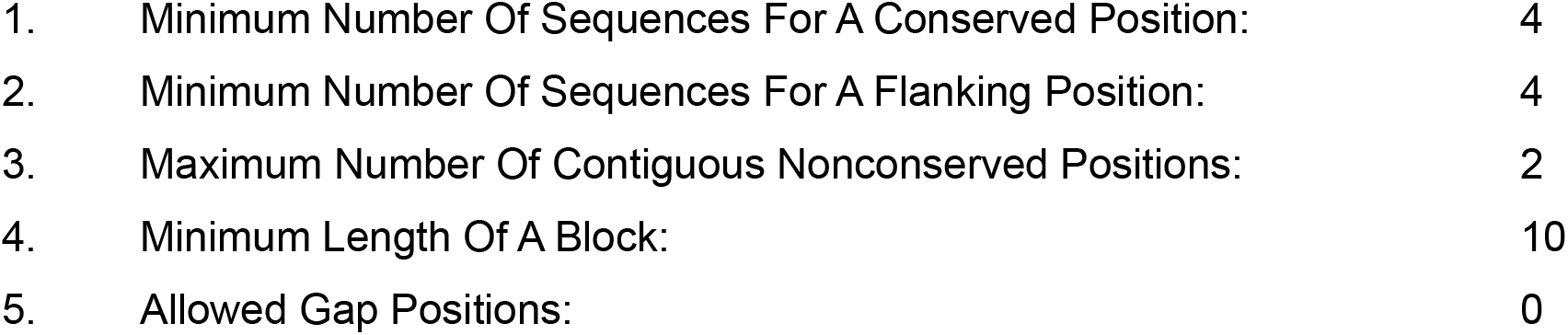

Thus, in the end, we were left with the concatenation of all the conserved blocks identified by Gblocks. These blocks were free of any gap and at most contained two substituted sites contiguously. Given the phylogenetic proximity of all four species under investigation, we assumed that it was unlikely that many instances of three or more contiguous amino acid substitutions would result from independent point substitution events. Therefore, to avoid the inclusion of insertion or deletion events within the alignment block, we have removed any gaps or contiguous substitution of three and more residues.

Since gaps are not allowed in our alignment, their maximum length is limited by the shortest ortholog. We use this qualification to measure the completeness of every alignment. If the overall alignment length is equal to the shortest ortholog’s length, this family will have attained 100% alignment saturation. The ‘Alignment Saturation’ level is calculated as per the following formula:

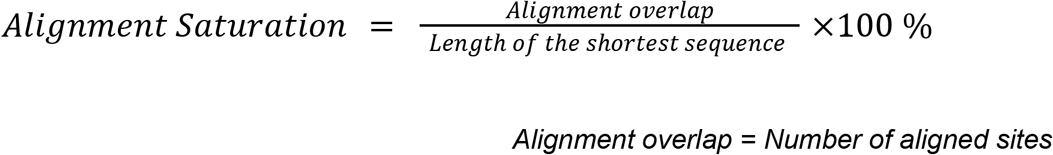

### % substitutions per site

The % substitutions per site were calculated as per the following formula:

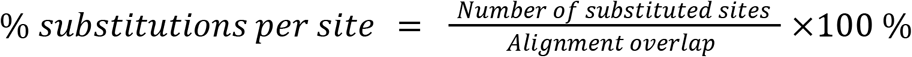

For total % substitutions per site, all substituted sites were used in the above formula. But for branch-specific substitution frequencies, only branch-specific sites were used. The branch-specific sites were identified as sites with species-specific substitution i.e. only one substitution in any given column that is specific to one of the four species sequences. #1#-specific substitutions were identified as columns with human-chimpanzee identity and gorilla-gibbon identity, as shown in Fig 3a.

We obtained the expected number of ‘No identity’ sites based on the following assumptions and calculations:

1. A ‘No identity’ site must undergo at least three independent substitution events.
2. The three lineages with the highest substitution rates in our analysis were gibbon, gorilla, and #1# plus human, with 0.0134, 0.0036, and 0.0036 substitutions per site, respectively.
3. Given that there were 7,313,620 aligned sites, the expected number of sites substituted on all the above three lineages is:

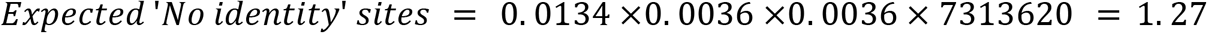

### Poisson corrected (PC) branch length

The PC length for each branch was calculated using the following formula:

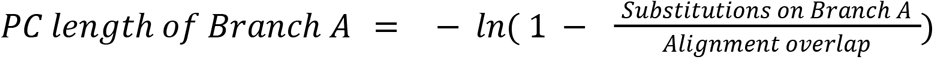

### RF metric and tree comparison

We computed the mean tree for the substitute families by obtaining the average PC length for each branch (Table 1). For every family, first, the ‘RF branch-score’ for each branch was calculated as the absolute difference (only the value of the difference, not its sign) between the PC length for the given branch of the family and the mean tree. Then the RF score for the family was obtained by adding all RF branch-scores using the following formula:

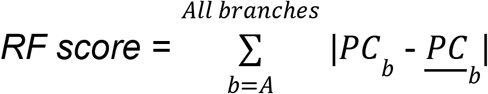

Here, *PC*_*b*_ is the Poisson corrected length of branch A for the given family and <u>PC</u> _*b*_ is the average Poisson corrected length of branch A of the mean tree (Table 1). A low RF score indicated that the family tree was close to the mean tree and thus diverged at a similar rate. We conducted a standard Z-test to evaluate if the RF score for the given family was significantly different from zero. The variance of RF score V(*z*_*i*_) was estimated from 1000 bootstrap

replications for the entire underlying alignment by using the following formula [60–63]:

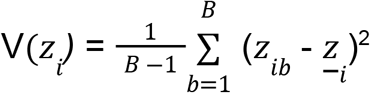

Here, *B* = 1000 (number of bootstrap replicates), *<u>z</u>* _*ib*_ is the value of *z* _*i*_ estimated at the b^th^bootstrap replication, and z_*i*_ is the average of the *z*_*ib*_.

We applied the FDR (false discovery rate) method for multiple testing correction. The statistical significance of RF scores were used to draw stacked histograms for the PC tree length distribution (Fig 4). For each family, the PC tree length was computed by adding all the branches of the given tree.

### Recalculation of Poisson Corrected branch lengths

To eliminate the null substitution branches, we recalculated the PC length for each branch using the following formula:

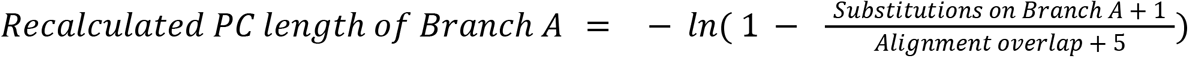

The recalculation involved appending an additional substituted column for each branch in the original underlying alignment, which increased the alignment overlap length by five residues, which was added to the denominator. The mean PC branch length was independently computed for each branch. We again conducted a standard Z-test to evaluate if the PC branch length for the given branch of every family was significantly different from its mean. The variance of recalculated length on each branch of every family was again calculated from 1000 bootstrap replications for the appended alignment by using the previously shown formula for V(*z*_*i*_ ). The same approach was used to detect departure from the mean normalised branch length calculated in the following section.

### Normalised branch-specific substitutions

To compare branches across the ortholog families, we normalised each branch of every ortholog family by using the following formula:

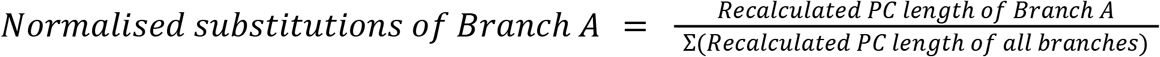

### Branch-specific fluctuation from the expected rate

To detect branch-specific rate fluctuation, we only retained families with at least four substitutions on either human, chimpanzee, gorilla, or #1# branch. Then, we tested if any branch was either significantly smaller or longer than both its mean branch-specific PC length and its mean normalised branch length. If a branch was smaller on both counts, it was considered a slow-evolving branch with a lower than expected number of substitutions. Conversely, if a branch was found to be bigger on both counts, it was considered a fast-evolving branch with a higher than expected number of substitutions.

### Molecular clock test

The chi-square statistic to test substitution rate equality between human and chimps was done by employing the following formula [64]:

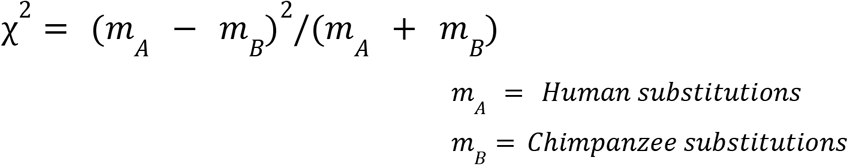

The significance of the chi-square p-value was calculated with one degree of freedom at a threshold of 0.05.

### Gene ontology enrichment analysis

The gene ontology enrichment analysis was performed using the PANTHER database on thehttp://geneontology.org/website [65].

### Identification and analysis of most divergent genes

The top five candidates on each hominid branch, from the candidates already identified to have higher than expected substitutions on the given branch, were manually curated after sorting to their % substitutions per site. Additionally, to identify the top candidates on the gibbon branch, we removed the criteria of minimum four substitutions on a hominid branch. Further, we visually inspected the alignments and removed candidates that were not fully reliable. Such filtered candidates are flagged in the ‘Comment’ section of the ‘NormDf.tsv’ file. The top 5 human candidates were validated with tissue-specific human expression data and Ensembl CDS alignment [43]. The duplicates of candidate genes were identified based on the Ensembl database’s paralogue information [39]. The nucleotide alignment of ADCYAP1 was manually created.

## Statistical analysis

Statistical analysis and tests were done using custom python codes.

## Data Availability

The data underlying this article are available in the article and its online supporting material at https://github.com/neelduti/EpisodicEvolution.

## Acknowledgements

We thank Tautz lab members, Dr. Julien Dutheil, and Dr. Chen Xie for constructive discussions. We also thank the Scientific IT group for their constant support. This work was supported by institutional funding through the Max Planck Society to DT.

## Author contributions

NP designed the study with support from DT. NP performed the analysis. NP and DT wrote the manuscript.

## Notes

https://github.com/neelduti/EpisodicEvolution

### Competing Interest Statement

The authors have declared no competing interest.

### Summary of Updates

Figures 1-5 have been revised. Supplemental files have been updated. The manuscript has been largely rewritten.

