## Supplemental Data for "Frequent lineage-specific substitution rate changes support an episodic model for protein evolution"

### Supporting information

#### Filtered ortholog families

Three ortholog families (out of 12,621) did not contain even a single alignment block. Two of these families (ENST00000262304, ENST00000301788) have truncated genes found next to genome gaps in *Nomascus leucogenys*. *Pan troglodytes* gene from ENST00000397748 family also appears to be truncated due to a genome gap. The truncated version of genes from these families do not share enough sequence overlap with their orthologs, and hence Gblocks does not find any conserved block.

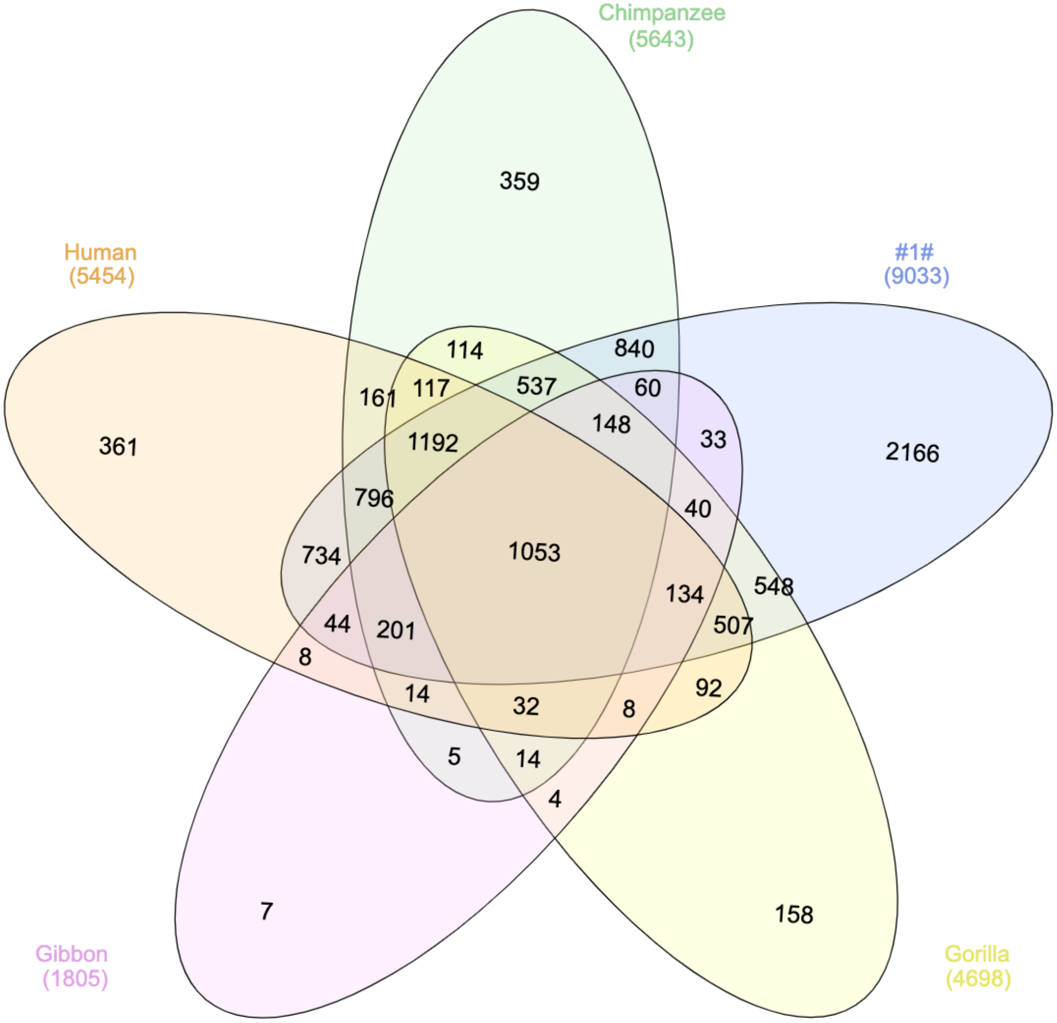


**S1 Fig**: Families with branch-specific null substitution.

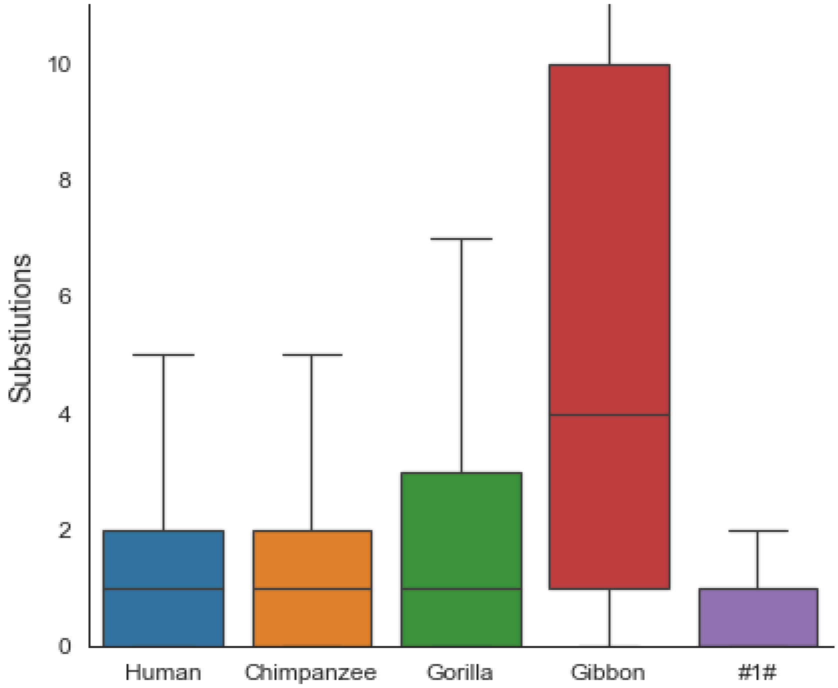


**S2 Fig**: Branch-specific substitution count interquartile range.

```
                    10        20        30        40        50        60
           =========+=========+=========+=========+=========+=========+
Human      MGFPPLLKGQASATRSSLASCSWVVFFLSCLSRHAPEIEGGRRWTELIRTMESRVLLRTF
Chimpanzee ---------------------------MSIRRLLILILKIGRRWTELIRTMESRVLLRTF
Gorilla    MGFPPLLKGQASATRSSLASCSWVVFFLSCLSRHAPEIEGGRRWTELIRTMESRVLLRTF
Gibbon     ---------------------------MSIRRLLFLILKIGRRWTELIRTMESRVLLRTF

                    70        80        90       100       110       120
           =========+=========+=========+=========+=========+=========+
Human      CLIFGLGAVWGLGVDPSLQIDVLTELELGESTTGVRQVPGLHNGTKAFLFQDTPRSIKAS
Chimpanzee CLIFGLGAVWGLGVDPSLQIDVLTELELGESTTGVRQVPGLHNGTKAFLFQDTPRSIKAS
Gorilla    CLIFGLGAVWGLGVDPCLQIDVLTELELGESTTGVRQVPGLHNGTKAFLFQDTPRSIKAS
Gibbon     CLIFGLGAVWGLGVDPSLQIDVLTELELGESTTGVRQVPGLHNGTKAFLFQDTPRSIKAS

                   130       140       150       160       170       180
           =========+=========+=========+=========+=========+=========+
Human      TATAEQFFQKLRNKHEFTILVTLKQTHLNSGVILSIHHLDHRYLELESSGHRNEVRLHYR
Chimpanzee TATAEQFFQKLRNKHEFTILVTLKQTHLNSGVIISIHHLDHRYLELESSGHRNEVRLHYR
Gorilla    TATAEQFFQKLRNKHEFTILVTLKQTHLNSGVILSIHHLDHRYLELESSGHRNEVRLHYR
Gibbon     TATAEQFFQKLRNKHEFTILVTLKQTHLNSGVILSIHHLDHRYLELESSGHRNEVRLHYR

                   190       200       210       220       230       240
           =========+=========+=========+=========+=========+=========+
Human      SGSHRPHTEVFPYILADDKWHKLSLAISASHLILHIDCNKIYERVVEKPSTDLPLGTTFW
Chimpanzee SGSHRPHTEVFPYILADDKWHKLSLAISASHLILHIDCNKIYERVVEKPSTDLPLGTTFW
Gorilla    SGSHRPHTEVFPYILADDKWHKLSLAISASHLILHIDCNKIYERVVEKPSTDLPLGTTFW
Gibbon     SGSHRPHTEVFPYILADDKWHKLSLAISASHLILHIDCNKIYERVVEKPSTDLPLGTTFW

                   250       260       270       280       290       300
           =========+=========+=========+=========+=========+=========+
Human      LGQRNNAHGYFKGIMQDVQLLVMPQGFIAQCPDLNRTCPTCNDFHGLVQKIMELQDILAK
Chimpanzee LGQRNNAHGYFKGIMQDVQLLVMPQGFIAQCPDLNRTCPTCNDFHGLVQKIMELQDILAK
Gorilla    LGQRNNAHGYFKGIMQDVQLLVMPQGFIAQCPDLNRTCPTCNDFHGLVQKIMELQDILAK
Gibbon     LGQRNNAHGYFKGIMQDVQLLVMPQGFIAQCPDLNRTCPTCNDFHGLVQKIMELQDILAK

                   310       320       330       340       350       360
           =========+=========+=========+=========+=========+=========+
Human      TSAKLSRAEQRMNRLDQCYCERTCTMKGTTYREFESWIDGCKNCTCLNGTIQCETLICPN
Chimpanzee TSAKLSRAEQRMNRLDQCYCERTCTMKGTTYREFESWIDGCKNCTCLNGTIQCETLICPN
Gorilla    TSAKLSRAEQRMNRLDQCYCERTCTMKGTTYREFESWVDGCKNCTCLNGTIQCETLICPN
Gibbon     TSAKLSRAEQRMNRLDQCYCERTCTMKGTTYREFESWIDGCKNCTCLNGTIQCETLICPN
```

```
                      370       380       390       400       410       420
             =========+=========+=========+=========+=========+=========+
Human        PDCPLKSALAYVDGKCCKECKSICQFQGRTYFEGERNTVYSSSGVCVLYECKDQTMKLVE
Chimpanzee   PDCPLKSALAYVDGKCCKECKSICQFQGRTYFEGERNTVYSSSGVCVLYECKDQTMKLVE
Gorilla      PDCPLKSALAYVDGKCCKECKSICQFQGQTYFEGERNTVYSSSGVCVLYECKDQTMKLVE
Gibbon       PDCPLKSALAYVDGKCCKECKSICQFQGRTYFEGERNTVYSSSGVCVLYECKDQTMKLVE

                      430       440       450       460       470       480
             =========+=========+=========+=========+=========+=========+
Human        SSGCPALDCPESHQITLSHSC---CKVCKGYDFCSERHNCMENSICRNLNDRAVCSCRDG
Chimpanzee   NSGCPALDCPESHQITLSHSC---CKVCKGYDFCSERRNCMENSVCRNLNDRAVCSCRDG
Gorilla      SSGCPALDCPESYQITLSHSCCKVCKVCKGYDFCSERHNCMENSVCRNLNDRAVCSCRDG
Gibbon       SSGCPALDCPESHQITLSHSC---CKVCKGYDFCSERHNCMENSVCRNLNDRAVCSCRDG

                      490       500       510       520       530       540
             =========+=========+=========+=========+=========+=========+
Human        FRALREDNAYCEDIDECAEGRHYCRENTMCVNTPGSFMCICKTGYIRIDDYSCTEHDECI
Chimpanzee   FRALREDNAYCEDIDECAEGRHYCRENTMCVNTPGSFMCICKTGYIRIDDYSCTEHDECI
Gorilla      FRALREDNAYCEDIDECAEGRHYCRENTMCVNTPGSFMCICKTGYIRIDDYSCTEHDECI
Gibbon       FRALREDNAYCEDIDECAEGRHYCRENTMCVNTPGSFMCICKTGYIRIDDYSCTEHDECI

                      550       560       570       580       590       600
             =========+=========+=========+=========+=========+=========+
Human        TNQHNCDENALCFNTVGGHNCVCKPGYTGNGTTCKAFCKDGCRNGGACIAANVCACPQGF
Chimpanzee   TNQHNCDENALCFNTVGGHNCVCKPGYTGNGTTCKAFCKDGCRNGGACIAANVCACPQGF
Gorilla      TNQHNCDENALCFNTVGGHNCVCKPGYTGNGTTCKAFCKDGCRNGGACIAANVCACPQGF
Gibbon       TNQHNCDENALCFNTVGGHNCVCKPGYTGNGTTCKAFCKDGCRNGGACIAANVCACPQGF

                      610       620       630       640       650       660
             =========+=========+=========+=========+=========+=========+
Human        TGPSCETDIDECSDGFVQCDSRANCINLPGWYHCECRDGYHDNGMFSPSGESCEDIDECG
Chimpanzee   TGPSCETDIDECSDGFVQCDSRANCINLPGWYHCECRDGYHDNGMFSPSGESCEDIDECG
Gorilla      TGPSCETDIDECSDGFVQCDSRANCINLPGWYHCECRDGYHDNGMFSPSGESCEDIDECG
Gibbon       TGPSCETDIDECSDGFVQCDSRANCINLPGWYHCECRDGYHDNGMFSPSGESCEDIDECG

                      670       680       690       700       710       720
             =========+=========+=========+=========+=========+=========+
Human        TGRHSCANDTICFNLDGGYDCRCPHGKNCTGDCIHDGKVKHNGQIWVLENDRCSVCSCQN
Chimpanzee   TGRHSCANDTICFNLDGGYDCRCPHGKNCTGDCIHDGKVKHNGQIWVLENDRCSVCSCQN
Gorilla      TGRHSCANDTICFNLDGGYDCRCPHGKNCTGDCIHDGKVKHNGQIWVLENDRCSVCSCQN
Gibbon       TGRHSCANDTICFNLDGGYDCRCPHGKNCTGDCIHDGKVKHNGQIWVLENDRCSVCSCQN
```

```
                 730       740       750       760       770       780
            =========+=========+=========+=========+=========+=========+
Human       GFVMCRRMVCDCENPTVDLFCCPECDPRLSSQCLHQNGETLYNSGDTWVQNCQQCRCLQG
Chimpanzee  GFVMCRRMVCDCENPTVDLFCCPECDPRLSSQCLHQNGETLYNSGDTWVQNCQQCRCLQG
Gorilla     GFVMCRRMVCDCENPTVDLFCCPECDPRLSSQCLHQNGETLYNSGDTWVQNCQQCRCLQG
Gibbon      GFVMCRRMVCDCENPTVDLFCCPECDPRLSSQCLHQNGETLYNSGDTWVQNCQQCRCLQG

                 790       800       810       820       830       840
            =========+=========+=========+=========+=========+=========+
Human       EVDCWPLPCPDVECEFSILPENECCPRCVTDPCQADTIRNDITKTCLDEMNVVRFTGSSW
Chimpanzee  EVDCWPLPCPDVECEFSILPENECCPRCVTDPCQADTIRNDITKTCLDEMNVVRFTGSSW
Gorilla     EVDCWPLPCPDVECEFSILPENECCPRCVTDPCQADTIRNDITKTCLDEMNVVRFTGSSW
Gibbon      EVDCWPLPCPDVECEFSILPENECCPRCVTDPCQADTIRNDITKTCLDEMNVVRFTGSSW

                 850       860
            =========+=========+=========
Human       IKHGTECTLCQCKNGHICCSVDPQCLQEL
Chimpanzee  IKHGTECTLCQCKNGHICCSVDPQCLQEL
Gorilla     IKHGTECTLCQCKNGHICCSVDPQCLQEL
Gibbon      IKHGTECTLCQCKNGHICCSVDPQCLQEL
```

**S3 Fig**: Protein alignment of the NELL orthologs. Sites retained in the final alignment are underlined by the blue blocks.

```
                         10        20        30        40        50        60
               =========+=========+=========+=========+=========+=========+
Human          MTALPGPLWLLGLALCALGGGGPGLRPPPGCPQRRLGARERRDVQREILAVLGLPGRPRP
Chimpanzee     MTALPGPLWLLGLALCALGGGGPGLRPPPGCPQRRLGARERRDVQREILAVLGLPGRPRP
Gorilla        MAARPGPLWLLGLTLCALGGGGPGLRPQPGCPQRRLGARERRDVQREILAVLGLPGRPRP
Gibbon         MAALPGPLWLLGLALCALGGGGPGLRPPPGCPQRRLGARERRDVQREILAVLGLPGRPRP

                         70        80        90       100       110       120
               =========+=========+=========+=========+=========+=========+
Human          RAPPAASRLPASAPLFMLDLYHAMAG-DDDEDGAPAERRLGRADLVMSFVNMVERDRALG
Chimpanzee     RAPPAASRLPASAPLFMLDLYHAMAG-DDDEDGAPAERRLGRADLVMSFVNMVERDRALG
Gorilla        RAPPAASRLPASAPLFMLDLYHAMAG-DDDEDGAPAEQRLGRADLVMSFVNMVERDRALG
Gibbon         RAPPAASRLPASAPLFMLDLYHAMAGDDDDEDGAPAEQRLGRADLVMSFVNMVERDRALG

                        130       140       150       160       170       180
               =========+=========+=========+=========+=========+=========+
Human          HQEPHWKEFRFDLTQIPAGEAVTAAEFRIYKVPSIHLLNRTLHVSMFQVVQEQSNRESDL
Chimpanzee     HQEPHWKEFRFDLTQIPAGEVVTAAEFRIYKVPSIHLLNRTLHVSMFQVVQEQSNRESDL
Gorilla        HQEPHWKEFRFDLTQIPAGEVVTAAEFRIYKVPSIHLLNRTLHVSMFQVVQEQSNRESDL
Gibbon         HQEPHWKEFRFDLTQIPAGEAVTAAEFRIYKVPSIHLLNRTLHVSMFQVVQEQSNRESDL

                        190       200       210       220       230       240
               =========+=========+=========+=========+=========+=========+
Human          FFLDLQTLRAGDEGWLVLDVTAASDCWLLKRHKDLGLRLYVETEDGHSVDPGLAGLLGQR
Chimpanzee     FFLDLQTLRAGDEGWLVLDVTAASDCWLLKRHKDLGLRLYVETEDGHSVDPGLAGLLGQQ
Gorilla        FFLDLQTLQAGDEGWLVLDVTAASDCWLLKHHKDLGLRLYVETEDGHSVDPGLAGLLGQR
Gibbon         FFLDLQTLQAGDEGWLVLDVTAASDCWLLKRHKDLGLRLYVETEDGHSVDPGLAGLLGQR

                        250       260       270       280       290       300
               =========+=========+=========+=========+=========+=========+
Human          APRSQQPFVVTFFRASPSPIRTPRAVRPLRRRQPKKSNELPQANRLPGIFDDVHGSHGRQ
Chimpanzee     APRSQQPFVVTFFRASPSPIRTPRAVRPLRRRQPKKTNELPQANRLPGIFDDVHGSHGRQ
Gorilla        APRSQQPFVVTFFRASPSPIRTPRAVRPLRRKQPKKTNELPQANRLPGIFDDVHGSHGRQ
Gibbon         APRSQQPFVVTFFRASPSPIRTPRAVRPLRRRQPKKTNELPQANRLPGIFDDIHGSHGRQ

                        310       320       330       340       350       360
               =========+=========+=========+=========+=========+=========+
Human          VCRRHELYVSFQDLGWLDWVIAPQGYSAYYCEGECSFPLDSCMNATNHAILQSLVHLMMP
Chimpanzee     VCRRHELYVSFQDLGWLDWVIAPQGYSAYYCEGECSFPLDSCMNATNHAILQSLVHLMMP
Gorilla        VCRRHELYVSFQDLGWLDWVIAPQGYSAYYCEGECSFPLDSCMNATNHAILQSLVHLMKP
Gibbon         VCRRHELYVSFQDLGWLDWVIAPQGYSAYYCEGECSFPLDSCMNATNHAILQSLVHLMTP
```

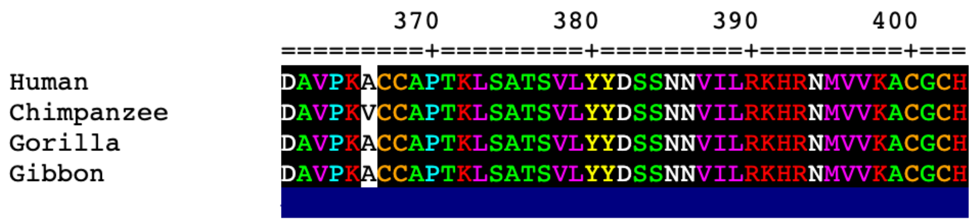


**S4 Fig**: Protein alignment of the BMP8B orthologs. Sites retained in the final alignment are underlined by the blue blocks.

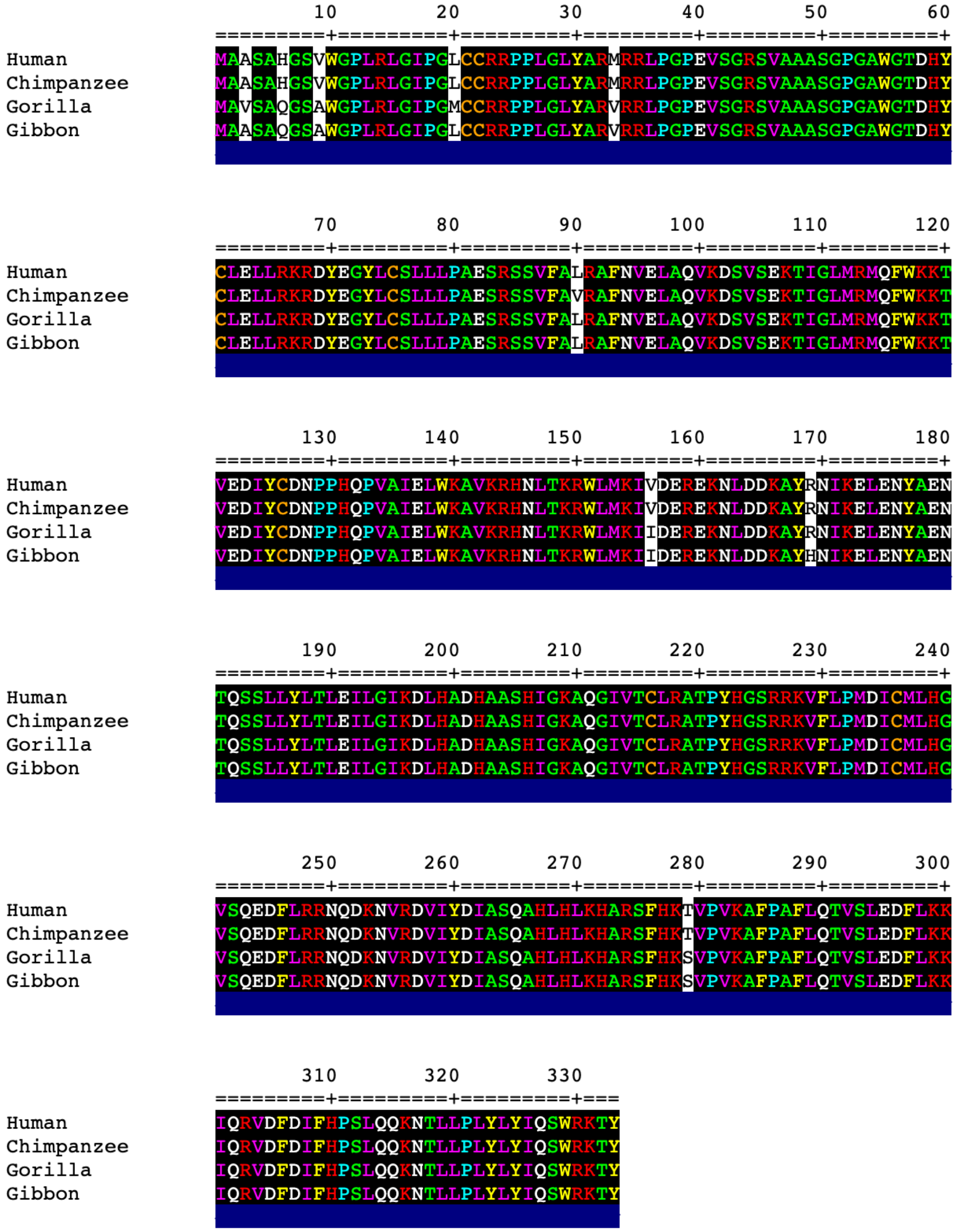


**S5 Fig**: Protein alignment of the NDUFAF6 orthologs. Sites retained in the final alignment are underlined by the blue blocks.

```
                     10        20        30        40        50        60
             =========+=========+=========+=========+=========+=========+
Human        MACPWKFLFKTKFHQYAMNGEKDINNNVEKAPCATSSPVTQDDLQYHNLSKQQNESPQPL
Chimpanzee   MACPWKFLFKTKFHQYAMNGEKDINNNVEKAPCATSSPVTQDDLQYHNLSKQQNESPQPL
Gorilla      MACPWKFLFKTKFHQYAMNGEKDINNNVEKAPCATSSPVTQDDLQYHNLSKQQNESPQPL
Gibbon       MACPWKFLFKTKFHHYAMNGEKDINNNVEKASCATSSPVTQDDLQYHRLSKQQNESPQPF

                     70        80        90       100       110       120
             =========+=========+=========+=========+=========+=========+
Human        VETGKKSPESLVKLDATPLSSPRHVRIKNWGSGMTFQDTLHHKAKGILTCRSKSCLGSIM
Chimpanzee   VETGKKSPESLVKLDATPLSSPRHVRIKNWGSGMTFQDTLHHKAKGILTCRSKSCLGSIM
Gorilla      VETGKKSPESLVKLDATPLSSPRHVRIKNWGSGMTFQDTLHHKAKGILTCRSKSCLGSIM
Gibbon       VETGKKSPESLVKLDATPLSSPRHVRIKNWGSGMTFQDTLHHKAKGILTCRSKSCLGSIM

                    130       140       150       160       170       180
             =========+=========+=========+=========+=========+=========+
Human        TPKSLTRGPRDKPTPPDELLPQAIEFVNQYYGSFKEAKIEEHLARVEAVTKEIETTGTYQ
Chimpanzee   TPKSLTRGPRDKPTPPDELLPQAIEFVNQYYGSFKEAKIEEHLARVEAVTKEIETTGTYQ
Gorilla      TPKSLTRGPRDKPTPPDELLPQAIEFVNQYYGSFKEAKIEEHLARVEAVTKEIETTGTYQ
Gibbon       TPKSLTRGPRDKPTPPDELLPQAIEFVNQYYGSFKEAKIEEHLARVEAVTKEIETTGTYQ

                    190       200       210       220       230       240
             =========+=========+=========+=========+=========+=========+
Human        LTGDELIFATKQAWRNAPRCIGRIQWSNLQVFDARSCSTAREMFEHICRHVRYSTNNGNI
Chimpanzee   LTGDELIFATKQAWRNAPRCIGRIQWSNLQVFDARSCSTAREMFEHICRHVRYSTNNGNI
Gorilla      LTGDELIFATKQAWRNAPRCIGRIQWSNLQVFDARSCSTAREMFEHICRHVRYSTNNGNI
Gibbon       LTGDELIFATKQAWRNAPRCIGRIQWSNLQVFDARSCSTAQEMFEHICRHVRYSTNNGNI

                    250       260       270       280       290       300
             =========+=========+=========+=========+=========+=========+
Human        RSAITVFPQRSDGKHDFRVWNAQLIRYAGYQMPDGSIRGDPANVEFT-------------
Chimpanzee   RSAITVFPQRSDGKHDFRVWNAQLIRYAGYQMPDGSIRGDPANVEFTQVPSPASATGHWG
Gorilla      RSAITVFPQRSDGKHDFRVWNAQLIRYAGYQMPDGSIRGDPANVEFT-------------
Gibbon       RSAITVFPQRSDGKHDFRVWNAQLIRYAGYQMPDGSIRGDPANVEFT-------------

                    310       320       330       340       350       360
             =========+=========+=========+=========+=========+=========+
Human        ------------------------------QLCIDLGWKPKYGRFDVVPLVLQANGRDPE
Chimpanzee   GEPHGERVTEWSPEETRSPGLQTHRACLVPQLCIDLGWKPKYGRFDVVPLVLQANGRDPE
Gorilla      ------------------------------QLCIDLGWKPKYGRFDVVPLVLQANGRDPE
Gibbon       ------------------------------QLCIDLGWKPKYGRFDVVPLVLQADGRDPE
```

```
                    370       380       390       400       410       420
                =========+=========+=========+=========+=========+=========+
Human           LFEIPPDLVLEVAMEHPKYEWFRELELKWYALPAVANMLLEVGGLEFPGCPFNGWYMGTE
Chimpanzee      LFEIPPDLVLEVAMEHPKYEWFRELELKWYALPAVANMLLEVGGLEFPGCPFNGWYMGTE
Gorilla         LFEIPPDLVLEVAMEHPKYEWFRELELKWYALPAVANMLLEVGGLEFPGCPFNGWYMGTE
Gibbon          LFEIPPDLVLEVAMEHPKYEWFRELELKWYALPAVANMLLEVGGLEFPACPFNGWYMGTE
```

```
                    430       440       450       460       470       480
                =========+=========+=========+=========+=========+=========+
Human           IGVRDFCDVQRYNILEEVGRRMGLETHKLASLWKDQAVVEINIAVLHSFQKQNVTIMDHH
Chimpanzee      IGVRDFCDVQRYNILEEVGRRMGLETHKLASLWKDQAVVEINIAVLHSFQKQNVTIMDHH
Gorilla         IGVRDFCDVQRYNILEEVGRRMGLETHKLASLWKDQAVVEINIAVLHSFQKQNVTIMDHH
Gibbon          IGVRDFCDVQRYNILEEVGRRMGLETHKLASLWKDQAVVEINIAVLHSFQKQNVTIMDHH
```

```
                    490       500       510       520       530       540
                =========+=========+=========+=========+=========+=========+
Human           SAAESFMKYMQNEYRSRGGCPADWIWLVPPMSGSITPVFHQEMLNYVLSPFYYYQVEAWK
Chimpanzee      SAAESFMKYMQNEYRSRGGCPADWIWLVPPMSGSITPVFHQEMLNYVLSPFYYYQVEAWK
Gorilla         SAAESFMKYMQNEYRSRGGCPADWIWLVPPMSGSITPVFHQEMLNYVLSPFYYYQVEAWK
Gibbon          SAAESFMKYMQNEYRSRGGCPADWIWLVPPMSGSITPVFHQEMLNYVLSPFYYYQVEAWK
```

```
                    550       560       570       580       590       600
                =========+=========+=========+=========+=========+=========+
Human           THVWQDEKRRPKRREIPLKVLVKAVLFACMLMRKTMASRVRVTILFATETGKSEALAWDL
Chimpanzee      THVWQDEKRRPKRREIPLKVLVKAVLFACMLMRKTMASRVRVTILFATETGKSEALAWDL
Gorilla         THVWQDEKRRPKRKEIPLKVLVKAVLFACMLMRKTMASRVRVTILFATETGKSEALAWDL
Gibbon          THVWQDEKRRPRRREIPLKVLVKAVLFACMLMRKTMASRVRVTILFATETGKSEALAWDL
```

```
                    610       620       630       640       650       660
                =========+=========+=========+=========+=========+=========+
Human           GALFSCAFNPKVVCMDKYRLSCLEEERLLLVVTSTFGNGDCPGNGEKLKKSLFMLKELNN
Chimpanzee      GALFSCAFNPKVVCMDKYRLSCLEEERLLLVVTSTFGNGDCPGNGEKLKKSLFMLKELNN
Gorilla         GALFSCAFNPKVVCMDNYRLSCLEEERLLLVVTSTFGNGDCPGNGEKLKKSLFMLKELNN
Gibbon          GALFSCAFNPKVVCMDKYRLSCLEEERLLLVVTSTFGNGDCPSNGEKLKKSLFMLKELNN
```

```
                    670       680       690       700       710       720
                =========+=========+=========+=========+=========+=========+
Human           KFRYAVFGLGSSMYPRFCAFAHDIDQKLSHLGASQLTPMGEGDELSGQEDAFRSWAVQTF
Chimpanzee      KFRYAVFGLGSSMYPRFCAFAHDIDQKLSHLGASQLTPMGEGDELSGQEDAFRSWAVQTF
Gorilla         KFRYAVFGLGSSMYPRFCAFAHDIDQKLSHLGASQLTPMGEGDELSGQEDAFRSWAVQTF
Gibbon          KFRYAVFGLGSSMYPRFCAFAHDIDQKLSHLGASQLTPTGEGDELSGQEDAFRSWAVQTF
```

```
                  730       740       750       760       770       780
         =========+=========+=========+=========+=========+=========+
Human       KAACETFDVRGKQHIQIPKLYTSNVTWDPHHYRLVQDSQPLDLSKALSSMHAKNVFTMRL
Chimpanzee  KAACEMFDVRGKQHIQIPKLYTSNVTWDPHHYRLVQDSQPLDLSKALSSMHAKNVFTMRL
Gorilla     KAACETFDVRGKQHIQIPKLYTSNVTWDPHHYRLVQDSQPLDLSKALSSMHAKNVFTMRL
Gibbon      KAACETFDVRGKQHIQIPKLYTSNVTWDPHRYRLVQDSQPLDLSKALSSMHAKNVFTMRL

                  790       800       810       820       830       840
         =========+=========+=========+=========+=========+=========+
Human       KSRQNLQSPTSSRATILVELSCEDGQGLNYLPGEHLGVCPGNQPALVQGILERVVDGPTP
Chimpanzee  KSRQNLQSPTSSRATILVELSCEDGQGLNYLPGEHLGVCPGNQPALVQGILERVVDGPTP
Gorilla     KSRQNLQSPTSSRATILVELSCEDGPGLNYLPGEHLGVCPGNQPALVQGILERVVDGPTP
Gibbon      KSRQNLQSPTSSRATILVELSCEEGQGLNYLPGEHLGVFPGNQPALVQGILERVVDNPAP

                  850       860       870       880       890       900
         =========+=========+=========+=========+=========+=========+
Human       HQTVRLEALDESGSYWVSDKRLPPCSLSQALTYFLDITTPPTQLLLQKLAQVATEEPERQ
Chimpanzee  HQTVHLEALDESGSYWVSDKRLPPCSLSQALTYFLDITTPPTQLLLLKLAQVATEATERQ
Gorilla     HQTVRLEALDESGSYWVSDKRLPPCSLSQALTYFLDITTPPTQLLLQKLAQVATEATERQ
Gibbon      HQTVRLEALDESGSYWVSDKRLPPCSLSQALTYFLDITTPPTQLLLQKLAQVATEDPERQ

                  910       920       930       940       950       960
         =========+=========+=========+=========+=========+=========+
Human       RLEALCQPSEYSKWKFTNSPTFLEVLEEFPSLRVSAGFLLSQLPILKPRFYSISSSRDHT
Chimpanzee  RLEALCQPSEYSKWKFTNSPTFLEVLEEFPSLRVSAGFLLSQLPILKPRFYSISSSRDHT
Gorilla     RLEALCQPSEYSKWKFTNSPTFLEVLEEFPSLRVSAGFLLSQLPILKPRFYSISSSRDHT
Gibbon      RLEALCQPSEYSKWKFTNSLTFLEVLEEFPSLRVSADFLLSQLPILKPRFYSISSSQDHT

                  970       980       990      1000      1010      1020
         =========+=========+=========+=========+=========+=========+
Human       PTEIHLTVAVVTYHTRDGQGPLHHGVCSTWLNSLKPQDPVPCFVRNASGFHLPEDPSHPC
Chimpanzee  PTEIHLTVAVVTYHTRDGQGPLHHGVCSTWLNSLKPQDPVPCFVRNASGFHLPEDPSHPC
Gorilla     PTEIHLTVAVLMYHTRDGQGPLHHGVCSTWLNSLKPQDPVPCFVRNASGFHLPEDPSHPC
Gibbon      PTEIHLTVAVVTYHTRDGQGPLHHGVCSTWLNSLKPQDPVPCFVRNASGFHLPEDPSHPC

                 1030      1040      1050      1060      1070      1080
         =========+=========+=========+=========+=========+=========+
Human       ILIGPGTGIAPFRSFWQQRLHDSQHKGVRGGRMTLVFGCRRPDEDHIYQEEMLEMAQKGV
Chimpanzee  ILIGPGTGIAPFRSFWQQRLHDSQHKGVRGGRMTLVFGCRRPDEDHIYQEEMLEMAQKGV
Gorilla     ILIGPGTGIAPFRSFWQQRLHDSQHKGVRGGRMTLVFGCRRPDEDHIYQEEMLEMAQKGV
Gibbon      ILIGPGTGIAPFRSFWQQRLHDSQHKGVRGGRMTLVFGCRRPDEDHIYQEEMLEMAQKGV
```

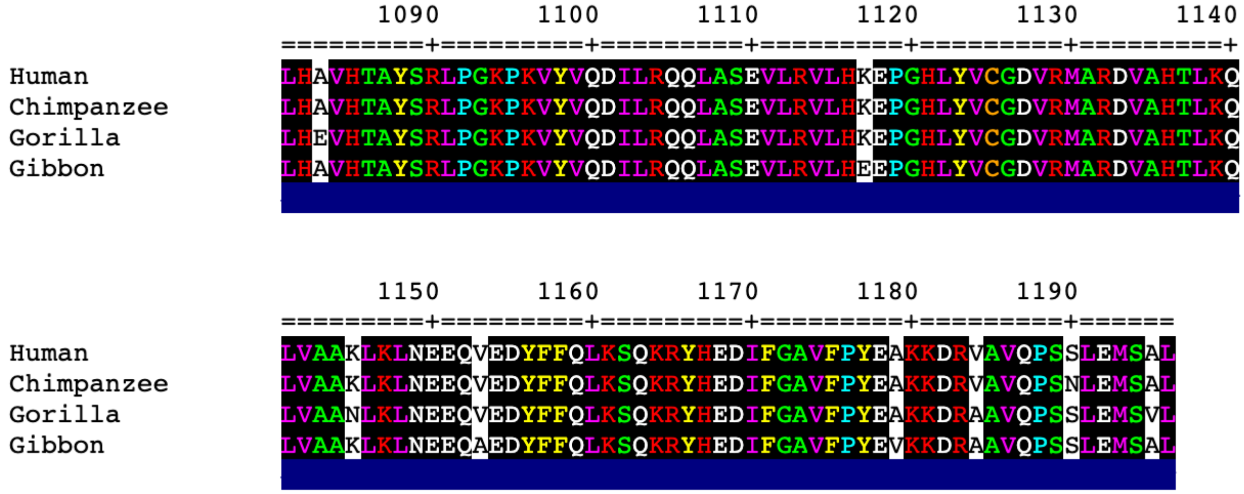


**S6 Fig**: Protein alignment of the NOS2 orthologs. Sites retained in the final alignment are underlined by the blue blocks.

**S1 Table**: Lower than expected substitution rate on the gibbon branch

| Gene | Gibbon % subs per site | Gibbon norm branch length | Human subs | Chimp subs | Gorilla subs | #1# subs | Gibbon subs | Align overlap | Align Sat | Higher in |
|---|---|---|---|---|---|---|---|---|---|---|
| ASIC1 | 0.00 | 0.04 | 0 | 20 | 0 | 0 | 0 | 307 | 64.09 | Chimpanzee |
| CALU | 0.00 | 0.04 | 0 | 20 | 0 | 0 | 0 | 320 | 99.07 | Chimpanzee |
| ZDHHC3 | 0.00 | 0.05 | 0 | 0 | 12 | 1 | 0 | 274 | 82.28 | Gorilla |
| POLR2A | 0.00 | 0.06 | 12 | 0 | 1 | 0 | 0 | 1904 | 97.14 | |
| SNAP25 | 0.00 | 0.07 | 0 | 0 | 9 | 0 | 0 | 206 | 100.00 | Gorilla |
| HM13 | 0.00 | 0.07 | 0 | 9 | 0 | 0 | 0 | 363 | 85.21 | Chimpanzee |
| SRF | 0.00 | 0.07 | 0 | 1 | 8 | 0 | 0 | 358 | 75.69 | Gorilla |
| VPS45 | 0.00 | 0.08 | 6 | 0 | 2 | 0 | 0 | 550 | 95.16 | |
| NELL2 | 0.00 | 0.08 | 1 | 3 | 4 | 0 | 0 | 826 | 98.45 | |
| DACT3 | 0.00 | 0.08 | 7 | 0 | 0 | 0 | 0 | 180 | 99.45 | Human |
| ITPKA | 0.00 | 0.08 | 0 | 0 | 7 | 0 | 0 | 312 | 75.54 | Gorilla |
| PGM1 | 0.00 | 0.08 | 7 | 0 | 0 | 0 | 0 | 474 | 81.72 | Human |
| GRIA4 | 0.00 | 0.08 | 1 | 5 | 1 | 0 | 0 | 897 | 99.45 | |
| GABRD | 0.00 | 0.09 | 0 | 0 | 6 | 0 | 0 | 319 | 70.42 | Gorilla |
| NEURL1 | 0.00 | 0.09 | 0 | 6 | 0 | 0 | 0 | 446 | 82.14 | Chimpanzee |
| CAAP1 | 0.00 | 0.09 | 0 | 2 | 4 | 0 | 0 | 342 | 98.84 | |
| BLVRB | 0.00 | 0.10 | 4 | 0 | 1 | 0 | 0 | 180 | 87.38 | Human |
| PAX5 | 0.00 | 0.10 | 0 | 0 | 5 | 0 | 0 | 364 | 93.09 | |
| LANCL3 | 0.00 | 0.10 | 0 | 0 | 5 | 0 | 0 | 400 | 97.09 | |
| HERC2 | 0.00 | 0.10 | 0 | 0 | 5 | 0 | 0 | 411 | 78.59 | |
| APBB1 | 0.00 | 0.10 | 0 | 5 | 0 | 0 | 0 | 495 | 86.24 | |
| KDM2A | 0.00 | 0.10 | 0 | 0 | 5 | 0 | 0 | 664 | 91.84 | |
| ZC3H7A | 0.10 | 0.11 | 2 | 2 | 9 | 0 | 1 | 971 | 100.00 | |
| UPRT | 0.00 | 0.11 | 0 | 4 | 0 | 0 | 0 | 138 | 79.77 | Chimpanzee |
| PAICS | 0.24 | 0.11 | 3 | 2 | 7 | 0 | 1 | 421 | 98.14 | Gorilla |
| PDCL | 0.00 | 0.11 | 4 | 0 | 0 | 0 | 0 | 301 | 100.00 | |
| UNC93B1 | 0.00 | 0.11 | 0 | 0 | 4 | 0 | 0 | 343 | 84.28 | |
| SEPT7 | 0.00 | 0.11 | 0 | 0 | 4 | 0 | 0 | 346 | 90.10 | |
| DYRK2 | 0.00 | 0.11 | 0 | 0 | 4 | 0 | 0 | 577 | 96.65 | |
| ACAP2 | 0.00 | 0.11 | 0 | 0 | 4 | 0 | 0 | 732 | 97.21 | |
| BMP8B | 0.25 | 0.12 | 1 | 2 | 5 | 3 | 1 | 402 | 100.00 | |
| SLC9A3 | 0.17 | 0.12 | 6 | 1 | 3 | 1 | 1 | 593 | 78.54 | |
| AASS | 0.11 | 0.13 | 2 | 1 | 5 | 1 | 1 | 872 | 97.76 | |
| CACUL1 | 0.28 | 0.14 | 1 | 1 | 6 | 0 | 1 | 352 | 95.65 | Gorilla |
| GMPS | 0.20 | 0.14 | 0 | 7 | 1 | 0 | 1 | 506 | 86.20 | Chimpanzee |
| NDUFAF6 | 0.30 | 0.14 | 0 | 1 | 2 | 5 | 1 | 333 | 100.00 | #1# |
| ZDHHC8 | 0.15 | 0.14 | 0 | 7 | 1 | 0 | 1 | 684 | 89.41 | |
| POU2F3 | 0.23 | 0.14 | 4 | 2 | 2 | 0 | 1 | 427 | 97.94 | |
| METTL11B | 0.37 | 0.15 | 1 | 1 | 5 | 0 | 1 | 272 | 97.84 | Gorilla |
| GFPT2 | 0.15 | 0.15 | 0 | 1 | 5 | 1 | 1 | 654 | 98.20 | |
| SIM1 | 0.13 | 0.15 | 4 | 3 | 0 | 0 | 1 | 765 | 99.87 | |
| AMPD3 | 0.13 | 0.15 | 3 | 0 | 4 | 0 | 1 | 774 | 99.74 | |
| GRIA1 | 0.11 | 0.15 | 1 | 1 | 5 | 0 | 1 | 912 | 99.56 | |
| BIN2 | 0.38 | 0.16 | 6 | 2 | 3 | 1 | 2 | 532 | 98.15 | |
| COPB2 | 0.22 | 0.17 | 1 | 0 | 10 | 0 | 2 | 889 | 99.44 | |
| TATDN1 | 0.35 | 0.17 | 1 | 1 | 4 | 0 | 1 | 284 | 95.62 | |
| NFATC1 | 0.15 | 0.17 | 1 | 5 | 0 | 0 | 1 | 665 | 77.96 | |
| NUDT9 | 0.58 | 0.17 | 3 | 3 | 4 | 1 | 2 | 345 | 98.85 | |
| TBX18 | 0.18 | 0.17 | 0 | 1 | 4 | 1 | 1 | 550 | 95.16 | |
| MYEF2 | 0.17 | 0.17 | 4 | 1 | 1 | 0 | 1 | 600 | 100.00 | |
| SLC25A24 | 0.46 | 0.18 | 0 | 0 | 10 | 0 | 2 | 436 | 95.20 | Gorilla |
| GLDC | 0.21 | 0.18 | 2 | 2 | 4 | 2 | 2 | 935 | 100.00 | |
| SPTBN4 | 0.08 | 0.18 | 2 | 4 | 3 | 1 | 2 | 2389 | 96.56 | |
| PCBP3 | 0.38 | 0.18 | 0 | 5 | 0 | 0 | 1 | 260 | 78.31 | Chimpanzee |
| PAX6 | 0.28 | 0.18 | 0 | 5 | 0 | 0 | 1 | 354 | 81.19 | |
| DNAJC9 | 0.41 | 0.18 | 0 | 4 | 1 | 0 | 1 | 241 | 97.57 | Chimpanzee |
| ESRRA | 0.24 | 0.18 | 0 | 0 | 5 | 0 | 1 | 422 | 100.00 | |
| SCAMP1 | 0.34 | 0.18 | 1 | 4 | 0 | 0 | 1 | 290 | 92.65 | |
| SYT12 | 0.24 | 0.18 | 1 | 0 | 4 | 0 | 1 | 420 | 100.00 | |
| CLIP2 | 0.30 | 0.18 | 2 | 2 | 9 | 1 | 3 | 1010 | 96.56 | |
| LONP1 | 0.15 | 0.18 | 0 | 1 | 4 | 0 | 1 | 664 | 91.97 | |
| XPO4 | 0.09 | 0.18 | 0 | 5 | 0 | 0 | 1 | 1078 | 94.48 | |
| ADAM23 | 0.12 | 0.18 | 0 | 0 | 4 | 1 | 1 | 832 | 100.00 | |
| TBXA2R | 0.61 | 0.19 | 1 | 0 | 8 | 0 | 2 | 328 | 95.63 | Gorilla |
| NACC2 | 0.52 | 0.19 | 1 | 2 | 6 | 0 | 2 | 382 | 78.60 | |
| NMBR | 0.51 | 0.19 | 2 | 3 | 4 | 0 | 2 | 390 | 100.00 | |
| PLCD3 | 0.29 | 0.19 | 1 | 5 | 3 | 0 | 2 | 682 | 88.57 | |
| ZNF707 | 0.55 | 0.19 | 2 | 4 | 2 | 1 | 2 | 366 | 98.65 | |
| ITPR1 | 0.08 | 0.19 | 8 | 0 | 1 | 0 | 2 | 2652 | 98.22 | |
| RCOR3 | 0.20 | 0.20 | 0 | 0 | 4 | 0 | 1 | 508 | 92.87 | |
| CCT6B | 0.58 | 0.20 | 1 | 3 | 6 | 2 | 3 | 515 | 97.17 | |
| PTCHD4 | 0.24 | 0.20 | 4 | 0 | 4 | 0 | 2 | 846 | 100.00 | |
| RBMX2 | 0.63 | 0.20 | 1 | 1 | 4 | 2 | 2 | 320 | 99.38 | |

| | | | | | | | | | | |
|---|---|---|---|---|---|---|---|---|---|---|
| INSC | 0.35 | 0.20 | 4 | 1 | 3 | 0 | 2 | 569 | 100.00 | |
| ADAMTS19 | 0.19 | 0.20 | 5 | 0 | 2 | 1 | 2 | 1059 | 99.53 | |
| NUP107 | 0.32 | 0.20 | 5 | 5 | 1 | 1 | 3 | 925 | 100.00 | |
| TMEM245 | 0.34 | 0.21 | 4 | 2 | 5 | 0 | 3 | 872 | 99.32 | |
| MEX3A | 0.47 | 0.21 | 0 | 0 | 7 | 0 | 2 | 424 | 94.64 | Gorilla |
| MEX3B | 0.36 | 0.21 | 1 | 1 | 5 | 0 | 2 | 549 | 96.49 | |
| ZBTB24 | 0.29 | 0.21 | 1 | 1 | 5 | 0 | 2 | 695 | 99.86 | |
| MCM4 | 0.24 | 0.21 | 2 | 0 | 5 | 0 | 2 | 851 | 98.84 | |
| TMEM43 | 0.50 | 0.21 | 1 | 2 | 4 | 0 | 2 | 400 | 100.00 | |
| GPAM | 0.24 | 0.21 | 3 | 0 | 4 | 0 | 2 | 828 | 100.00 | |
| TAX1BP1 | 0.51 | 0.22 | 3 | 4 | 6 | 1 | 4 | 778 | 99.87 | |
| TLN2 | 0.14 | 0.22 | 2 | 6 | 2 | 0 | 3 | 2125 | 92.84 | |
| GALNT14 | 0.54 | 0.22 | 2 | 4 | 3 | 1 | 3 | 554 | 100.00 | |
| CDH24 | 0.50 | 0.23 | 6 | 5 | 2 | 0 | 4 | 793 | 96.83 | |
| ULK1 | 0.77 | 0.23 | 2 | 12 | 2 | 0 | 5 | 646 | 76.00 | Chimpanzee |
| ADA | 0.57 | 0.23 | 5 | 1 | 0 | 0 | 2 | 352 | 96.97 | |
| NIM1K | 0.46 | 0.23 | 0 | 1 | 5 | 0 | 2 | 436 | 100.00 | |
| NUMBL | 0.35 | 0.23 | 0 | 0 | 5 | 1 | 2 | 575 | 95.67 | |
| PDSS2 | 0.50 | 0.23 | 4 | 0 | 2 | 0 | 2 | 399 | 100.00 | |
| ATP6AP1 | 0.47 | 0.23 | 0 | 2 | 4 | 0 | 2 | 422 | 94.62 | |
| ALDH1A2 | 0.40 | 0.23 | 2 | 0 | 4 | 0 | 2 | 500 | 96.53 | |
| MGAT4B | 0.39 | 0.23 | 0 | 1 | 1 | 4 | 2 | 507 | 92.52 | |
| GALNT3 | 0.32 | 0.23 | 1 | 4 | 1 | 0 | 2 | 633 | 100.00 | |
| NR2C1 | 0.51 | 0.24 | 3 | 5 | 1 | 0 | 3 | 592 | 98.18 | |
| PRR12 | 0.21 | 0.24 | 2 | 3 | 4 | 0 | 3 | 1435 | 99.03 | |
| TTC17 | 0.26 | 0.24 | 1 | 3 | 4 | 1 | 3 | 1141 | 100.00 | |
| PCDH8 | 0.32 | 0.24 | 2 | 4 | 2 | 1 | 3 | 928 | 95.38 | |
| SEC23IP | 0.40 | 0.24 | 6 | 4 | 2 | 0 | 4 | 992 | 99.20 | |
| NUP205 | 0.20 | 0.24 | 2 | 5 | 5 | 0 | 4 | 1998 | 99.30 | |
| SIPA1L2 | 0.23 | 0.24 | 4 | 3 | 5 | 0 | 4 | 1722 | 100.00 | |
| TP53BP2 | 0.44 | 0.24 | 5 | 6 | 3 | 1 | 5 | 1125 | 99.21 | |
| FBXL4 | 0.81 | 0.24 | 4 | 5 | 4 | 2 | 5 | 621 | 100.00 | |
| CHPF | 0.39 | 0.25 | 6 | 1 | 1 | 0 | 3 | 763 | 99.22 | |
| LACTB | 0.55 | 0.25 | 3 | 0 | 5 | 0 | 3 | 547 | 100.00 | |
| TRAF5 | 0.54 | 0.25 | 5 | 0 | 3 | 0 | 3 | 557 | 100.00 | |
| FLNB | 0.16 | 0.25 | 5 | 3 | 3 | 0 | 4 | 2541 | 98.30 | |
| AMOTL1 | 0.32 | 0.25 | 1 | 3 | 4 | 0 | 3 | 938 | 98.12 | |
| CAMSAP2 | 0.27 | 0.25 | 2 | 6 | 1 | 2 | 4 | 1464 | 98.32 | |
| CIC | 0.21 | 0.25 | 5 | 2 | 6 | 1 | 5 | 2353 | 98.66 | |
| BAG6 | 0.37 | 0.25 | 4 | 2 | 5 | 0 | 4 | 1073 | 95.89 | |
| SLC26A7 | 0.47 | 0.25 | 2 | 2 | 4 | 0 | 3 | 635 | 97.54 | |
| FAM135A | 0.54 | 0.25 | 4 | 7 | 10 | 2 | 8 | 1470 | 97.03 | |
| TPX2 | 0.63 | 0.25 | 3 | 3 | 5 | 0 | 4 | 634 | 90.96 | |
| CRBN | 0.62 | 0.25 | 4 | 2 | 2 | 0 | 3 | 487 | 99.39 | |
| GNL2 | 0.41 | 0.25 | 1 | 2 | 4 | 1 | 3 | 730 | 100.00 | |
| MANEA | 0.65 | 0.25 | 2 | 4 | 2 | 0 | 3 | 462 | 100.00 | |
| GATAD2A | 0.47 | 0.25 | 1 | 1 | 4 | 2 | 3 | 634 | 100.00 | |
| MAP9 | 0.65 | 0.25 | 1 | 2 | 4 | 1 | 3 | 464 | 100.00 | |
| NID1 | 0.56 | 0.26 | 6 | 5 | 5 | 3 | 7 | 1247 | 100.00 | |
| CDC42BPB | 0.27 | 0.26 | 2 | 5 | 3 | 0 | 4 | 1462 | 91.78 | |
| AFDN | 0.23 | 0.26 | 1 | 4 | 4 | 1 | 4 | 1709 | 95.32 | |
| EML6 | 0.22 | 0.26 | 3 | 4 | 2 | 1 | 4 | 1858 | 97.79 | |
| SLC26A4 | 0.51 | 0.26 | 5 | 2 | 3 | 0 | 4 | 780 | 100.00 | |
| CLSTN2 | 0.46 | 0.26 | 4 | 4 | 2 | 0 | 4 | 868 | 95.91 | |
| ELOA | 0.54 | 0.26 | 4 | 3 | 3 | 0 | 4 | 747 | 100.00 | |
| NDST4 | 0.46 | 0.26 | 2 | 4 | 2 | 2 | 4 | 872 | 100.00 | |
| SORBS1 | 0.46 | 0.27 | 6 | 3 | 5 | 1 | 6 | 1293 | 100.00 | |
| SLC4A2 | 0.47 | 0.27 | 1 | 2 | 9 | 0 | 5 | 1055 | 95.39 | |
| DDX18 | 0.75 | 0.27 | 7 | 3 | 1 | 1 | 5 | 670 | 100.00 | |
| LGR6 | 0.89 | 0.27 | 7 | 6 | 6 | 1 | 8 | 896 | 100.00 | |
| GRIP2 | 0.88 | 0.28 | 12 | 6 | 7 | 0 | 10 | 1132 | 99.30 | Human |
| RET | 0.82 | 0.28 | 12 | 7 | 3 | 0 | 9 | 1097 | 98.47 | Human |
| MYOM1 | 0.60 | 0.28 | 13 | 3 | 2 | 1 | 8 | 1333 | 97.16 | Human |
| DPP6 | 0.64 | 0.29 | 2 | 5 | 4 | 0 | 5 | 786 | 97.88 | |
| IQSEC1 | 0.47 | 0.29 | 4 | 2 | 2 | 3 | 5 | 1066 | 97.09 | |
| IGSF22 | 0.55 | 0.30 | 10 | 1 | 4 | 0 | 7 | 1281 | 98.69 | |
| LRIG2 | 0.81 | 0.30 | 3 | 8 | 3 | 1 | 7 | 859 | 87.03 | |
| KIF16B | 0.65 | 0.30 | 4 | 3 | 8 | 4 | 9 | 1392 | 100.00 | |
| CEP128 | 0.92 | 0.31 | 6 | 8 | 6 | 1 | 10 | 1091 | 99.73 | |
| LUZP1 | 0.93 | 0.31 | 7 | 4 | 6 | 3 | 10 | 1076 | 100.00 | |
| TMC3 | 1.00 | 0.32 | 7 | 7 | 7 | 1 | 11 | 1100 | 100.00 | |
| PCDH15 | 0.92 | 0.32 | 8 | 11 | 13 | 2 | 17 | 1847 | 98.72 | |
| HERC1 | 0.23 | 0.32 | 7 | 6 | 7 | 1 | 11 | 4854 | 100.00 | |
| LCT | 1.19 | 0.35 | 11 | 11 | 10 | 9 | 23 | 1927 | 100.00 | |

**S2 Table**: Lower than expected substitution rate on the human branch

| Gene | Human % subs per site | Human norm branch length | Human subs | Chimp subs | Gorilla subs | #1# subs | Gibbon subs | Align overlap | Align Sat | Higher in |
|---|---|---|---|---|---|---|---|---|---|---|
| CEP295NL | 0.00 | 0.01 | 0 | 5 | 9 | 7 | 41 | 589 | 98.66 | Gibbon |
| PODXL | 0.00 | 0.02 | 0 | 4 | 7 | 1 | 39 | 478 | 85.97 | Gibbon |
| C2orf81 | 0.00 | 0.02 | 0 | 4 | 9 | 2 | 33 | 537 | 99.63 | Gibbon |
| HSF5 | 0.00 | 0.02 | 0 | 3 | 7 | 2 | 32 | 593 | 99.50 | Gibbon |
| PTCD3 | 0.00 | 0.02 | 0 | 5 | 6 | 2 | 29 | 686 | 99.56 | Gibbon |
| MUC1 | 0.00 | 0.02 | 0 | 3 | 6 | 0 | 29 | 470 | 97.31 | Gibbon |
| LEKR1 | 0.00 | 0.02 | 0 | 1 | 6 | 0 | 31 | 692 | 100.00 | Gibbon |
| RIPK1 | 0.00 | 0.02 | 0 | 4 | 2 | 2 | 30 | 671 | 100.00 | Gibbon |
| NOS2 | 0.00 | 0.03 | 0 | 4 | 8 | 1 | 21 | 1153 | 100.00 | |
| PER2 | 0.00 | 0.03 | 0 | 3 | 4 | 3 | 22 | 1255 | 100.00 | |
| SEL1L2 | 0.00 | 0.03 | 0 | 5 | 4 | 1 | 21 | 688 | 100.00 | Gibbon |
| RFC1 | 0.00 | 0.03 | 0 | 3 | 4 | 2 | 22 | 1102 | 97.52 | |
| USP54 | 0.00 | 0.03 | 0 | 7 | 6 | 1 | 17 | 1437 | 100.00 | |
| CD4 | 0.00 | 0.03 | 0 | 4 | 7 | 3 | 15 | 458 | 100.00 | |
| CEP68 | 0.00 | 0.03 | 0 | 2 | 10 | 2 | 15 | 573 | 98.79 | Gorilla |
| CAGE1 | 0.12 | 0.03 | 1 | 6 | 13 | 0 | 42 | 839 | 100.00 | Gibbon |
| IFT81 | 0.00 | 0.03 | 0 | 1 | 4 | 0 | 23 | 676 | 100.00 | Gibbon |
| CCPG1 | 0.00 | 0.03 | 0 | 3 | 4 | 1 | 20 | 786 | 97.64 | |
| GCM2 | 0.00 | 0.03 | 0 | 5 | 7 | 3 | 13 | 506 | 100.00 | |
| TEX11 | 0.11 | 0.03 | 1 | 10 | 11 | 2 | 36 | 906 | 99.34 | Gibbon |
| CHFR | 0.00 | 0.03 | 0 | 2 | 5 | 1 | 19 | 543 | 85.92 | Gibbon |
| C10orf67 | 0.00 | 0.03 | 0 | 4 | 2 | 4 | 17 | 547 | 99.64 | |
| ITGA10 | 0.00 | 0.03 | 0 | 5 | 4 | 0 | 18 | 1167 | 100.00 | |
| C3orf20 | 0.12 | 0.03 | 1 | 3 | 11 | 3 | 39 | 853 | 99.53 | Gibbon |
| ATF7IP2 | 0.00 | 0.03 | 0 | 3 | 4 | 0 | 19 | 671 | 99.70 | Gibbon |
| SYNPO2L | 0.00 | 0.03 | 0 | 3 | 5 | 0 | 18 | 976 | 99.90 | |
| IARS | 0.00 | 0.03 | 0 | 3 | 8 | 1 | 14 | 1136 | 96.35 | |
| RSL1D1 | 0.00 | 0.03 | 0 | 2 | 5 | 1 | 17 | 490 | 100.00 | |
| VARS2 | 0.00 | 0.03 | 0 | 4 | 3 | 0 | 18 | 1010 | 94.75 | |
| FAM171A1 | 0.00 | 0.03 | 0 | 2 | 10 | 0 | 13 | 851 | 97.15 | |
| ZNF536 | 0.00 | 0.03 | 0 | 1 | 7 | 2 | 15 | 1299 | 99.92 | |
| MAPKBP1 | 0.07 | 0.03 | 1 | 7 | 20 | 2 | 24 | 1497 | 98.88 | Gorilla |
| KIAA1257 | 0.00 | 0.03 | 0 | 3 | 6 | 1 | 14 | 558 | 97.55 | |
| MBD4 | 0.00 | 0.03 | 0 | 3 | 7 | 1 | 13 | 578 | 100.00 | |
| DEPDC1 | 0.00 | 0.03 | 0 | 4 | 4 | 1 | 15 | 722 | 100.00 | |
| PLA2G6 | 0.00 | 0.03 | 0 | 3 | 8 | 0 | 13 | 751 | 93.29 | |
| SWT1 | 0.00 | 0.04 | 0 | 2 | 5 | 1 | 15 | 521 | 94.90 | |
| NOX3 | 0.00 | 0.04 | 0 | 4 | 8 | 0 | 11 | 568 | 100.00 | Gorilla |
| ATG9B | 0.00 | 0.04 | 0 | 2 | 6 | 2 | 13 | 757 | 96.80 | |
| USP38 | 0.00 | 0.04 | 0 | 4 | 3 | 0 | 16 | 977 | 100.00 | |
| SMYD4 | 0.00 | 0.04 | 0 | 4 | 3 | 4 | 12 | 683 | 91.80 | |
| KIF9 | 0.00 | 0.04 | 0 | 4 | 7 | 0 | 12 | 790 | 100.00 | |
| CENPC | 0.11 | 0.04 | 1 | 5 | 7 | 1 | 35 | 943 | 100.00 | Gibbon |
| CCDC77 | 0.00 | 0.04 | 0 | 3 | 4 | 0 | 15 | 474 | 99.79 | |
| SERGEF | 0.00 | 0.04 | 0 | 5 | 2 | 1 | 14 | 454 | 99.34 | |
| TRMT61B | 0.00 | 0.04 | 0 | 4 | 3 | 1 | 14 | 464 | 97.68 | |
| SPINT1 | 0.00 | 0.04 | 0 | 1 | 6 | 2 | 13 | 523 | 98.87 | |
| TREH | 0.00 | 0.04 | 0 | 3 | 6 | 0 | 13 | 582 | 99.83 | |
| TRPM6 | 0.05 | 0.04 | 1 | 4 | 7 | 1 | 36 | 2007 | 99.50 | |
| SETMAR | 0.00 | 0.04 | 0 | 3 | 5 | 1 | 13 | 684 | 100.00 | |
| USP43 | 0.00 | 0.04 | 0 | 4 | 3 | 0 | 15 | 1103 | 100.00 | |
| PIWIL4 | 0.00 | 0.04 | 0 | 5 | 7 | 0 | 10 | 806 | 99.38 | |
| PELP1 | 0.00 | 0.04 | 0 | 4 | 2 | 1 | 15 | 1159 | 98.30 | |
| PNLDC1 | 0.00 | 0.04 | 0 | 5 | 2 | 2 | 12 | 505 | 97.12 | |
| RPS6KL1 | 0.00 | 0.04 | 0 | 2 | 8 | 2 | 9 | 549 | 100.00 | Gorilla |
| DNTTIP2 | 0.00 | 0.04 | 0 | 2 | 7 | 0 | 12 | 753 | 99.87 | |
| UTP14A | 0.00 | 0.04 | 0 | 5 | 4 | 0 | 12 | 754 | 98.69 | |
| INPP5D | 0.00 | 0.04 | 0 | 3 | 4 | 0 | 14 | 1027 | 96.34 | |
| PHKB | 0.00 | 0.04 | 0 | 10 | 2 | 0 | 9 | 1084 | 99.18 | |
| ZNF16 | 0.00 | 0.04 | 0 | 7 | 1 | 1 | 11 | 627 | 94.86 | |
| CCDC80 | 0.00 | 0.04 | 0 | 2 | 5 | 0 | 13 | 921 | 98.82 | |
| MAPK8IP3 | 0.00 | 0.04 | 0 | 6 | 3 | 0 | 11 | 1189 | 90.49 | |
| CKAP5 | 0.00 | 0.04 | 0 | 3 | 4 | 0 | 13 | 1989 | 97.88 | |
| SMC1B | 0.08 | 0.04 | 1 | 6 | 4 | 0 | 33 | 1197 | 98.76 | Gibbon |
| CTTN | 0.00 | 0.04 | 0 | 1 | 4 | 1 | 13 | 512 | 93.26 | |
| SELENOO | 0.00 | 0.04 | 0 | 2 | 5 | 0 | 12 | 509 | 100.00 | |
| CYP17A1 | 0.00 | 0.04 | 0 | 1 | 6 | 1 | 11 | 503 | 99.41 | |
| SSC4D | 0.00 | 0.04 | 0 | 4 | 5 | 0 | 10 | 469 | 100.00 | |
| RXFP1 | 0.00 | 0.04 | 0 | 1 | 9 | 0 | 9 | 726 | 96.16 | |
| THNSL1 | 0.00 | 0.04 | 0 | 1 | 5 | 2 | 11 | 743 | 100.00 | |
| IQCA1 | 0.00 | 0.04 | 0 | 4 | 1 | 2 | 12 | 830 | 100.00 | |
| ANO6 | 0.00 | 0.04 | 0 | 1 | 11 | 0 | 7 | 901 | 97.83 | Gorilla |
| SEMA7A | 0.00 | 0.04 | 0 | 4 | 6 | 2 | 7 | 664 | 100.00 | |

| | | | | | | | | | | |
|---|---|---|---|---|---|---|---|---|---|---|
| NOL6 | 0.09 | 0.04 | 1 | 3 | 5 | 2 | 30 | 1111 | 96.95 | Gibbon |
| FAM124A | 0.00 | 0.04 | 0 | 2 | 4 | 0 | 12 | 582 | 100.00 | |
| TFR2 | 0.00 | 0.04 | 0 | 4 | 1 | 1 | 12 | 610 | 87.39 | |
| B4GALNT4 | 0.00 | 0.04 | 0 | 2 | 4 | 0 | 12 | 627 | 71.41 | |
| METTL25 | 0.00 | 0.04 | 0 | 3 | 4 | 1 | 10 | 603 | 100.00 | |
| BACH1 | 0.00 | 0.04 | 0 | 2 | 5 | 1 | 10 | 736 | 100.00 | |
| LBR | 0.00 | 0.04 | 0 | 6 | 4 | 1 | 7 | 615 | 100.00 | |
| JMY | 0.00 | 0.04 | 0 | 6 | 2 | 0 | 10 | 930 | 100.00 | |
| NCOR1 | 0.00 | 0.04 | 0 | 3 | 5 | 0 | 10 | 2360 | 98.91 | |
| OTOL1 | 0.00 | 0.05 | 0 | 1 | 6 | 0 | 10 | 463 | 98.51 | |
| ZWILCH | 0.00 | 0.05 | 0 | 1 | 7 | 0 | 9 | 513 | 92.93 | Gorilla |
| P3H3 | 0.00 | 0.05 | 0 | 2 | 4 | 0 | 11 | 736 | 100.00 | |
| SPICE1 | 0.00 | 0.05 | 0 | 5 | 3 | 0 | 9 | 673 | 93.47 | |
| DCLK3 | 0.00 | 0.05 | 0 | 2 | 4 | 0 | 11 | 817 | 100.00 | |
| STAT6 | 0.00 | 0.05 | 0 | 1 | 7 | 0 | 9 | 839 | 99.64 | |
| WDR63 | 0.00 | 0.05 | 0 | 5 | 1 | 1 | 10 | 836 | 97.55 | |
| GAS2L3 | 0.00 | 0.05 | 0 | 2 | 5 | 2 | 8 | 694 | 100.00 | |
| BMP2K | 0.00 | 0.05 | 0 | 1 | 4 | 0 | 12 | 1138 | 99.56 | |
| LOXL4 | 0.00 | 0.05 | 0 | 4 | 5 | 1 | 7 | 756 | 100.00 | |
| NEFH | 0.12 | 0.05 | 1 | 4 | 4 | 3 | 25 | 834 | 96.08 | Gibbon |
| TTLL13P | 0.13 | 0.05 | 1 | 5 | 6 | 3 | 22 | 775 | 97.12 | |
| BRIP1 | 0.08 | 0.05 | 1 | 3 | 6 | 0 | 27 | 1215 | 98.86 | |
| NUP210L | 0.11 | 0.05 | 2 | 7 | 6 | 3 | 40 | 1777 | 96.31 | |
| NARFL | 0.00 | 0.05 | 0 | 2 | 4 | 0 | 10 | 464 | 97.68 | |
| MIEF2 | 0.00 | 0.05 | 0 | 2 | 4 | 1 | 9 | 461 | 99.35 | |
| CAPN15 | 0.00 | 0.05 | 0 | 1 | 8 | 0 | 7 | 565 | 80.48 | Gorilla |
| ZSCAN22 | 0.00 | 0.05 | 0 | 4 | 4 | 0 | 8 | 489 | 99.59 | |
| ABHD15 | 0.00 | 0.05 | 0 | 4 | 5 | 0 | 7 | 468 | 100.00 | |
| ZBTB47 | 0.00 | 0.05 | 0 | 4 | 2 | 0 | 10 | 648 | 98.93 | |
| VPS9D1 | 0.00 | 0.05 | 0 | 4 | 3 | 0 | 9 | 631 | 100.00 | |
| CRIM1 | 0.00 | 0.05 | 0 | 3 | 4 | 1 | 8 | 925 | 100.00 | |
| CCDC39 | 0.12 | 0.05 | 1 | 4 | 5 | 0 | 26 | 867 | 100.00 | Gibbon |
| MPHOSPH9 | 0.09 | 0.05 | 1 | 6 | 7 | 0 | 22 | 1147 | 100.00 | |
| CEP350 | 0.13 | 0.05 | 4 | 12 | 18 | 8 | 55 | 3063 | 98.97 | |
| JCAD | 0.15 | 0.05 | 2 | 10 | 8 | 2 | 34 | 1332 | 98.38 | Gibbon |
| OTOF | 0.10 | 0.05 | 2 | 8 | 13 | 2 | 31 | 1962 | 99.70 | |
| CC2D2B | 0.19 | 0.05 | 2 | 6 | 5 | 6 | 36 | 1043 | 98.58 | Gibbon |
| CIZ1 | 0.12 | 0.05 | 1 | 3 | 9 | 4 | 18 | 860 | 95.56 | |
| CDKL3 | 0.00 | 0.05 | 0 | 1 | 4 | 1 | 9 | 537 | 100.00 | |
| GNL3L | 0.00 | 0.05 | 0 | 2 | 5 | 0 | 8 | 575 | 98.80 | |
| AMHR2 | 0.00 | 0.05 | 0 | 2 | 4 | 1 | 8 | 573 | 100.00 | |
| ASIC5 | 0.00 | 0.05 | 0 | 2 | 4 | 2 | 7 | 505 | 100.00 | |
| TMCO3 | 0.00 | 0.05 | 0 | 5 | 2 | 1 | 7 | 564 | 84.30 | |
| BCLAF3 | 0.00 | 0.05 | 0 | 1 | 6 | 1 | 7 | 682 | 100.00 | |
| RNF216 | 0.00 | 0.05 | 0 | 2 | 6 | 2 | 5 | 705 | 82.65 | |
| MAP3K15 | 0.00 | 0.05 | 0 | 7 | 2 | 0 | 6 | 943 | 82.43 | |
| STARD13 | 0.00 | 0.05 | 0 | 1 | 6 | 1 | 7 | 930 | 100.00 | |
| ZNF512B | 0.00 | 0.05 | 0 | 3 | 5 | 1 | 6 | 806 | 94.38 | |
| KIDINS220 | 0.00 | 0.05 | 0 | 2 | 6 | 1 | 6 | 1706 | 98.44 | |
| NCOA3 | 0.07 | 0.05 | 1 | 5 | 3 | 2 | 23 | 1415 | 99.72 | |
| AASDH | 0.09 | 0.05 | 1 | 4 | 10 | 1 | 18 | 1098 | 100.00 | |
| NUP160 | 0.07 | 0.05 | 1 | 8 | 6 | 5 | 14 | 1436 | 100.00 | |
| CSPG4 | 0.20 | 0.05 | 4 | 13 | 21 | 7 | 47 | 2046 | 94.29 | |
| ST14 | 0.12 | 0.05 | 1 | 2 | 6 | 1 | 23 | 852 | 99.65 | Gibbon |
| SERINC4 | 0.00 | 0.05 | 0 | 1 | 4 | 0 | 9 | 487 | 94.02 | |
| TNS2 | 0.07 | 0.05 | 1 | 4 | 4 | 0 | 24 | 1361 | 100.00 | |
| BOC | 0.09 | 0.05 | 1 | 7 | 5 | 1 | 19 | 1111 | 99.64 | |
| TXLNG | 0.00 | 0.05 | 0 | 6 | 1 | 0 | 7 | 514 | 98.09 | |
| NCAPD3 | 0.13 | 0.05 | 2 | 10 | 15 | 3 | 22 | 1492 | 99.60 | |
| GIGYF2 | 0.00 | 0.05 | 0 | 1 | 7 | 0 | 6 | 1217 | 94.41 | |
| CDC42BPG | 0.14 | 0.05 | 2 | 8 | 14 | 2 | 25 | 1442 | 98.03 | |
| PLCB2 | 0.09 | 0.05 | 1 | 2 | 8 | 1 | 20 | 1083 | 100.00 | |
| FUK | 0.10 | 0.05 | 1 | 6 | 6 | 1 | 18 | 1004 | 94.10 | |
| TNRC6C | 0.05 | 0.05 | 1 | 2 | 11 | 0 | 18 | 1865 | 97.75 | |
| TTLL4 | 0.20 | 0.05 | 2 | 3 | 13 | 3 | 29 | 1015 | 94.07 | |
| PLEKHG2 | 0.14 | 0.05 | 2 | 8 | 9 | 2 | 29 | 1383 | 99.78 | |
| INCENP | 0.11 | 0.06 | 1 | 2 | 7 | 2 | 19 | 903 | 100.00 | |
| TNS3 | 0.07 | 0.06 | 1 | 2 | 4 | 1 | 23 | 1359 | 96.93 | |
| CEP135 | 0.09 | 0.06 | 1 | 6 | 4 | 1 | 19 | 1140 | 100.00 | |
| SNAP47 | 0.00 | 0.06 | 0 | 2 | 4 | 1 | 6 | 463 | 99.78 | |
| LRRC45 | 0.00 | 0.06 | 0 | 1 | 5 | 0 | 7 | 659 | 99.10 | |
| PCNX1 | 0.04 | 0.06 | 1 | 7 | 4 | 1 | 18 | 2251 | 97.03 | |
| IGF2R | 0.12 | 0.06 | 3 | 7 | 7 | 4 | 45 | 2440 | 99.27 | |
| FGD5 | 0.16 | 0.06 | 2 | 3 | 7 | 2 | 34 | 1241 | 96.20 | Gibbon |
| SETX | 0.28 | 0.06 | 7 | 23 | 20 | 1 | 84 | 2511 | 97.48 | Gibbon |
| TDRD1 | 0.17 | 0.06 | 2 | 4 | 7 | 3 | 31 | 1149 | 99.65 | Gibbon |
| GPR179 | 0.30 | 0.06 | 7 | 11 | 21 | 3 | 89 | 2347 | 99.75 | Gibbon |
| EGFLAM | 0.11 | 0.06 | 1 | 2 | 5 | 0 | 21 | 911 | 94.90 | |

| | | | | | | | | | | |
|---|---|---|---|---|---|---|---|---|---|---|
| TTF2 | 0.17 | 0.06 | 2 | 6 | 6 | 4 | 28 | 1161 | 100.00 | |
| FHDC1 | 0.18 | 0.06 | 2 | 7 | 11 | 3 | 23 | 1120 | 99.82 | |
| CFAP61 | 0.16 | 0.06 | 2 | 6 | 15 | 1 | 22 | 1236 | 99.92 | Gorilla |
| GAK | 0.09 | 0.06 | 1 | 3 | 6 | 3 | 16 | 1072 | 87.87 | |
| ABCA12 | 0.08 | 0.06 | 2 | 11 | 10 | 1 | 22 | 2547 | 99.88 | |
| WDR64 | 0.19 | 0.06 | 2 | 8 | 7 | 4 | 24 | 1049 | 99.71 | |
| MST1R | 0.15 | 0.06 | 2 | 4 | 10 | 5 | 24 | 1312 | 100.00 | |
| MPDZ | 0.10 | 0.06 | 2 | 6 | 14 | 3 | 20 | 2048 | 98.27 | |
| MASTL | 0.11 | 0.06 | 1 | 5 | 4 | 1 | 17 | 877 | 99.89 | |
| HEPHL1 | 0.09 | 0.06 | 1 | 3 | 5 | 0 | 19 | 1153 | 100.00 | |
| POLG | 0.08 | 0.06 | 1 | 2 | 7 | 2 | 16 | 1225 | 99.59 | |
| ANKAR | 0.07 | 0.06 | 1 | 4 | 5 | 0 | 18 | 1434 | 100.00 | |
| UHRF1BP1 | 0.07 | 0.06 | 1 | 3 | 6 | 3 | 15 | 1440 | 100.00 | |
| CCDC18 | 0.21 | 0.06 | 3 | 6 | 11 | 2 | 38 | 1453 | 100.00 | Gibbon |
| AXDND1 | 0.20 | 0.06 | 2 | 6 | 5 | 4 | 26 | 1009 | 99.80 | |
| C5 | 0.13 | 0.06 | 2 | 5 | 6 | 2 | 28 | 1586 | 96.35 | |
| RIN1 | 0.13 | 0.06 | 1 | 5 | 5 | 2 | 14 | 782 | 100.00 | |
| MYOM2 | 0.10 | 0.06 | 1 | 3 | 7 | 1 | 15 | 1028 | 80.50 | |
| THADA | 0.23 | 0.06 | 4 | 8 | 9 | 1 | 52 | 1772 | 94.56 | Gibbon |
| TACC1 | 0.13 | 0.06 | 1 | 2 | 6 | 1 | 16 | 759 | 97.18 | |
| WDHD1 | 0.09 | 0.06 | 1 | 2 | 5 | 0 | 18 | 1068 | 97.09 | |
| RUBCN | 0.11 | 0.06 | 1 | 8 | 2 | 0 | 15 | 932 | 96.48 | |
| NLRX1 | 0.11 | 0.06 | 1 | 4 | 3 | 4 | 14 | 900 | 99.89 | |
| CNTRL | 0.17 | 0.06 | 4 | 8 | 15 | 5 | 40 | 2305 | 99.27 | |
| PTPRQ | 0.20 | 0.07 | 4 | 9 | 13 | 3 | 42 | 2011 | 93.75 | |
| AP5B1 | 0.12 | 0.07 | 1 | 3 | 4 | 0 | 17 | 839 | 96.22 | |
| MTR | 0.08 | 0.07 | 1 | 1 | 4 | 0 | 19 | 1232 | 98.56 | |
| SEC24A | 0.09 | 0.07 | 1 | 5 | 2 | 0 | 17 | 1093 | 100.00 | |
| LRIG3 | 0.09 | 0.07 | 1 | 3 | 5 | 1 | 15 | 1119 | 100.00 | |
| TRANK1 | 0.07 | 0.07 | 2 | 6 | 6 | 3 | 23 | 2886 | 98.67 | |
| CASP8AP2 | 0.21 | 0.07 | 4 | 8 | 8 | 5 | 44 | 1941 | 98.48 | |
| LRIG1 | 0.19 | 0.07 | 2 | 3 | 5 | 1 | 28 | 1073 | 98.53 | Gibbon |
| TTC21A | 0.16 | 0.07 | 2 | 4 | 9 | 1 | 23 | 1285 | 97.35 | |
| NPAT | 0.22 | 0.07 | 3 | 4 | 10 | 2 | 34 | 1371 | 97.86 | Gibbon |
| KNDC1 | 0.19 | 0.07 | 2 | 9 | 3 | 1 | 23 | 1038 | 73.46 | |
| GPR158 | 0.16 | 0.07 | 2 | 3 | 7 | 3 | 23 | 1214 | 100.00 | |
| COL12A1 | 0.07 | 0.07 | 2 | 4 | 4 | 3 | 25 | 3033 | 99.97 | |
| RP1 | 0.30 | 0.07 | 6 | 17 | 18 | 3 | 50 | 2026 | 99.22 | |
| CARMIL2 | 0.16 | 0.07 | 2 | 5 | 6 | 1 | 23 | 1265 | 93.29 | |
| RGS3 | 0.17 | 0.07 | 2 | 3 | 5 | 3 | 23 | 1160 | 97.81 | |
| IQGAP3 | 0.13 | 0.07 | 2 | 4 | 8 | 1 | 21 | 1496 | 94.15 | |
| TTC28 | 0.09 | 0.07 | 2 | 4 | 3 | 1 | 26 | 2116 | 92.20 | |
| BOD1L1 | 0.30 | 0.07 | 9 | 14 | 22 | 5 | 80 | 3048 | 100.00 | Gibbon |
| FAM135B | 0.14 | 0.07 | 2 | 5 | 5 | 0 | 23 | 1398 | 99.43 | |
| ZFYVE26 | 0.16 | 0.08 | 4 | 6 | 9 | 1 | 39 | 2471 | 98.33 | |
| MAP2 | 0.16 | 0.08 | 3 | 6 | 8 | 3 | 26 | 1827 | 100.00 | |
| PDZD2 | 0.32 | 0.08 | 8 | 15 | 16 | 1 | 68 | 2523 | 97.11 | Gibbon |
| ZBED9 | 0.23 | 0.08 | 3 | 4 | 5 | 2 | 31 | 1325 | 100.00 | |
| ATP10B | 0.21 | 0.08 | 3 | 3 | 11 | 3 | 24 | 1460 | 100.00 | |
| MAP1A | 0.26 | 0.08 | 8 | 10 | 12 | 9 | 65 | 3037 | 99.90 | |
| NOTCH4 | 0.21 | 0.08 | 4 | 5 | 7 | 3 | 36 | 1861 | 97.28 | |
| GOLGA4 | 0.24 | 0.08 | 5 | 15 | 9 | 2 | 36 | 2112 | 98.00 | |
| LAMA3 | 0.27 | 0.08 | 8 | 13 | 16 | 10 | 55 | 3005 | 95.34 | |
| VWF | 0.25 | 0.08 | 5 | 8 | 10 | 5 | 38 | 1974 | 98.06 | |
| REV3L | 0.17 | 0.08 | 5 | 6 | 16 | 4 | 35 | 2893 | 99.97 | |
| ZNF292 | 0.15 | 0.08 | 4 | 4 | 11 | 3 | 32 | 2653 | 98.48 | |
| DNAH1 | 0.30 | 0.09 | 12 | 15 | 35 | 10 | 67 | 4060 | 96.90 | |

**S3 Table**: Lower than expected substitution rate on the chimpanzee branch

| Gene | Chimp % subs per site | Chimp norm branch length | Human subs | Chimp subs | Gorilla subs | #1# subs | Gibbon subs | Align overlap | Align Sat | Higher in |
|---|---|---|---|---|---|---|---|---|---|---|
| CYLC1 | 0.00 | 0.02 | 1 | 0 | 7 | 5 | 35 | 630 | 97.83 | Gibbon |
| NSD1 | 0.00 | 0.02 | 8 | 0 | 5 | 1 | 29 | 2618 | 99.47 | |
| TROAP | 0.00 | 0.02 | 3 | 0 | 4 | 2 | 31 | 771 | 97.10 | Gibbon |
| SDK2 | 0.00 | 0.02 | 3 | 0 | 5 | 1 | 31 | 1893 | 94.65 | |
| LTN1 | 0.00 | 0.02 | 6 | 0 | 6 | 3 | 21 | 1763 | 100.00 | |
| C1orf87 | 0.00 | 0.03 | 2 | 0 | 5 | 0 | 27 | 546 | 100.00 | Gibbon |
| ADGRG7 | 0.00 | 0.03 | 2 | 0 | 3 | 4 | 25 | 797 | 100.00 | Gibbon |
| CD3EAP | 0.00 | 0.03 | 4 | 0 | 8 | 1 | 20 | 501 | 98.43 | |
| KCTD19 | 0.00 | 0.03 | 2 | 0 | 5 | 2 | 23 | 734 | 95.82 | Gibbon |
| FGD3 | 0.00 | 0.03 | 2 | 0 | 5 | 3 | 20 | 678 | 100.00 | Gibbon |
| CDKL2 | 0.00 | 0.03 | 3 | 0 | 7 | 2 | 17 | 505 | 100.00 | |
| BICDL2 | 0.00 | 0.03 | 5 | 0 | 10 | 2 | 12 | 507 | 100.00 | Gorilla |
| ADAR | 0.00 | 0.03 | 5 | 0 | 5 | 1 | 18 | 1171 | 99.24 | |
| SPAG1 | 0.00 | 0.03 | 2 | 0 | 8 | 1 | 17 | 847 | 97.13 | |
| PLEKHA6 | 0.00 | 0.03 | 2 | 0 | 5 | 2 | 19 | 1133 | 97.67 | |
| ARAP3 | 0.00 | 0.03 | 6 | 0 | 6 | 0 | 16 | 1490 | 97.32 | |
| BFSP1 | 0.00 | 0.03 | 1 | 0 | 5 | 2 | 19 | 650 | 99.54 | Gibbon |
| TCIRG1 | 0.00 | 0.03 | 2 | 0 | 5 | 5 | 15 | 830 | 100.00 | |
| PER1 | 0.00 | 0.03 | 3 | 0 | 4 | 0 | 20 | 1262 | 99.53 | |
| EFHC1 | 0.00 | 0.03 | 2 | 0 | 4 | 1 | 18 | 641 | 100.00 | Gibbon |
| SSH3 | 0.00 | 0.03 | 6 | 0 | 4 | 0 | 15 | 598 | 97.71 | |
| CC2D1B | 0.00 | 0.03 | 1 | 0 | 4 | 2 | 18 | 771 | 99.87 | |
| CHD9 | 0.00 | 0.03 | 2 | 0 | 9 | 1 | 13 | 2606 | 99.96 | |
| ZNHIT6 | 0.00 | 0.03 | 2 | 0 | 7 | 0 | 15 | 467 | 99.79 | |
| NUB1 | 0.00 | 0.03 | 3 | 0 | 4 | 0 | 17 | 621 | 97.18 | Gibbon |
| RPAP2 | 0.00 | 0.03 | 2 | 0 | 5 | 1 | 16 | 612 | 100.00 | |
| LCOR | 0.06 | 0.03 | 6 | 1 | 7 | 2 | 37 | 1554 | 99.87 | Gibbon |
| PLD2 | 0.00 | 0.03 | 6 | 0 | 3 | 0 | 15 | 933 | 100.00 | |
| TMEM108 | 0.00 | 0.04 | 3 | 0 | 4 | 0 | 16 | 518 | 100.00 | Gibbon |
| POLQ | 0.14 | 0.04 | 10 | 3 | 18 | 9 | 67 | 2215 | 90.19 | Gibbon |
| NOX1 | 0.00 | 0.04 | 1 | 0 | 7 | 0 | 15 | 564 | 100.00 | |
| PPEF1 | 0.00 | 0.04 | 2 | 0 | 4 | 2 | 15 | 596 | 97.07 | |
| HHIPL2 | 0.00 | 0.04 | 4 | 0 | 4 | 0 | 15 | 720 | 99.45 | |
| CARF | 0.00 | 0.04 | 8 | 0 | 5 | 1 | 9 | 592 | 100.00 | Human |
| LLGL2 | 0.00 | 0.04 | 3 | 0 | 6 | 1 | 13 | 922 | 93.51 | |
| LRRIQ4 | 0.00 | 0.04 | 3 | 0 | 4 | 0 | 15 | 560 | 100.00 | |
| LETM1 | 0.00 | 0.04 | 3 | 0 | 7 | 0 | 12 | 670 | 94.23 | |
| SON | 0.04 | 0.04 | 9 | 1 | 8 | 0 | 31 | 2355 | 97.84 | |
| ADGRA2 | 0.00 | 0.04 | 3 | 0 | 10 | 0 | 9 | 1028 | 91.13 | |
| PLBD1 | 0.00 | 0.04 | 4 | 0 | 1 | 0 | 16 | 553 | 100.00 | Gibbon |
| ZNF333 | 0.00 | 0.04 | 4 | 0 | 1 | 0 | 16 | 605 | 91.95 | |
| FGD2 | 0.00 | 0.04 | 2 | 0 | 4 | 0 | 15 | 647 | 99.85 | |
| CLGN | 0.00 | 0.04 | 1 | 0 | 5 | 3 | 12 | 537 | 90.10 | |
| SHQ1 | 0.00 | 0.04 | 1 | 0 | 6 | 4 | 10 | 577 | 100.00 | |
| ZNF341 | 0.00 | 0.04 | 2 | 0 | 4 | 1 | 14 | 792 | 100.00 | |
| MAP3K14 | 0.00 | 0.04 | 4 | 0 | 3 | 0 | 14 | 933 | 99.36 | |
| ASAP3 | 0.00 | 0.04 | 2 | 0 | 4 | 2 | 13 | 902 | 99.89 | |
| TMC2 | 0.00 | 0.04 | 6 | 0 | 3 | 0 | 12 | 898 | 99.45 | |
| AGBL3 | 0.00 | 0.04 | 3 | 0 | 6 | 0 | 12 | 919 | 99.89 | |
| CLCN1 | 0.00 | 0.04 | 5 | 0 | 4 | 0 | 12 | 976 | 98.79 | |
| ALS2 | 0.00 | 0.04 | 3 | 0 | 5 | 1 | 12 | 1527 | 93.80 | |
| C10orf71 | 0.16 | 0.04 | 12 | 2 | 7 | 3 | 47 | 1252 | 97.13 | Gibbon |
| ACSL6 | 0.00 | 0.04 | 5 | 0 | 2 | 0 | 13 | 668 | 95.43 | |
| RPUSD2 | 0.00 | 0.04 | 3 | 0 | 6 | 1 | 10 | 536 | 98.35 | |
| LMTK2 | 0.07 | 0.04 | 5 | 1 | 9 | 4 | 26 | 1431 | 95.34 | |
| CFAP58 | 0.00 | 0.04 | 1 | 0 | 6 | 1 | 12 | 869 | 100.00 | |
| RFX7 | 0.00 | 0.04 | 2 | 0 | 4 | 1 | 13 | 1294 | 94.94 | |
| KIAA1211L | 0.12 | 0.04 | 4 | 1 | 5 | 3 | 31 | 814 | 96.67 | Gibbon |
| CORIN | 0.10 | 0.04 | 7 | 1 | 6 | 0 | 29 | 1042 | 100.00 | Gibbon |
| TCTN3 | 0.00 | 0.04 | 2 | 0 | 7 | 2 | 8 | 524 | 96.86 | |
| SLC5A9 | 0.00 | 0.04 | 3 | 0 | 5 | 1 | 10 | 701 | 99.29 | |
| ATP6V0A2 | 0.00 | 0.04 | 1 | 0 | 9 | 0 | 9 | 856 | 100.00 | |
| ATP9B | 0.00 | 0.04 | 1 | 0 | 5 | 1 | 12 | 1034 | 92.40 | |
| HEPH | 0.00 | 0.04 | 2 | 0 | 4 | 2 | 11 | 1154 | 95.37 | |
| BICRA | 0.00 | 0.04 | 6 | 0 | 5 | 0 | 8 | 1080 | 78.49 | |
| TRIP11 | 0.05 | 0.04 | 6 | 1 | 7 | 6 | 22 | 1963 | 99.95 | |
| MAP3K21 | 0.11 | 0.04 | 4 | 1 | 9 | 2 | 25 | 884 | 99.66 | |
| LRRC71 | 0.00 | 0.04 | 1 | 0 | 4 | 0 | 13 | 507 | 100.00 | |
| TRIM66 | 0.08 | 0.04 | 4 | 1 | 10 | 1 | 25 | 1233 | 100.00 | |
| BEST3 | 0.00 | 0.04 | 1 | 0 | 4 | 1 | 12 | 659 | 98.65 | |
| MTA1 | 0.00 | 0.04 | 3 | 0 | 6 | 0 | 9 | 553 | 78.00 | |
| TMC4 | 0.00 | 0.04 | 3 | 0 | 5 | 0 | 10 | 587 | 98.99 | |
| ABI3BP | 0.06 | 0.04 | 9 | 1 | 5 | 0 | 26 | 1651 | 95.88 | |
| TIGD7 | 0.00 | 0.04 | 2 | 0 | 6 | 3 | 7 | 549 | 100.00 | |

| | | | | | | | | | | |
|---|---|---|---|---|---|---|---|---|---|---|
| TSC1 | 0.00 | 0.04 | 4 | 0 | 3 | 1 | 10 | 1164 | 100.00 | |
| STK31 | 0.13 | 0.05 | 5 | 1 | 5 | 7 | 21 | 762 | 90.39 | |
| XIRP1 | 0.09 | 0.05 | 5 | 1 | 8 | 0 | 25 | 1104 | 98.48 | |
| FAM83F | 0.00 | 0.05 | 4 | 0 | 1 | 1 | 11 | 488 | 97.60 | |
| PPP2R3A | 0.09 | 0.05 | 4 | 1 | 9 | 1 | 24 | 1150 | 100.00 | |
| TMEM62 | 0.00 | 0.05 | 1 | 0 | 9 | 0 | 7 | 539 | 92.45 | Gorilla |
| SLC45A4 | 0.00 | 0.05 | 4 | 0 | 1 | 0 | 12 | 744 | 99.73 | |
| UNC5CL | 0.00 | 0.05 | 1 | 0 | 3 | 4 | 9 | 518 | 100.00 | |
| TSR1 | 0.00 | 0.05 | 1 | 0 | 4 | 0 | 12 | 748 | 99.73 | |
| MINDY1 | 0.00 | 0.05 | 6 | 0 | 2 | 1 | 8 | 513 | 99.23 | |
| NCF2 | 0.00 | 0.05 | 5 | 0 | 4 | 0 | 8 | 526 | 100.00 | |
| NTN4 | 0.00 | 0.05 | 3 | 0 | 4 | 0 | 10 | 620 | 99.52 | |
| RFWD3 | 0.00 | 0.05 | 3 | 0 | 4 | 2 | 8 | 728 | 97.33 | |
| PLCL1 | 0.00 | 0.05 | 1 | 0 | 4 | 3 | 9 | 1095 | 100.00 | |
| ZC3H13 | 0.00 | 0.05 | 2 | 0 | 6 | 1 | 8 | 1665 | 99.94 | |
| LAMA5 | 0.12 | 0.05 | 10 | 2 | 14 | 1 | 33 | 1635 | 74.22 | |
| GPR156 | 0.12 | 0.05 | 3 | 1 | 8 | 3 | 23 | 801 | 98.52 | |
| CNTROB | 0.12 | 0.05 | 3 | 1 | 9 | 2 | 23 | 829 | 95.95 | |
| SPIDR | 0.11 | 0.05 | 2 | 1 | 8 | 3 | 24 | 904 | 99.12 | Gibbon |
| ABCG2 | 0.00 | 0.05 | 4 | 0 | 1 | 3 | 8 | 655 | 100.00 | |
| ALDH1L2 | 0.00 | 0.05 | 3 | 0 | 4 | 0 | 9 | 853 | 96.06 | |
| SORL1 | 0.05 | 0.05 | 9 | 1 | 8 | 0 | 19 | 2101 | 96.64 | |
| KMT2E | 0.00 | 0.05 | 1 | 0 | 5 | 1 | 9 | 1630 | 100.00 | |
| FASTKD1 | 0.12 | 0.05 | 2 | 1 | 4 | 3 | 26 | 810 | 100.00 | Gibbon |
| SSFA2 | 0.08 | 0.05 | 5 | 1 | 5 | 2 | 23 | 1243 | 100.00 | |
| HEATR1 | 0.09 | 0.05 | 6 | 2 | 13 | 4 | 31 | 2143 | 100.00 | |
| MOCOS | 0.11 | 0.05 | 4 | 1 | 5 | 1 | 24 | 886 | 99.77 | Gibbon |
| ECT2L | 0.11 | 0.05 | 3 | 1 | 13 | 2 | 16 | 903 | 100.00 | Gorilla |
| PUM3 | 0.00 | 0.05 | 4 | 0 | 1 | 0 | 10 | 647 | 100.00 | |
| CD2AP | 0.00 | 0.05 | 4 | 0 | 2 | 0 | 9 | 639 | 100.00 | |
| LRRC40 | 0.00 | 0.05 | 3 | 0 | 6 | 0 | 6 | 602 | 100.00 | |
| SEMA4C | 0.00 | 0.05 | 1 | 0 | 4 | 0 | 10 | 833 | 100.00 | |
| ACTN3 | 0.00 | 0.05 | 5 | 0 | 3 | 0 | 7 | 902 | 97.83 | |
| SKIV2L | 0.00 | 0.05 | 4 | 0 | 3 | 0 | 8 | 1190 | 97.94 | |
| TAOK2 | 0.00 | 0.05 | 1 | 0 | 5 | 1 | 8 | 1234 | 99.92 | |
| RAB3GAP2 | 0.00 | 0.05 | 2 | 0 | 4 | 0 | 9 | 1364 | 98.13 | |
| CTAGE5 | 0.14 | 0.05 | 5 | 2 | 12 | 1 | 34 | 1409 | 99.79 | Gibbon |
| EXO1 | 0.12 | 0.05 | 7 | 1 | 2 | 0 | 23 | 813 | 96.79 | Gibbon |
| FKBP15 | 0.09 | 0.05 | 7 | 1 | 4 | 1 | 20 | 1129 | 92.62 | |
| HCLS1 | 0.00 | 0.05 | 4 | 0 | 3 | 0 | 7 | 474 | 99.58 | |
| PTPN7 | 0.00 | 0.05 | 2 | 0 | 4 | 1 | 7 | 465 | 100.00 | |
| RFTN1 | 0.00 | 0.05 | 1 | 0 | 6 | 2 | 5 | 479 | 100.00 | |
| TMEM266 | 0.00 | 0.05 | 1 | 0 | 4 | 2 | 7 | 507 | 97.69 | |
| GOLGA5 | 0.00 | 0.05 | 4 | 0 | 2 | 1 | 7 | 619 | 91.70 | |
| TYRO3 | 0.00 | 0.05 | 2 | 0 | 5 | 1 | 6 | 734 | 96.58 | |
| FAM193B | 0.00 | 0.05 | 2 | 0 | 5 | 1 | 6 | 751 | 99.87 | |
| BBX | 0.00 | 0.05 | 4 | 0 | 4 | 0 | 6 | 940 | 100.00 | |
| TBC1D4 | 0.00 | 0.05 | 4 | 0 | 3 | 0 | 7 | 1298 | 100.00 | |
| TRIO | 0.00 | 0.05 | 1 | 0 | 4 | 0 | 9 | 2787 | 97.21 | |
| PCM1 | 0.10 | 0.05 | 10 | 2 | 7 | 3 | 29 | 1945 | 98.93 | |
| TXNDC11 | 0.11 | 0.05 | 4 | 1 | 10 | 1 | 16 | 881 | 99.77 | |
| PLCH2 | 0.15 | 0.05 | 4 | 2 | 7 | 2 | 35 | 1293 | 95.07 | Gibbon |
| IL17RA | 0.12 | 0.06 | 3 | 1 | 7 | 1 | 19 | 809 | 99.02 | |
| IMPG2 | 0.09 | 0.06 | 2 | 1 | 6 | 1 | 21 | 1165 | 98.73 | |
| CEP104 | 0.11 | 0.06 | 4 | 1 | 8 | 4 | 14 | 886 | 99.89 | |
| TOPBP1 | 0.07 | 0.06 | 5 | 1 | 5 | 2 | 18 | 1522 | 100.00 | |
| NLRC4 | 0.12 | 0.06 | 2 | 1 | 4 | 0 | 23 | 861 | 99.88 | Gibbon |
| DHX37 | 0.10 | 0.06 | 4 | 1 | 5 | 0 | 20 | 979 | 91.92 | |
| HPS5 | 0.09 | 0.06 | 3 | 1 | 5 | 0 | 21 | 1100 | 97.43 | |
| PMS1 | 0.11 | 0.06 | 2 | 1 | 6 | 2 | 18 | 929 | 99.89 | |
| MEGF8 | 0.08 | 0.06 | 4 | 2 | 5 | 2 | 33 | 2426 | 94.25 | |
| CSMD1 | 0.11 | 0.06 | 10 | 4 | 13 | 4 | 49 | 3536 | 100.00 | |
| ARAP1 | 0.08 | 0.06 | 3 | 1 | 6 | 2 | 17 | 1260 | 91.04 | |
| CDK12 | 0.07 | 0.06 | 2 | 1 | 7 | 2 | 17 | 1441 | 98.16 | |
| ANKEF1 | 0.13 | 0.06 | 2 | 1 | 6 | 0 | 19 | 776 | 100.00 | |
| PKD2L1 | 0.13 | 0.06 | 3 | 1 | 6 | 3 | 15 | 790 | 98.75 | |
| ZNF628 | 0.10 | 0.06 | 5 | 1 | 7 | 0 | 15 | 994 | 94.58 | |
| ERBIN | 0.07 | 0.06 | 2 | 1 | 6 | 0 | 19 | 1415 | 99.79 | |
| INTS1 | 0.05 | 0.06 | 3 | 1 | 4 | 1 | 19 | 2053 | 96.25 | |
| ARFGEF3 | 0.05 | 0.06 | 2 | 1 | 4 | 2 | 19 | 2177 | 100.00 | |
| ECPAS | 0.05 | 0.06 | 7 | 1 | 4 | 0 | 16 | 1916 | 98.86 | |
| CUL7 | 0.06 | 0.06 | 6 | 1 | 9 | 1 | 11 | 1660 | 95.95 | |
| KIF24 | 0.22 | 0.06 | 5 | 3 | 8 | 0 | 44 | 1348 | 98.54 | Gibbon |
| MICAL1 | 0.18 | 0.06 | 7 | 2 | 7 | 0 | 27 | 1085 | 99.91 | Gibbon |
| TTBK2 | 0.19 | 0.06 | 6 | 3 | 12 | 1 | 37 | 1541 | 98.53 | |
| THSD4 | 0.10 | 0.06 | 2 | 1 | 4 | 0 | 20 | 1016 | 99.90 | |
| SALL3 | 0.08 | 0.06 | 3 | 1 | 5 | 0 | 18 | 1244 | 96.43 | |
| HMGXB3 | 0.08 | 0.06 | 4 | 1 | 5 | 1 | 16 | 1292 | 100.00 | |

| | | | | | | | | | | |
|---|---|---|---|---|---|---|---|---|---|---|
| ZNF687 | 0.08 | 0.06 | 3 | 1 | 5 | 3 | 15 | 1235 | 100.00 | |
| NUP210 | 0.16 | 0.06 | 13 | 3 | 8 | 2 | 32 | 1845 | 97.77 | |
| CHD6 | 0.08 | 0.06 | 3 | 2 | 5 | 2 | 30 | 2661 | 98.70 | |
| CILP | 0.09 | 0.06 | 4 | 1 | 4 | 0 | 17 | 1130 | 97.67 | |
| SALL4 | 0.09 | 0.06 | 5 | 1 | 3 | 1 | 16 | 1053 | 100.00 | |
| CUX2 | 0.08 | 0.06 | 5 | 1 | 4 | 1 | 15 | 1200 | 94.27 | |
| ARHGAP6 | 0.10 | 0.06 | 4 | 1 | 7 | 4 | 10 | 973 | 100.00 | |
| DHX57 | 0.07 | 0.06 | 3 | 1 | 3 | 5 | 14 | 1386 | 100.00 | |
| CSPP1 | 0.19 | 0.06 | 5 | 2 | 8 | 1 | 25 | 1052 | 94.69 | |
| HELB | 0.18 | 0.07 | 4 | 2 | 5 | 4 | 25 | 1087 | 100.00 | |
| PIWIL3 | 0.11 | 0.07 | 3 | 1 | 4 | 1 | 16 | 872 | 98.87 | |
| TRPA1 | 0.09 | 0.07 | 2 | 1 | 5 | 0 | 17 | 1080 | 98.72 | |
| ACIN1 | 0.08 | 0.07 | 3 | 1 | 4 | 0 | 17 | 1293 | 99.31 | |
| TUT1 | 0.11 | 0.07 | 4 | 1 | 6 | 1 | 13 | 912 | 100.00 | |
| NBEAL2 | 0.15 | 0.07 | 12 | 4 | 10 | 5 | 38 | 2591 | 97.11 | |
| AKAP13 | 0.30 | 0.07 | 13 | 8 | 23 | 9 | 74 | 2711 | 98.19 | Gibbon |
| ARHGEF11 | 0.13 | 0.07 | 6 | 2 | 5 | 4 | 22 | 1557 | 100.00 | |
| ABCC10 | 0.15 | 0.07 | 5 | 2 | 10 | 4 | 18 | 1371 | 96.21 | |
| ARHGAP11A | 0.20 | 0.07 | 4 | 2 | 3 | 1 | 27 | 1023 | 100.00 | Gibbon |
| ASXL2 | 0.14 | 0.07 | 6 | 2 | 6 | 0 | 23 | 1387 | 99.86 | |
| GREB1 | 0.12 | 0.07 | 7 | 2 | 6 | 4 | 18 | 1624 | 93.76 | |
| TNKS1BP1 | 0.23 | 0.07 | 10 | 4 | 14 | 7 | 29 | 1728 | 100.00 | |
| GLI2 | 0.20 | 0.07 | 5 | 3 | 12 | 1 | 29 | 1505 | 97.92 | |
| DISP3 | 0.14 | 0.07 | 3 | 2 | 9 | 3 | 19 | 1392 | 100.00 | |
| URB1 | 0.27 | 0.07 | 9 | 6 | 12 | 7 | 56 | 2231 | 98.33 | Gibbon |
| TG | 0.33 | 0.07 | 28 | 9 | 25 | 1 | 67 | 2693 | 98.11 | Human |
| NUP153 | 0.20 | 0.07 | 9 | 3 | 9 | 0 | 28 | 1505 | 99.93 | |
| SVIL | 0.16 | 0.08 | 7 | 3 | 7 | 6 | 24 | 1844 | 94.61 | |
| TNC | 0.18 | 0.08 | 12 | 4 | 8 | 1 | 33 | 2192 | 99.59 | |
| DNAH8 | 0.16 | 0.08 | 11 | 7 | 21 | 3 | 47 | 4500 | 97.72 | |
| LRRK2 | 0.24 | 0.09 | 9 | 6 | 13 | 1 | 47 | 2527 | 100.00 | |
| SYNE1 | 0.16 | 0.09 | 34 | 14 | 24 | 7 | 88 | 8735 | 99.44 | |
| FRAS1 | 0.30 | 0.10 | 20 | 12 | 21 | 7 | 70 | 3948 | 99.37 | |

**S4 Table**: Lower than expected substitution rate on the gorilla branch

| Gene | Gorilla % subs per site | Gorilla norm branch length | Human subs | Chimp subs | Gorilla subs | #1# subs | Gibbon subs | Align overlap | Align Sat | Higher in |
|---|---|---|---|---|---|---|---|---|---|---|
| TUBGCP6 | 0.00 | 0.02 | 2 | 18 | 0 | 1 | 18 | 690 | 47.85 | Chimpanzee |
| MASP1 | 0.00 | 0.02 | 1 | 32 | 0 | 0 | 5 | 518 | 74.11 | Chimpanzee |
| PSD3 | 0.00 | 0.03 | 5 | 3 | 0 | 0 | 20 | 911 | 96.00 | |
| LTBP4 | 0.00 | 0.03 | 3 | 6 | 0 | 0 | 19 | 1483 | 94.22 | |
| GPR50 | 0.00 | 0.03 | 4 | 7 | 0 | 0 | 16 | 617 | 100.00 | |
| SENP6 | 0.00 | 0.03 | 2 | 4 | 0 | 1 | 19 | 1112 | 100.00 | |
| LRP1 | 0.00 | 0.03 | 6 | 2 | 0 | 0 | 17 | 3937 | 93.32 | |
| TMEM131 | 0.00 | 0.04 | 4 | 4 | 0 | 2 | 13 | 1882 | 99.95 | |
| LMLN2 | 0.00 | 0.04 | 6 | 1 | 0 | 1 | 14 | 725 | 92.24 | |
| SMC5 | 0.00 | 0.04 | 4 | 1 | 0 | 0 | 17 | 1061 | 98.61 | |
| SOWAHC | 0.00 | 0.04 | 8 | 2 | 0 | 1 | 10 | 508 | 99.61 | Human |
| NUP98 | 0.00 | 0.04 | 3 | 4 | 0 | 0 | 14 | 1486 | 98.87 | |
| ADAM12 | 0.22 | 0.04 | 3 | 5 | 2 | 3 | 57 | 906 | 99.67 | Gibbon |
| CCDC102B | 0.00 | 0.04 | 4 | 5 | 0 | 0 | 11 | 513 | 100.00 | |
| TXNDC16 | 0.00 | 0.04 | 4 | 2 | 0 | 2 | 12 | 791 | 99.87 | |
| SLC12A7 | 0.00 | 0.04 | 5 | 2 | 0 | 1 | 12 | 1028 | 97.16 | |
| SUGP2 | 0.00 | 0.04 | 4 | 3 | 0 | 1 | 12 | 1020 | 93.41 | |
| PLEKHS1 | 0.00 | 0.04 | 2 | 4 | 0 | 0 | 13 | 462 | 99.57 | |
| AK7 | 0.00 | 0.04 | 4 | 3 | 0 | 0 | 12 | 687 | 99.71 | |
| KL | 0.00 | 0.04 | 2 | 4 | 0 | 1 | 12 | 739 | 100.00 | |
| EPHA1 | 0.00 | 0.04 | 4 | 2 | 0 | 1 | 12 | 975 | 100.00 | |
| MTHFSD | 0.00 | 0.04 | 1 | 5 | 0 | 0 | 12 | 383 | 100.00 | |
| PLIN2 | 0.00 | 0.04 | 3 | 4 | 0 | 0 | 11 | 405 | 92.68 | |
| NR1I2 | 0.00 | 0.04 | 4 | 3 | 0 | 0 | 11 | 469 | 99.15 | |
| VRK3 | 0.00 | 0.04 | 2 | 6 | 0 | 0 | 10 | 474 | 100.00 | |
| ALKBH8 | 0.00 | 0.04 | 5 | 5 | 0 | 1 | 7 | 661 | 99.55 | |
| ANKFY1 | 0.00 | 0.05 | 2 | 5 | 0 | 1 | 9 | 1101 | 97.69 | |
| FBF1 | 0.13 | 0.05 | 5 | 3 | 1 | 1 | 28 | 777 | 83.01 | Gibbon |
| ABAT | 0.00 | 0.05 | 4 | 2 | 0 | 0 | 10 | 456 | 88.54 | |
| MKKS | 0.00 | 0.05 | 1 | 5 | 0 | 0 | 10 | 570 | 100.00 | |
| ATF6B | 0.00 | 0.05 | 3 | 5 | 0 | 0 | 8 | 703 | 100.00 | |
| COL10A1 | 0.00 | 0.05 | 7 | 3 | 0 | 1 | 5 | 680 | 100.00 | |
| RRBP1 | 0.08 | 0.05 | 6 | 7 | 1 | 1 | 21 | 1201 | 87.09 | |
| SLC7A10 | 0.00 | 0.05 | 3 | 5 | 0 | 0 | 7 | 511 | 99.03 | |
| ERMAP | 0.00 | 0.05 | 3 | 1 | 0 | 4 | 7 | 475 | 100.00 | |
| DAG1 | 0.00 | 0.05 | 4 | 1 | 0 | 0 | 10 | 894 | 100.00 | |
| WIZ | 0.00 | 0.05 | 7 | 1 | 0 | 1 | 6 | 702 | 80.05 | |
| PGAP1 | 0.00 | 0.05 | 5 | 4 | 0 | 0 | 6 | 922 | 100.00 | |
| ATR | 0.04 | 0.05 | 7 | 4 | 1 | 4 | 19 | 2612 | 99.69 | |
| MYO6 | 0.00 | 0.05 | 5 | 1 | 0 | 1 | 8 | 1267 | 98.22 | |
| KIAA0408 | 0.14 | 0.05 | 5 | 2 | 1 | 1 | 25 | 694 | 100.00 | Gibbon |
| KIAA0825 | 0.16 | 0.05 | 14 | 8 | 2 | 0 | 29 | 1232 | 97.70 | Human |
| MCF2L2 | 0.09 | 0.05 | 8 | 5 | 1 | 2 | 17 | 1089 | 98.64 | |
| SORBS2 | 0.08 | 0.05 | 6 | 9 | 1 | 0 | 17 | 1193 | 99.42 | |
| DHX38 | 0.00 | 0.05 | 1 | 4 | 0 | 0 | 9 | 1218 | 99.27 | |
| ZNF629 | 0.00 | 0.05 | 5 | 3 | 0 | 1 | 5 | 866 | 99.65 | |
| SCAF1 | 0.00 | 0.05 | 4 | 2 | 0 | 1 | 7 | 1042 | 88.23 | |
| HECW2 | 0.00 | 0.05 | 2 | 5 | 0 | 0 | 7 | 1572 | 100.00 | |
| MMRN1 | 0.16 | 0.05 | 5 | 13 | 2 | 0 | 31 | 1228 | 100.00 | |
| USPL1 | 0.18 | 0.05 | 9 | 3 | 2 | 0 | 36 | 1091 | 99.91 | Gibbon |
| METTL4 | 0.00 | 0.06 | 4 | 0 | 0 | 0 | 9 | 423 | 100.00 | |
| PRF1 | 0.00 | 0.06 | 0 | 4 | 0 | 1 | 8 | 491 | 95.71 | |
| CNPPD1 | 0.00 | 0.06 | 5 | 3 | 0 | 0 | 5 | 396 | 96.59 | |
| ITGB4 | 0.07 | 0.06 | 6 | 3 | 1 | 0 | 21 | 1521 | 87.11 | |
| TTC38 | 0.00 | 0.06 | 4 | 3 | 0 | 1 | 5 | 411 | 92.57 | |
| DHX35 | 0.00 | 0.06 | 6 | 3 | 0 | 0 | 4 | 675 | 97.40 | |
| LARP1 | 0.00 | 0.06 | 2 | 7 | 0 | 0 | 4 | 907 | 90.07 | |
| PRDM10 | 0.00 | 0.06 | 6 | 1 | 0 | 0 | 6 | 1158 | 100.00 | |
| CRB2 | 0.22 | 0.06 | 5 | 6 | 2 | 1 | 34 | 926 | 87.28 | Gibbon |
| PCSK5 | 0.11 | 0.06 | 5 | 10 | 2 | 0 | 31 | 1825 | 100.00 | |
| AOAH | 0.15 | 0.06 | 4 | 4 | 1 | 1 | 20 | 671 | 98.24 | Gibbon |
| TLR3 | 0.11 | 0.06 | 2 | 8 | 1 | 0 | 19 | 904 | 100.00 | |
| ARHGEF2 | 0.10 | 0.06 | 5 | 5 | 1 | 0 | 19 | 1030 | 99.90 | |
| ZNF598 | 0.12 | 0.06 | 8 | 4 | 1 | 2 | 14 | 829 | 100.00 | |
| FBLN7 | 0.00 | 0.06 | 4 | 2 | 0 | 0 | 6 | 439 | 100.00 | |
| MAP3K11 | 0.00 | 0.06 | 4 | 1 | 0 | 1 | 6 | 847 | 100.00 | |
| JARID2 | 0.00 | 0.06 | 5 | 1 | 0 | 0 | 6 | 1177 | 96.08 | |
| MN1 | 0.00 | 0.06 | 4 | 2 | 0 | 0 | 6 | 1104 | 99.37 | |
| SEPT4 | 0.20 | 0.06 | 8 | 3 | 2 | 3 | 29 | 991 | 99.50 | Gibbon |
| TDRD15 | 0.25 | 0.06 | 11 | 8 | 4 | 1 | 54 | 1604 | 99.75 | Gibbon |
| TTC3 | 0.21 | 0.06 | 13 | 11 | 4 | 4 | 46 | 1946 | 97.79 | Gibbon |
| FRMD7 | 0.14 | 0.06 | 2 | 4 | 1 | 3 | 18 | 713 | 100.00 | |
| TMEM132A | 0.10 | 0.06 | 7 | 3 | 1 | 1 | 16 | 994 | 97.26 | |
| APAF1 | 0.08 | 0.06 | 4 | 3 | 1 | 2 | 18 | 1248 | 100.00 | |

| | | | | | | | | | | |
|---|---|---|---|---|---|---|---|---|---|---|
| ARID2 | 0.06 | 0.06 | 4 | 2 | 1 | 0 | 21 | 1742 | 97.76 | |
| SLC7A9 | 0.00 | 0.06 | 4 | 0 | 0 | 0 | 7 | 458 | 94.05 | |
| ZFP64 | 0.00 | 0.06 | 4 | 0 | 0 | 0 | 7 | 623 | 94.97 | |
| NEMP1 | 0.00 | 0.06 | 1 | 1 | 0 | 4 | 5 | 443 | 99.77 | |
| SLC7A13 | 0.00 | 0.06 | 1 | 4 | 0 | 1 | 5 | 470 | 100.00 | |
| PACS2 | 0.00 | 0.06 | 2 | 4 | 0 | 0 | 5 | 665 | 81.80 | |
| RBL1 | 0.00 | 0.06 | 4 | 0 | 0 | 0 | 7 | 1066 | 100.00 | |
| ADAM22 | 0.00 | 0.06 | 4 | 1 | 0 | 0 | 6 | 906 | 100.00 | |
| SDCCAG8 | 0.14 | 0.06 | 4 | 2 | 1 | 1 | 18 | 704 | 98.74 | |
| CGN | 0.09 | 0.06 | 4 | 4 | 1 | 1 | 16 | 1072 | 98.35 | |
| KIFC2 | 0.13 | 0.07 | 7 | 2 | 1 | 0 | 15 | 784 | 99.37 | |
| UBA7 | 0.11 | 0.07 | 4 | 2 | 1 | 3 | 15 | 893 | 92.16 | |
| ABCA2 | 0.10 | 0.07 | 3 | 8 | 2 | 1 | 26 | 2023 | 85.58 | |
| LTBP1 | 0.06 | 0.07 | 8 | 5 | 1 | 2 | 9 | 1689 | 99.47 | |
| CTC1 | 0.20 | 0.07 | 5 | 4 | 2 | 2 | 26 | 1019 | 87.92 | Gibbon |
| ARHGAP29 | 0.16 | 0.07 | 4 | 4 | 2 | 1 | 28 | 1261 | 100.00 | |
| PBXIP1 | 0.15 | 0.07 | 5 | 3 | 1 | 0 | 15 | 685 | 96.61 | |
| CCDC146 | 0.10 | 0.07 | 4 | 1 | 1 | 2 | 16 | 955 | 100.00 | |
| NBAS | 0.13 | 0.07 | 7 | 15 | 3 | 0 | 28 | 2353 | 99.58 | |
| CACNA1A | 0.05 | 0.07 | 2 | 2 | 1 | 9 | 10 | 1994 | 83.12 | |
| HEATR5A | 0.10 | 0.07 | 7 | 5 | 2 | 0 | 24 | 2046 | 100.00 | |
| THEMIS | 0.15 | 0.07 | 5 | 2 | 1 | 1 | 14 | 680 | 100.00 | |
| AP4E1 | 0.09 | 0.07 | 1 | 4 | 1 | 0 | 17 | 1056 | 99.44 | |
| ESYT3 | 0.11 | 0.07 | 2 | 5 | 1 | 0 | 15 | 877 | 99.89 | |
| NHS | 0.07 | 0.07 | 4 | 3 | 1 | 2 | 13 | 1441 | 97.76 | |
| MIA3 | 0.32 | 0.07 | 11 | 13 | 6 | 2 | 59 | 1880 | 98.58 | Gibbon |
| PPRC1 | 0.18 | 0.07 | 5 | 8 | 3 | 3 | 31 | 1638 | 98.44 | |
| CACNA1G | 0.09 | 0.07 | 2 | 5 | 2 | 0 | 26 | 2234 | 96.42 | |
| TEP1 | 0.28 | 0.07 | 13 | 16 | 7 | 2 | 63 | 2502 | 98.00 | Gibbon |
| EPG5 | 0.12 | 0.08 | 4 | 6 | 3 | 3 | 32 | 2558 | 99.73 | |
| KIF14 | 0.24 | 0.08 | 7 | 5 | 4 | 2 | 42 | 1648 | 100.00 | Gibbon |
| OLFML2A | 0.15 | 0.08 | 4 | 2 | 1 | 0 | 14 | 651 | 100.00 | |
| PPFIBP1 | 0.10 | 0.08 | 2 | 4 | 1 | 1 | 13 | 956 | 95.41 | |
| ANKFN1 | 0.17 | 0.08 | 7 | 7 | 2 | 2 | 16 | 1144 | 99.65 | |
| ARAP2 | 0.18 | 0.08 | 4 | 10 | 3 | 0 | 28 | 1704 | 100.00 | |
| TCF3 | 0.15 | 0.08 | 4 | 3 | 1 | 1 | 11 | 651 | 96.73 | |
| BIRC6 | 0.04 | 0.08 | 9 | 4 | 2 | 0 | 17 | 4836 | 99.57 | |
| PPL | 0.30 | 0.08 | 7 | 11 | 5 | 0 | 45 | 1683 | 98.13 | Gibbon |
| CACNA1B | 0.11 | 0.08 | 2 | 14 | 2 | 1 | 12 | 1833 | 87.20 | |
| CASKIN1 | 0.18 | 0.09 | 2 | 5 | 2 | 3 | 18 | 1124 | 89.42 | |
| WWC2 | 0.18 | 0.09 | 4 | 6 | 2 | 1 | 17 | 1128 | 98.34 | |
| TET3 | 0.12 | 0.09 | 6 | 7 | 2 | 0 | 15 | 1735 | 98.58 | |
| ADGRG6 | 0.16 | 0.09 | 4 | 2 | 2 | 0 | 21 | 1223 | 99.92 | |
| ADGRD1 | 0.23 | 0.09 | 4 | 5 | 2 | 1 | 17 | 869 | 100.00 | |
| AFF1 | 0.26 | 0.09 | 4 | 6 | 3 | 0 | 27 | 1172 | 98.90 | |
| HTT | 0.13 | 0.09 | 9 | 6 | 4 | 2 | 30 | 2976 | 97.64 | |
| TANGO6 | 0.28 | 0.09 | 4 | 4 | 3 | 0 | 28 | 1082 | 98.90 | Gibbon |
| WNK1 | 0.18 | 0.09 | 10 | 8 | 5 | 1 | 37 | 2785 | 99.96 | |
| FHOD3 | 0.19 | 0.09 | 4 | 4 | 3 | 0 | 27 | 1567 | 100.00 | |
| PLCH1 | 0.18 | 0.09 | 7 | 7 | 3 | 2 | 19 | 1693 | 100.00 | |
| C2CD3 | 0.31 | 0.09 | 9 | 8 | 7 | 3 | 53 | 2243 | 97.61 | Gibbon |
| COL22A1 | 0.30 | 0.09 | 11 | 6 | 4 | 2 | 25 | 1342 | 85.86 | |
| PFAS | 0.28 | 0.09 | 5 | 6 | 3 | 1 | 22 | 1078 | 97.29 | |
| HIVEP1 | 0.30 | 0.10 | 18 | 10 | 8 | 4 | 49 | 2688 | 99.37 | |
| CEP152 | 0.35 | 0.10 | 11 | 9 | 6 | 3 | 36 | 1709 | 99.94 | |
| ZNF646 | 0.28 | 0.10 | 4 | 6 | 5 | 0 | 39 | 1802 | 99.78 | |
| ADAMTSL3 | 0.30 | 0.10 | 11 | 7 | 5 | 0 | 31 | 1684 | 100.00 | |
| DCHS1 | 0.15 | 0.10 | 5 | 11 | 5 | 1 | 32 | 3297 | 100.00 | |
| VPS13C | 0.19 | 0.10 | 15 | 16 | 7 | 1 | 34 | 3636 | 99.32 | |
| LYST | 0.24 | 0.10 | 15 | 16 | 9 | 0 | 52 | 3736 | 98.58 | |
| HYDIN | 0.46 | 0.11 | 21 | 24 | 23 | 5 | 139 | 5005 | 98.48 | Gibbon |
| DNAH9 | 0.35 | 0.11 | 15 | 18 | 15 | 5 | 84 | 4247 | 96.24 | |
| FAT1 | 0.36 | 0.11 | 30 | 21 | 16 | 2 | 74 | 4401 | 97.56 | |

**S5 Table**: Higher than expected substitution rate on the human branch

| Gene | Human % subs per site | Human norm branch length | Human subs | Chimp subs | Gorilla subs | #1# subs | Gibbon subs | Align overlap | Align Sat | Lower in |
|---|---|---|---|---|---|---|---|---|---|---|
| ADCYAP1 | 7.60 | 0.74 | 13 | 0 | 0 | 0 | 1 | 171 | 97.16 | |
| BIRC5 | 4.40 | 0.42 | 4 | 0 | 1 | 1 | 1 | 91 | 100.00 | |
| DACT3 | 3.89 | 0.67 | 7 | 0 | 0 | 0 | 0 | 180 | 99.45 | Gibbon |
| PVALEF | 3.76 | 0.40 | 5 | 0 | 0 | 0 | 5 | 133 | 99.25 | |
| PGLYRP1 | 3.57 | 0.33 | 7 | 1 | 5 | 0 | 6 | 196 | 100.00 | |
| PSORS1C2 | 2.94 | 0.42 | 4 | 1 | 1 | 0 | 1 | 136 | 100.00 | |
| BTNL2 | 2.88 | 0.37 | 7 | 2 | 2 | 1 | 5 | 243 | 89.67 | |
| DMKN | 2.59 | 0.29 | 9 | 1 | 3 | 0 | 16 | 347 | 74.62 | |
| TMED3 | 2.48 | 0.55 | 5 | 0 | 0 | 0 | 1 | 202 | 93.09 | |
| DCXR | 2.46 | 0.41 | 6 | 0 | 1 | 0 | 5 | 244 | 100.00 | |
| DHDH | 2.40 | 0.48 | 8 | 2 | 0 | 0 | 4 | 333 | 99.70 | |
| BLVRB | 2.22 | 0.50 | 4 | 0 | 1 | 0 | 0 | 180 | 87.38 | Gibbon |
| CD37 | 2.16 | 0.35 | 6 | 3 | 2 | 0 | 4 | 278 | 98.93 | |
| TMBIM4 | 2.12 | 0.50 | 6 | 0 | 0 | 0 | 3 | 283 | 100.00 | |
| ODF1 | 2.00 | 0.46 | 5 | 0 | 1 | 0 | 2 | 250 | 100.00 | |
| NACAD | 1.86 | 0.23 | 19 | 11 | 8 | 0 | 45 | 1019 | 95.06 | |
| VWA2 | 1.77 | 0.26 | 10 | 3 | 4 | 4 | 17 | 566 | 86.94 | |
| EPCAM | 1.71 | 0.43 | 5 | 1 | 0 | 0 | 3 | 292 | 91.25 | |
| MRPL39 | 1.70 | 0.32 | 6 | 2 | 2 | 1 | 6 | 352 | 99.72 | |
| FAM118A | 1.68 | 0.37 | 6 | 2 | 0 | 0 | 6 | 357 | 100.00 | |
| CA6 | 1.62 | 0.33 | 5 | 2 | 2 | 1 | 3 | 309 | 99.04 | |
| KIF26B | 1.58 | 0.31 | 24 | 8 | 6 | 2 | 35 | 1518 | 98.64 | |
| SOWAHC | 1.57 | 0.35 | 8 | 2 | 0 | 1 | 10 | 508 | 99.61 | Gorilla |
| EDA2R | 1.57 | 0.43 | 5 | 1 | 1 | 0 | 2 | 318 | 100.00 | |
| MAP7D1 | 1.56 | 0.33 | 10 | 1 | 1 | 0 | 16 | 640 | 87.19 | |
| COL25A1 | 1.53 | 0.36 | 10 | 4 | 3 | 1 | 8 | 653 | 99.85 | |
| RBM44 | 1.52 | 0.29 | 16 | 6 | 7 | 0 | 25 | 1050 | 99.81 | |
| SLC7A2 | 1.49 | 0.31 | 10 | 5 | 2 | 0 | 13 | 671 | 96.41 | |
| PGM1 | 1.48 | 0.67 | 7 | 0 | 0 | 0 | 0 | 474 | 81.72 | Gibbon |
| IQCA1L | 1.40 | 0.27 | 10 | 8 | 4 | 0 | 14 | 715 | 99.58 | |
| CARF | 1.35 | 0.32 | 8 | 0 | 5 | 1 | 9 | 592 | 100.00 | Chimpanzee |
| NOM1 | 1.34 | 0.24 | 11 | 6 | 8 | 3 | 16 | 822 | 95.80 | |
| ALDH16A1 | 1.24 | 0.29 | 8 | 2 | 4 | 4 | 8 | 647 | 97.29 | |
| SLC15A5 | 1.22 | 0.31 | 7 | 3 | 3 | 0 | 8 | 575 | 99.48 | |
| BRCA1 | 1.19 | 0.24 | 21 | 6 | 12 | 0 | 47 | 1771 | 97.09 | |
| SMG6 | 1.16 | 0.41 | 16 | 3 | 3 | 2 | 13 | 1383 | 98.93 | |
| KIAA0825 | 1.14 | 0.26 | 14 | 8 | 2 | 0 | 29 | 1232 | 97.70 | Gorilla |
| RET | 1.09 | 0.36 | 12 | 7 | 3 | 0 | 9 | 1097 | 98.47 | Gibbon |
| GRIP2 | 1.06 | 0.33 | 12 | 6 | 7 | 0 | 10 | 1132 | 99.30 | Gibbon |
| TG | 1.04 | 0.21 | 28 | 9 | 25 | 1 | 67 | 2693 | 98.11 | Chimpanzee |
| MYO18B | 0.99 | 0.24 | 22 | 13 | 10 | 4 | 41 | 2217 | 96.94 | |
| MYOM1 | 0.98 | 0.44 | 13 | 3 | 2 | 1 | 8 | 1333 | 97.16 | Gibbon |

**S6 Table**: Higher than expected substitution rate on the chimpanzee branch

| Gene | Chimp % subs per site | Chimp norm branch length | Human subs | Chimp subs | Gorilla subs | #1# subs | Gibbon subs | Align overlap | Align Sat | Lower in |
|---|---|---|---|---|---|---|---|---|---|---|
| SCART1 | 7.85 | 0.55 | 11 | 52 | 6 | 0 | 23 | 662 | 72.51 | |
| ASIC1 | 6.51 | 0.84 | 0 | 20 | 0 | 0 | 0 | 307 | 64.09 | Gibbon |
| CALU | 6.25 | 0.84 | 0 | 20 | 0 | 0 | 0 | 320 | 99.07 | Gibbon |
| MASP1 | 6.18 | 0.77 | 1 | 32 | 0 | 0 | 5 | 518 | 74.11 | Gorilla |
| COL23A1 | 6.09 | 0.52 | 0 | 14 | 1 | 0 | 9 | 230 | 56.10 | |
| SCML1 | 5.21 | 0.36 | 5 | 17 | 5 | 2 | 16 | 326 | 99.09 | |
| SRI | 4.14 | 0.58 | 0 | 7 | 1 | 0 | 1 | 169 | 85.35 | |
| SLC22A31 | 3.65 | 0.35 | 4 | 5 | 1 | 0 | 2 | 137 | 40.53 | |
| CXorf67 | 3.11 | 0.29 | 1 | 12 | 4 | 3 | 20 | 386 | 97.97 | |
| CARD19 | 2.99 | 0.50 | 0 | 5 | 0 | 0 | 2 | 167 | 91.26 | |
| C11orf74 | 2.92 | 0.33 | 1 | 5 | 1 | 0 | 6 | 171 | 87.24 | |
| UPRT | 2.90 | 0.56 | 0 | 4 | 0 | 0 | 0 | 138 | 79.77 | Gibbon |
| NCS1 | 2.84 | 0.56 | 0 | 4 | 0 | 0 | 0 | 141 | 81.98 | |
| LYAR | 2.65 | 0.38 | 2 | 10 | 4 | 0 | 8 | 378 | 100.00 | |
| PIMREG | 2.61 | 0.38 | 1 | 7 | 1 | 0 | 7 | 268 | 98.17 | |
| TUBGCP6 | 2.61 | 0.43 | 2 | 18 | 0 | 1 | 18 | 690 | 47.85 | Gorilla |
| SPDYE4 | 2.53 | 0.32 | 0 | 6 | 1 | 0 | 10 | 237 | 100.00 | |
| PEX11G | 2.49 | 0.41 | 1 | 6 | 2 | 0 | 3 | 241 | 100.00 | |
| RNF168 | 2.48 | 0.30 | 2 | 9 | 2 | 0 | 15 | 363 | 77.23 | |
| HM13 | 2.48 | 0.72 | 0 | 9 | 0 | 0 | 0 | 363 | 85.21 | Gibbon |
| DFFA | 2.44 | 0.32 | 2 | 8 | 2 | 0 | 11 | 328 | 100.00 | |
| ANKLE2 | 2.22 | 0.33 | 3 | 5 | 3 | 1 | 1 | 225 | 76.53 | |
| DDX47 | 2.14 | 0.53 | 0 | 9 | 1 | 0 | 4 | 420 | 92.31 | |
| FAM53B | 2.03 | 0.50 | 0 | 6 | 1 | 0 | 2 | 296 | 86.55 | |
| MTG1 | 2.02 | 0.35 | 1 | 5 | 1 | 0 | 5 | 248 | 82.67 | |
| DERL3 | 1.93 | 0.39 | 0 | 4 | 3 | 0 | 1 | 207 | 88.09 | |
| PCBP3 | 1.92 | 0.55 | 0 | 5 | 0 | 0 | 1 | 260 | 78.31 | Gibbon |
| COX10 | 1.90 | 0.35 | 0 | 6 | 2 | 0 | 7 | 316 | 79.40 | |
| NIPAL3 | 1.86 | 0.50 | 0 | 6 | 1 | 0 | 2 | 322 | 87.50 | |
| ULK1 | 1.86 | 0.50 | 2 | 12 | 2 | 0 | 5 | 646 | 76.00 | Gibbon |
| ZNF280C | 1.68 | 0.30 | 3 | 12 | 8 | 4 | 12 | 715 | 99.86 | |
| DNAJC9 | 1.66 | 0.46 | 0 | 4 | 1 | 0 | 1 | 241 | 97.57 | Gibbon |
| ARHGAP40 | 1.63 | 0.29 | 1 | 9 | 2 | 0 | 17 | 553 | 100.00 | |
| QRFPR | 1.62 | 0.35 | 0 | 7 | 3 | 0 | 8 | 431 | 100.00 | |
| SPG7 | 1.61 | 0.38 | 3 | 10 | 3 | 1 | 7 | 621 | 80.34 | |
| CYP24A1 | 1.61 | 0.35 | 1 | 7 | 2 | 1 | 7 | 436 | 97.32 | |
| ABR | 1.59 | 0.69 | 0 | 10 | 0 | 0 | 1 | 628 | 74.76 | |
| ADAMTS10 | 1.57 | 0.58 | 3 | 13 | 0 | 0 | 3 | 828 | 78.78 | |
| TAF6L | 1.52 | 0.47 | 0 | 6 | 0 | 0 | 4 | 395 | 68.34 | |
| NMUR2 | 1.50 | 0.32 | 4 | 6 | 1 | 0 | 6 | 401 | 99.50 | |
| CAPS2 | 1.50 | 0.35 | 3 | 7 | 2 | 0 | 6 | 468 | 97.91 | |
| CPNE7 | 1.47 | 0.37 | 2 | 6 | 1 | 0 | 5 | 408 | 77.57 | |
| CARS2 | 1.42 | 0.27 | 1 | 8 | 4 | 1 | 14 | 564 | 100.00 | |
| EHMT1 | 1.40 | 0.59 | 1 | 12 | 0 | 0 | 4 | 857 | 77.91 | |
| GMPS | 1.38 | 0.57 | 0 | 7 | 1 | 0 | 1 | 506 | 86.20 | Gibbon |
| NEURL1 | 1.35 | 0.64 | 0 | 6 | 0 | 0 | 0 | 446 | 82.14 | Gibbon |
| ACOX3 | 1.29 | 0.38 | 2 | 8 | 2 | 0 | 7 | 620 | 89.86 | |
| CFAP54 | 1.25 | 0.31 | 2 | 7 | 2 | 1 | 9 | 558 | 85.71 | |
| DDIAS | 1.21 | 0.26 | 4 | 12 | 4 | 0 | 25 | 995 | 99.70 | |
| BLM | 0.93 | 0.24 | 6 | 13 | 10 | 1 | 24 | 1394 | 99.64 | |

**S7 Table**: Higher than expected substitution rate on the gorilla branch

| Gene | Gorilla % subs per site | Gorilla norm branch length | Human subs | Chimp subs | Gorilla subs | #1# subs | Gibbon subs | Align overlap | Align Sat | Lower in |
|---|---|---|---|---|---|---|---|---|---|---|
| PPDPF | 10.91 | 0.65 | 0 | 0 | 6 | 0 | 0 | 55 | 50.00 | |
| RNF128 | 7.26 | 0.76 | 1 | 0 | 23 | 0 | 3 | 317 | 78.86 | |
| TRIM14 | 6.85 | 0.40 | 0 | 1 | 5 | 0 | 4 | 73 | 66.36 | |
| PRAC2 | 5.15 | 0.36 | 1 | 0 | 7 | 1 | 8 | 136 | 97.14 | |
| KCTD18 | 4.52 | 0.54 | 1 | 2 | 16 | 1 | 7 | 354 | 88.06 | |
| ZDHHC3 | 4.38 | 0.73 | 0 | 0 | 12 | 1 | 0 | 274 | 82.28 | Gibbon |
| SNAP25 | 4.37 | 0.72 | 0 | 0 | 9 | 0 | 0 | 206 | 100.00 | Gibbon |
| CARD8 | 3.67 | 0.31 | 8 | 3 | 13 | 0 | 16 | 354 | 87.19 | |
| RCC1 | 3.37 | 0.52 | 1 | 0 | 12 | 1 | 6 | 356 | 84.16 | |
| WDR38 | 3.30 | 0.41 | 4 | 0 | 10 | 0 | 8 | 303 | 98.70 | |
| NOX4 | 3.23 | 0.62 | 0 | 0 | 12 | 0 | 4 | 371 | 85.48 | |
| MKRN1 | 3.17 | 0.60 | 1 | 1 | 15 | 1 | 4 | 473 | 98.13 | |
| PALM3 | 3.15 | 0.31 | 1 | 5 | 19 | 4 | 31 | 604 | 92.35 | |
| APOOL | 3.00 | 0.54 | 0 | 0 | 6 | 0 | 2 | 200 | 86.58 | |
| RFPL2 | 2.97 | 0.30 | 5 | 3 | 10 | 4 | 10 | 337 | 99.41 | |
| R3HDML | 2.92 | 0.37 | 2 | 0 | 7 | 2 | 6 | 240 | 100.00 | |
| PLAUR | 2.90 | 0.39 | 1 | 1 | 6 | 0 | 5 | 207 | 79.31 | |
| SAPCD2 | 2.90 | 0.48 | 1 | 1 | 10 | 0 | 6 | 345 | 96.64 | |
| HGFAC | 2.88 | 0.45 | 1 | 7 | 18 | 0 | 11 | 625 | 96.60 | |
| SPATA32 | 2.88 | 0.33 | 3 | 1 | 11 | 3 | 13 | 382 | 100.00 | |
| NSMCE4A | 2.83 | 0.45 | 0 | 1 | 7 | 0 | 5 | 247 | 75.30 | |
| QRICH2 | 2.66 | 0.27 | 15 | 14 | 40 | 10 | 69 | 1502 | 88.46 | |
| AMACR | 2.65 | 0.36 | 3 | 3 | 10 | 1 | 9 | 377 | 98.69 | |
| ANAPC11 | 2.59 | 0.40 | 1 | 1 | 5 | 0 | 3 | 193 | 99.48 | |
| THEGL | 2.59 | 0.34 | 1 | 5 | 12 | 1 | 14 | 464 | 100.00 | |
| RITA1 | 2.56 | 0.40 | 2 | 1 | 5 | 1 | 1 | 195 | 98.48 | |
| FGF11 | 2.54 | 0.50 | 0 | 0 | 5 | 1 | 1 | 197 | 100.00 | |
| ANO8 | 2.47 | 0.58 | 0 | 0 | 10 | 0 | 4 | 405 | 69.71 | |
| IQCD | 2.45 | 0.36 | 1 | 2 | 11 | 2 | 12 | 449 | 100.00 | |
| DYTN | 2.45 | 0.27 | 4 | 3 | 13 | 1 | 25 | 531 | 99.62 | |
| TBXA2R | 2.44 | 0.57 | 1 | 0 | 8 | 0 | 2 | 328 | 95.63 | Gibbon |
| DRD4 | 2.40 | 0.50 | 0 | 1 | 8 | 0 | 4 | 334 | 86.53 | |
| SLC25A24 | 2.29 | 0.65 | 0 | 0 | 10 | 0 | 2 | 436 | 95.20 | Gibbon |
| FAM161B | 2.29 | 0.34 | 4 | 2 | 16 | 3 | 20 | 700 | 99.43 | |
| ITPKA | 2.24 | 0.67 | 0 | 0 | 7 | 0 | 0 | 312 | 75.54 | Gibbon |
| SRF | 2.23 | 0.65 | 0 | 1 | 8 | 0 | 0 | 358 | 75.69 | Gibbon |
| CCDC112 | 2.18 | 0.55 | 3 | 0 | 11 | 0 | 3 | 505 | 98.44 | |
| MRPL46 | 2.15 | 0.35 | 2 | 2 | 6 | 1 | 4 | 279 | 100.00 | |
| HMGCLL1 | 2.09 | 0.46 | 0 | 0 | 5 | 0 | 3 | 239 | 78.88 | |
| FBXW8 | 2.09 | 0.60 | 0 | 0 | 11 | 1 | 3 | 527 | 98.69 | |
| GRIN3B | 2.07 | 0.29 | 7 | 3 | 15 | 3 | 22 | 726 | 75.39 | |
| TYSND1 | 2.05 | 0.32 | 4 | 6 | 11 | 0 | 11 | 536 | 99.26 | |
| RNASEH1 | 2.05 | 0.40 | 0 | 0 | 5 | 0 | 5 | 244 | 99.19 | |
| BICDL2 | 1.97 | 0.32 | 5 | 0 | 10 | 2 | 12 | 507 | 100.00 | Chimpanzee |
| MVD | 1.97 | 0.42 | 1 | 0 | 7 | 0 | 6 | 356 | 93.93 | |
| IRF7 | 1.94 | 0.32 | 1 | 1 | 9 | 4 | 11 | 465 | 90.12 | |
| LAMP2 | 1.92 | 0.36 | 3 | 0 | 7 | 0 | 7 | 364 | 92.86 | |
| SGCB | 1.90 | 0.50 | 1 | 0 | 6 | 0 | 2 | 316 | 99.68 | |
| GABRD | 1.88 | 0.64 | 0 | 0 | 6 | 0 | 0 | 319 | 70.42 | Gibbon |
| DALRD3 | 1.86 | 0.45 | 1 | 0 | 7 | 1 | 4 | 376 | 100.00 | |
| SLC17A9 | 1.86 | 0.39 | 2 | 0 | 6 | 0 | 5 | 323 | 77.64 | |
| RNF39 | 1.85 | 0.42 | 2 | 0 | 7 | 0 | 5 | 379 | 100.00 | |
| METTL11B | 1.84 | 0.46 | 1 | 1 | 5 | 0 | 1 | 272 | 97.84 | Gibbon |
| HADH | 1.81 | 0.35 | 2 | 2 | 7 | 0 | 7 | 387 | 99.23 | |
| SNAPC2 | 1.80 | 0.37 | 2 | 1 | 6 | 0 | 5 | 334 | 100.00 | |
| CEP68 | 1.75 | 0.32 | 0 | 2 | 10 | 2 | 15 | 573 | 98.79 | Human |
| SLC15A3 | 1.71 | 0.35 | 2 | 2 | 7 | 1 | 6 | 409 | 81.64 | |
| CACUL1 | 1.70 | 0.50 | 1 | 1 | 6 | 0 | 1 | 352 | 95.65 | Gibbon |
| IGFN1 | 1.69 | 0.29 | 8 | 9 | 19 | 2 | 27 | 1126 | 90.51 | |
| SNX32 | 1.68 | 0.50 | 0 | 0 | 5 | 0 | 2 | 297 | 90.83 | |
| TMEM62 | 1.67 | 0.46 | 1 | 0 | 9 | 0 | 7 | 539 | 92.45 | Chimpanzee |
| EFCAB6 | 1.67 | 0.28 | 6 | 9 | 25 | 4 | 45 | 1501 | 100.00 | |
| PAICS | 1.66 | 0.45 | 3 | 2 | 7 | 0 | 1 | 421 | 98.14 | Gibbon |
| MEX3A | 1.65 | 0.57 | 0 | 0 | 7 | 0 | 2 | 424 | 94.64 | Gibbon |
| NRROS | 1.64 | 0.29 | 4 | 3 | 11 | 1 | 18 | 670 | 99.41 | |
| OTOP1 | 1.64 | 0.29 | 1 | 1 | 10 | 4 | 17 | 611 | 100.00 | |
| OTUD4 | 1.56 | 0.49 | 1 | 3 | 16 | 1 | 9 | 1027 | 97.07 | |
| SH2B2 | 1.52 | 0.36 | 1 | 0 | 7 | 0 | 9 | 460 | 80.99 | |
| ZCCHC8 | 1.46 | 0.44 | 0 | 2 | 9 | 2 | 5 | 617 | 91.54 | |
| RPS6KL1 | 1.46 | 0.35 | 0 | 2 | 8 | 2 | 9 | 549 | 100.00 | Human |
| ITIH6 | 1.45 | 0.32 | 5 | 4 | 15 | 0 | 21 | 1031 | 98.10 | |
| CLMN | 1.44 | 0.33 | 2 | 4 | 14 | 2 | 18 | 972 | 100.00 | |
| ECT2L | 1.44 | 0.35 | 3 | 1 | 13 | 2 | 16 | 903 | 100.00 | Chimpanzee |
| CAPN15 | 1.42 | 0.43 | 0 | 1 | 8 | 0 | 7 | 565 | 80.48 | Human |

| | | | | | | | | | | |
|---|---|---|---|---|---|---|---|---|---|---|
| NOX3 | 1.41 | 0.32 | 0 | 4 | 8 | 0 | 11 | 568 | 100.00 | Human |
| CWF19L2 | 1.41 | 0.34 | 5 | 4 | 12 | 0 | 12 | 854 | 98.96 | |
| IQCH | 1.37 | 0.31 | 5 | 5 | 14 | 1 | 19 | 1024 | 99.71 | |
| ZWILCH | 1.36 | 0.36 | 0 | 1 | 7 | 0 | 9 | 513 | 92.93 | Human |
| MAPKBP1 | 1.34 | 0.36 | 1 | 7 | 20 | 2 | 24 | 1497 | 98.88 | Human |
| RASA3 | 1.32 | 0.41 | 0 | 5 | 8 | 0 | 4 | 608 | 82.16 | |
| EVPL | 1.30 | 0.34 | 9 | 8 | 25 | 0 | 30 | 1920 | 97.96 | |
| CHD1L | 1.28 | 0.36 | 2 | 3 | 11 | 1 | 11 | 859 | 95.76 | |
| ITGA8 | 1.24 | 0.36 | 4 | 4 | 13 | 0 | 13 | 1045 | 99.71 | |
| KTN1 | 1.23 | 0.37 | 3 | 5 | 14 | 0 | 14 | 1136 | 93.57 | |
| ANO6 | 1.22 | 0.50 | 0 | 1 | 11 | 0 | 7 | 901 | 97.83 | Human |
| CFAP61 | 1.21 | 0.31 | 2 | 6 | 15 | 1 | 22 | 1236 | 99.92 | Human |
| PXDNL | 1.16 | 0.26 | 10 | 6 | 17 | 1 | 30 | 1463 | 100.00 | |
| CFAP65 | 1.14 | 0.25 | 14 | 10 | 21 | 1 | 36 | 1838 | 97.09 | |
| CEP192 | 1.10 | 0.26 | 14 | 11 | 28 | 6 | 49 | 2537 | 100.00 | |
| CUBN | 1.04 | 0.25 | 15 | 17 | 34 | 3 | 66 | 3280 | 95.99 | |

**S8 Table**: Higher than expected substitution rate on the gibbon branch

| Gene | Gibbon % subs per site | Gibbon norm branch length | Human subs | Chimp subs | Gorilla subs | #1# subs | Gibbon subs | Align overlap | Align Sat | Lower in |
|---|---|---|---|---|---|---|---|---|---|---|
| RBMXL3 | 11.97 | 0.64 | 9 | 19 | 18 | 9 | 99 | 827 | 91.89 | |
| MRPL4 | 10.80 | 0.70 | 1 | 0 | 4 | 0 | 19 | 176 | 61.54 | |
| MUC13 | 9.70 | 0.72 | 4 | 1 | 9 | 1 | 46 | 474 | 95.76 | |
| FAM71E2 | 9.48 | 0.68 | 13 | 8 | 14 | 1 | 79 | 833 | 95.42 | |
| CD58 | 8.91 | 0.71 | 1 | 1 | 4 | 0 | 22 | 247 | 98.80 | |
| BEND2 | 8.77 | 0.64 | 7 | 8 | 9 | 10 | 64 | 730 | 96.43 | |
| MISP | 8.45 | 0.72 | 7 | 4 | 6 | 2 | 56 | 663 | 98.51 | |
| FAM53A | 8.44 | 0.63 | 5 | 5 | 6 | 1 | 33 | 391 | 98.24 | |
| CD72 | 8.38 | 0.70 | 2 | 2 | 4 | 2 | 30 | 358 | 99.72 | |
| PODXL | 8.16 | 0.72 | 0 | 4 | 7 | 1 | 39 | 478 | 85.97 | Human |
| AC025287.4 | 8.14 | 0.66 | 3 | 4 | 6 | 0 | 31 | 381 | 100.00 | |
| TEX13A | 8.09 | 0.64 | 0 | 5 | 8 | 2 | 31 | 383 | 96.72 | |
| PLA2G4C | 8.04 | 0.78 | 1 | 1 | 5 | 0 | 37 | 460 | 88.63 | |
| PASD1 | 7.98 | 0.60 | 9 | 13 | 8 | 5 | 56 | 702 | 92.01 | |
| TAF7L | 7.88 | 0.68 | 4 | 4 | 2 | 2 | 32 | 406 | 95.75 | |
| C16orf71 | 7.84 | 0.67 | 5 | 3 | 9 | 0 | 40 | 510 | 98.27 | |
| CXorf66 | 7.80 | 0.64 | 1 | 6 | 5 | 1 | 28 | 359 | 99.72 | |
| C7orf61 | 7.73 | 0.62 | 0 | 1 | 4 | 1 | 15 | 194 | 97.49 | |
| MROH9 | 7.60 | 0.69 | 4 | 12 | 6 | 4 | 65 | 855 | 99.53 | |
| RIPK3 | 7.58 | 0.79 | 2 | 4 | 1 | 0 | 38 | 501 | 99.01 | |
| C4orf50 | 7.57 | 0.67 | 10 | 11 | 18 | 8 | 100 | 1321 | 94.02 | |
| ZBP1 | 7.51 | 0.68 | 2 | 5 | 5 | 0 | 32 | 426 | 99.30 | |
| RNF213 | 7.15 | 0.74 | 37 | 42 | 36 | 12 | 358 | 5006 | 95.28 | |
| C16orf46 | 7.14 | 0.64 | 5 | 2 | 4 | 1 | 27 | 378 | 97.42 | |
| GP1BA | 7.02 | 0.73 | 2 | 2 | 9 | 0 | 44 | 627 | 96.17 | |
| A3GALT2 | 6.97 | 0.62 | 3 | 2 | 5 | 1 | 23 | 330 | 97.06 | |
| CEP295NL | 6.96 | 0.63 | 0 | 5 | 9 | 7 | 41 | 589 | 98.66 | Human |
| DSPP | 6.89 | 0.58 | 7 | 20 | 19 | 6 | 73 | 1059 | 83.72 | |
| FRMD1 | 6.80 | 0.66 | 3 | 4 | 2 | 0 | 23 | 338 | 99.12 | |
| C2orf78 | 6.72 | 0.62 | 6 | 11 | 14 | 1 | 55 | 819 | 98.67 | |
| NLRP1 | 6.51 | 0.76 | 8 | 5 | 10 | 3 | 93 | 1428 | 97.14 | |
| ICOSLG | 6.37 | 0.61 | 2 | 3 | 4 | 3 | 23 | 361 | 87.83 | |
| ADAM12 | 6.29 | 0.78 | 3 | 5 | 2 | 3 | 57 | 906 | 99.67 | Gorilla |
| C19orf57 | 6.29 | 0.69 | 4 | 3 | 8 | 0 | 40 | 636 | 99.84 | |
| ZNF853 | 6.25 | 0.70 | 2 | 4 | 3 | 0 | 29 | 464 | 83.30 | |
| MUC1 | 6.17 | 0.70 | 0 | 3 | 6 | 0 | 29 | 470 | 97.31 | Human |
| C2orf81 | 6.15 | 0.65 | 0 | 4 | 9 | 2 | 33 | 537 | 99.63 | Human |
| MUM1 | 6.11 | 0.69 | 5 | 2 | 7 | 1 | 40 | 655 | 92.12 | |
| CEP72 | 6.10 | 0.65 | 7 | 1 | 7 | 2 | 37 | 607 | 99.51 | |
| PTPRH | 6.00 | 0.71 | 6 | 5 | 5 | 2 | 51 | 850 | 87.81 | |
| TESMIN | 5.96 | 0.66 | 3 | 4 | 3 | 0 | 26 | 436 | 92.37 | |
| C1orf127 | 5.95 | 0.61 | 5 | 8 | 9 | 0 | 39 | 655 | 84.41 | |
| NUTM1 | 5.89 | 0.61 | 3 | 4 | 8 | 0 | 28 | 475 | 99.37 | |
| ZNF114 | 5.85 | 0.64 | 3 | 4 | 2 | 1 | 23 | 393 | 98.99 | |
| SELPLG | 5.76 | 0.59 | 1 | 3 | 9 | 1 | 24 | 417 | 97.66 | |
| RP1L1 | 5.71 | 0.56 | 17 | 19 | 25 | 6 | 86 | 1507 | 79.61 | |
| JHY | 5.69 | 0.67 | 4 | 5 | 9 | 1 | 44 | 773 | 99.74 | |
| DTX3L | 5.68 | 0.77 | 1 | 3 | 5 | 0 | 42 | 740 | 100.00 | |
| ZNF683 | 5.63 | 0.68 | 2 | 2 | 5 | 1 | 28 | 497 | 96.69 | |
| SBSN | 5.62 | 0.68 | 1 | 0 | 5 | 2 | 24 | 427 | 77.08 | |
| CFAP157 | 5.62 | 0.67 | 4 | 2 | 3 | 0 | 25 | 445 | 99.78 | |
| MUC4 | 5.59 | 0.61 | 7 | 10 | 9 | 3 | 49 | 877 | 85.56 | |
| SPOCD1 | 5.56 | 0.64 | 10 | 6 | 15 | 4 | 66 | 1186 | 99.41 | |
| CYLC1 | 5.56 | 0.68 | 1 | 0 | 7 | 5 | 35 | 630 | 97.83 | Chimpanzee |
| ANKK1 | 5.55 | 0.57 | 9 | 8 | 4 | 2 | 34 | 613 | 88.97 | |
| ZNF557 | 5.52 | 0.67 | 1 | 0 | 4 | 0 | 17 | 308 | 83.70 | |
| TULP2 | 5.47 | 0.61 | 3 | 1 | 4 | 0 | 17 | 311 | 67.61 | |
| C12orf50 | 5.42 | 0.62 | 1 | 3 | 4 | 1 | 20 | 369 | 89.13 | |
| HSF5 | 5.40 | 0.68 | 0 | 3 | 7 | 2 | 32 | 593 | 99.50 | Human |
| ZNF473 | 5.40 | 0.69 | 4 | 8 | 5 | 1 | 47 | 871 | 100.00 | |
| CDHR5 | 5.38 | 0.58 | 9 | 7 | 5 | 3 | 37 | 688 | 98.43 | |
| POLRMT | 5.28 | 0.71 | 3 | 4 | 7 | 3 | 50 | 947 | 93.12 | |
| SPERT | 5.28 | 0.62 | 3 | 3 | 5 | 0 | 23 | 436 | 97.98 | |
| ERICH6B | 5.27 | 0.60 | 5 | 3 | 7 | 5 | 34 | 645 | 96.85 | |
| PABPC3 | 5.26 | 0.61 | 6 | 3 | 6 | 3 | 33 | 627 | 99.37 | |
| C1orf94 | 5.21 | 0.58 | 5 | 11 | 4 | 0 | 31 | 595 | 99.50 | |
| CFAP100 | 5.18 | 0.65 | 2 | 4 | 4 | 4 | 31 | 599 | 98.04 | |
| TEX28 | 5.12 | 0.67 | 0 | 1 | 5 | 1 | 21 | 410 | 100.00 | |
| MEFV | 5.12 | 0.61 | 5 | 7 | 5 | 6 | 40 | 781 | 100.00 | |
| UNC93A | 5.10 | 0.64 | 0 | 0 | 4 | 1 | 15 | 294 | 79.67 | |
| TFB2M | 5.10 | 0.70 | 1 | 0 | 4 | 0 | 20 | 392 | 99.75 | |
| CCDC129 | 5.07 | 0.56 | 6 | 13 | 14 | 4 | 51 | 1006 | 95.45 | |
| HJURP | 5.07 | 0.56 | 10 | 11 | 5 | 1 | 37 | 730 | 98.38 | |

| | | | | | | | | | | |
|---|---|---|---|---|---|---|---|---|---|---|
| MKI67 | 5.05 | 0.61 | 24 | 24 | 40 | 14 | 158 | 3127 | 96.13 | |
| CAGE1 | 5.01 | 0.65 | 1 | 6 | 13 | 0 | 42 | 839 | 100.00 | Human |
| SHROOM1 | 4.99 | 0.61 | 5 | 3 | 11 | 4 | 40 | 802 | 94.35 | |
| RHAG | 4.98 | 0.66 | 1 | 2 | 4 | 0 | 20 | 402 | 98.29 | |
| ADPRHL1 | 4.96 | 0.61 | 14 | 17 | 13 | 3 | 76 | 1532 | 91.57 | |
| SFI1 | 4.95 | 0.62 | 6 | 7 | 12 | 5 | 53 | 1071 | 92.33 | |
| CTU2 | 4.95 | 0.63 | 2 | 4 | 2 | 3 | 24 | 485 | 94.36 | |
| C1orf87 | 4.95 | 0.72 | 2 | 0 | 5 | 0 | 27 | 546 | 100.00 | Chimpanzee |
| CCDC15 | 4.94 | 0.74 | 4 | 2 | 3 | 3 | 44 | 890 | 95.19 | |
| TTC6 | 4.94 | 0.63 | 16 | 15 | 10 | 1 | 75 | 1518 | 92.50 | |
| LRRC66 | 4.89 | 0.60 | 6 | 3 | 15 | 2 | 43 | 880 | 100.00 | |
| PARP9 | 4.88 | 0.65 | 3 | 7 | 8 | 1 | 40 | 819 | 100.00 | |
| CCNB3 | 4.88 | 0.66 | 6 | 6 | 16 | 4 | 68 | 1393 | 100.00 | |
| AC099489.1 | 4.88 | 0.62 | 27 | 25 | 23 | 8 | 137 | 2810 | 88.20 | |
| PLEKHG4B | 4.85 | 0.62 | 9 | 13 | 15 | 6 | 75 | 1547 | 98.54 | |
| HTRA4 | 4.83 | 0.65 | 1 | 2 | 4 | 2 | 23 | 476 | 100.00 | |
| KIAA0391 | 4.83 | 0.68 | 3 | 1 | 4 | 2 | 28 | 580 | 99.49 | |
| ZAN | 4.83 | 0.52 | 23 | 16 | 35 | 10 | 94 | 1948 | 81.54 | |
| ZFR2 | 4.82 | 0.56 | 3 | 11 | 13 | 5 | 44 | 913 | 98.28 | |
| PMFBP1 | 4.80 | 0.66 | 5 | 4 | 7 | 3 | 43 | 895 | 90.68 | |
| FBXW12 | 4.79 | 0.63 | 0 | 2 | 7 | 0 | 21 | 438 | 94.60 | |
| ZNF831 | 4.78 | 0.60 | 9 | 9 | 30 | 4 | 80 | 1674 | 99.94 | |
| ZBBX | 4.76 | 0.55 | 11 | 6 | 11 | 0 | 38 | 799 | 99.88 | |
| CCDC33 | 4.72 | 0.57 | 7 | 9 | 5 | 0 | 31 | 657 | 91.50 | |
| CD5 | 4.70 | 0.67 | 1 | 1 | 4 | 2 | 23 | 489 | 98.79 | |
| TOGARAM2 | 4.67 | 0.54 | 18 | 7 | 10 | 1 | 46 | 984 | 97.23 | |
| OVCH1 | 4.67 | 0.64 | 4 | 7 | 12 | 2 | 49 | 1049 | 96.33 | |
| CCDC110 | 4.62 | 0.59 | 8 | 6 | 6 | 1 | 35 | 757 | 94.39 | |
| ZMYND15 | 4.60 | 0.59 | 2 | 3 | 12 | 3 | 33 | 718 | 97.29 | |
| C19orf44 | 4.59 | 0.58 | 2 | 2 | 10 | 5 | 30 | 653 | 99.39 | |
| CCDC116 | 4.59 | 0.57 | 5 | 6 | 7 | 0 | 28 | 610 | 99.51 | |
| BRCA2 | 4.57 | 0.78 | 12 | 12 | 17 | 1 | 156 | 3411 | 99.85 | |
| C3orf20 | 4.57 | 0.65 | 1 | 3 | 11 | 3 | 39 | 853 | 99.53 | Human |
| SLC9C1 | 4.57 | 0.63 | 12 | 6 | 8 | 3 | 53 | 1160 | 98.98 | |
| C2orf16 | 4.57 | 0.62 | 43 | 37 | 43 | 17 | 228 | 4994 | 94.08 | |
| SYCP2L | 4.56 | 0.65 | 3 | 3 | 9 | 2 | 37 | 812 | 100.00 | |
| RAB44 | 4.53 | 0.69 | 4 | 4 | 6 | 1 | 41 | 905 | 97.42 | |
| RECQL4 | 4.52 | 0.58 | 8 | 5 | 10 | 9 | 47 | 1039 | 91.38 | |
| PPP1R15A | 4.52 | 0.62 | 7 | 3 | 4 | 1 | 30 | 664 | 98.52 | |
| AJM1 | 4.50 | 0.66 | 2 | 1 | 6 | 1 | 26 | 578 | 85.50 | |
| RAD51AP2 | 4.49 | 0.55 | 7 | 14 | 15 | 4 | 52 | 1158 | 100.00 | |
| LEKR1 | 4.48 | 0.75 | 0 | 1 | 6 | 0 | 31 | 692 | 100.00 | Human |
| ALPK2 | 4.47 | 0.61 | 15 | 16 | 14 | 13 | 93 | 2079 | 99.71 | |
| RIPK1 | 4.47 | 0.73 | 0 | 4 | 2 | 2 | 30 | 671 | 100.00 | Human |
| UMODL1 | 4.47 | 0.57 | 20 | 13 | 8 | 2 | 61 | 1365 | 95.19 | |
| SH3D21 | 4.41 | 0.58 | 7 | 6 | 5 | 2 | 31 | 703 | 98.18 | |
| TMEM44 | 4.39 | 0.60 | 2 | 5 | 1 | 0 | 17 | 387 | 89.38 | |
| KIAA1210 | 4.39 | 0.60 | 8 | 7 | 22 | 7 | 70 | 1594 | 96.14 | |
| CROCC2 | 4.33 | 0.53 | 10 | 12 | 25 | 7 | 64 | 1479 | 94.14 | |
| PTPRC | 4.32 | 0.77 | 2 | 5 | 3 | 3 | 55 | 1273 | 97.62 | |
| R3HCC1L | 4.30 | 0.56 | 10 | 5 | 8 | 1 | 34 | 790 | 99.75 | |
| KIAA1614 | 4.30 | 0.55 | 13 | 7 | 15 | 1 | 47 | 1093 | 91.85 | |
| CCDC40 | 4.30 | 0.62 | 7 | 9 | 10 | 0 | 48 | 1117 | 98.24 | |
| FAM83C | 4.29 | 0.58 | 4 | 3 | 10 | 3 | 32 | 746 | 100.00 | |
| R3HCC1 | 4.29 | 0.61 | 1 | 2 | 5 | 1 | 19 | 443 | 95.27 | |
| DNAI1 | 4.28 | 0.70 | 2 | 4 | 3 | 0 | 29 | 677 | 96.85 | |
| CATSPERE | 4.28 | 0.61 | 3 | 11 | 4 | 4 | 39 | 911 | 97.75 | |
| LRRC53 | 4.27 | 0.64 | 13 | 5 | 7 | 2 | 53 | 1240 | 99.60 | |
| NINL | 4.27 | 0.58 | 10 | 9 | 17 | 1 | 54 | 1264 | 92.06 | |
| ZNF195 | 4.27 | 0.72 | 5 | 1 | 1 | 0 | 27 | 633 | 100.00 | |
| AKAP4 | 4.26 | 0.65 | 2 | 3 | 8 | 3 | 36 | 845 | 100.00 | |
| ENAM | 4.26 | 0.65 | 6 | 7 | 5 | 1 | 41 | 963 | 99.90 | |
| PARP4 | 4.25 | 0.62 | 8 | 8 | 14 | 3 | 59 | 1387 | 92.28 | |
| STARD9 | 4.25 | 0.60 | 27 | 26 | 59 | 18 | 194 | 4564 | 99.33 | |
| SCARF1 | 4.23 | 0.65 | 3 | 4 | 6 | 1 | 32 | 756 | 95.09 | |
| PTCD3 | 4.23 | 0.64 | 0 | 5 | 6 | 2 | 29 | 686 | 99.56 | Human |
| RCSD1 | 4.21 | 0.67 | 0 | 1 | 4 | 0 | 17 | 404 | 97.82 | |
| TACC3 | 4.18 | 0.61 | 5 | 2 | 10 | 1 | 33 | 789 | 97.77 | |
| IQCC | 4.13 | 0.62 | 2 | 2 | 4 | 1 | 20 | 484 | 98.78 | |
| CDCA2 | 4.12 | 0.61 | 6 | 6 | 7 | 4 | 40 | 971 | 99.08 | |
| COL18A1 | 4.11 | 0.64 | 11 | 6 | 7 | 1 | 49 | 1192 | 83.01 | |
| PKD1L3 | 4.11 | 0.55 | 17 | 11 | 16 | 11 | 69 | 1679 | 98.76 | |
| KIAA1551 | 4.09 | 0.64 | 10 | 7 | 19 | 2 | 71 | 1734 | 99.31 | |
| CSF2RB | 4.09 | 0.59 | 5 | 3 | 10 | 4 | 36 | 881 | 97.56 | |
| MDC1 | 4.08 | 0.62 | 7 | 16 | 21 | 3 | 80 | 1961 | 95.85 | |
| ERICH3 | 4.08 | 0.57 | 15 | 10 | 18 | 2 | 62 | 1521 | 99.93 | |
| MARCO | 4.05 | 0.60 | 5 | 2 | 3 | 1 | 21 | 518 | 99.62 | |
| March10 | 4.05 | 0.61 | 7 | 2 | 7 | 1 | 31 | 765 | 99.61 | |

| | | | | | | | | | | |
|---|---|---|---|---|---|---|---|---|---|---|
| EFCC1 | 4.04 | 0.60 | 4 | 2 | 3 | 0 | 18 | 445 | 89.54 | |
| LRRIQ1 | 4.03 | 0.61 | 9 | 7 | 21 | 4 | 68 | 1688 | 99.76 | |
| TROAP | 4.02 | 0.71 | 3 | 0 | 4 | 2 | 31 | 771 | 97.10 | Chimpanzee |
| L1TD1 | 4.02 | 0.58 | 5 | 7 | 5 | 5 | 34 | 846 | 97.92 | |
| KIAA1755 | 4.01 | 0.65 | 7 | 6 | 7 | 1 | 45 | 1121 | 94.52 | |
| GGT6 | 4.01 | 0.62 | 2 | 2 | 4 | 1 | 20 | 499 | 100.00 | |
| CLEC20A | 4.00 | 0.61 | 3 | 0 | 4 | 0 | 16 | 400 | 100.00 | |
| SLC9C2 | 3.99 | 0.68 | 3 | 5 | 9 | 1 | 44 | 1103 | 99.73 | |
| MYO15B | 3.98 | 0.56 | 17 | 15 | 34 | 6 | 96 | 2411 | 92.41 | |
| ADCY10 | 3.97 | 0.63 | 7 | 9 | 14 | 3 | 60 | 1510 | 99.47 | |
| TEX11 | 3.97 | 0.57 | 1 | 10 | 11 | 2 | 36 | 906 | 99.34 | Human |
| MORC1 | 3.97 | 0.63 | 5 | 7 | 6 | 2 | 39 | 982 | 99.80 | |
| MRPS5 | 3.96 | 0.60 | 1 | 0 | 5 | 2 | 17 | 429 | 99.77 | |
| BDP1 | 3.93 | 0.65 | 16 | 16 | 18 | 0 | 97 | 2468 | 95.44 | |
| MAMDC4 | 3.92 | 0.63 | 6 | 6 | 9 | 4 | 47 | 1199 | 99.67 | |
| AHNAK2 | 3.91 | 0.64 | 6 | 3 | 5 | 0 | 31 | 793 | 100.00 | |
| C10orf90 | 3.91 | 0.59 | 4 | 5 | 6 | 1 | 27 | 691 | 100.00 | |
| LRMP | 3.91 | 0.62 | 9 | 9 | 10 | 3 | 56 | 1434 | 99.93 | |
| WDR60 | 3.89 | 0.60 | 3 | 5 | 9 | 3 | 35 | 899 | 95.33 | |
| IRAK1 | 3.88 | 0.62 | 3 | 2 | 4 | 0 | 20 | 515 | 76.52 | |
| NOL8 | 3.88 | 0.60 | 7 | 7 | 8 | 1 | 39 | 1006 | 93.06 | |
| WDR97 | 3.88 | 0.53 | 18 | 12 | 18 | 2 | 59 | 1522 | 97.25 | |
| DENND3 | 3.87 | 0.56 | 9 | 11 | 10 | 4 | 47 | 1215 | 98.46 | |
| PKD1L1 | 3.86 | 0.58 | 19 | 27 | 22 | 7 | 105 | 2719 | 96.35 | |
| CCDC57 | 3.85 | 0.60 | 8 | 4 | 4 | 2 | 31 | 805 | 97.58 | |
| SGO1 | 3.85 | 0.60 | 5 | 3 | 2 | 0 | 20 | 520 | 98.67 | |
| LINS1 | 3.85 | 0.58 | 6 | 2 | 8 | 2 | 29 | 754 | 99.60 | |
| CEP164 | 3.84 | 0.75 | 6 | 3 | 4 | 2 | 54 | 1405 | 98.25 | |
| KANK4 | 3.82 | 0.58 | 8 | 6 | 11 | 0 | 38 | 995 | 100.00 | |
| CASS4 | 3.82 | 0.60 | 4 | 10 | 3 | 0 | 30 | 786 | 100.00 | |
| KIAA1211L | 3.81 | 0.66 | 4 | 1 | 5 | 3 | 31 | 814 | 96.67 | Chimpanzee |
| GPR179 | 3.79 | 0.67 | 7 | 11 | 21 | 3 | 89 | 2347 | 99.75 | Human |
| FANCA | 3.79 | 0.55 | 9 | 9 | 12 | 9 | 50 | 1319 | 93.15 | |
| PPP1R15B | 3.79 | 0.59 | 6 | 3 | 5 | 2 | 27 | 713 | 100.00 | |
| TSEN2 | 3.79 | 0.60 | 1 | 2 | 5 | 0 | 17 | 449 | 98.90 | |
| NWD1 | 3.78 | 0.59 | 13 | 9 | 12 | 2 | 55 | 1454 | 94.42 | |
| FAM208B | 3.78 | 0.58 | 29 | 19 | 14 | 0 | 90 | 2382 | 98.96 | |
| RAB11FIP1 | 3.77 | 0.60 | 8 | 4 | 10 | 0 | 38 | 1008 | 86.97 | |
| ZNF408 | 3.76 | 0.67 | 3 | 2 | 5 | 0 | 27 | 718 | 100.00 | |
| C10orf71 | 3.75 | 0.64 | 12 | 2 | 7 | 3 | 47 | 1252 | 97.13 | Chimpanzee |
| ALMS1 | 3.72 | 0.58 | 31 | 36 | 33 | 3 | 147 | 3947 | 97.41 | |
| MICALL2 | 3.71 | 0.58 | 7 | 6 | 7 | 0 | 32 | 862 | 96.96 | |
| CENPC | 3.71 | 0.67 | 1 | 5 | 7 | 1 | 35 | 943 | 100.00 | Human |
| CEP295 | 3.71 | 0.58 | 19 | 15 | 26 | 7 | 95 | 2561 | 98.69 | |
| PTPRJ | 3.68 | 0.67 | 4 | 5 | 8 | 2 | 45 | 1222 | 95.32 | |
| ADAD2 | 3.68 | 0.65 | 3 | 4 | 1 | 0 | 21 | 571 | 100.00 | |
| CRB2 | 3.67 | 0.66 | 5 | 6 | 2 | 1 | 34 | 926 | 87.28 | Gorilla |
| HASPIN | 3.67 | 0.63 | 2 | 7 | 5 | 0 | 29 | 790 | 100.00 | |
| SLC6A16 | 3.67 | 0.63 | 4 | 4 | 4 | 1 | 27 | 736 | 100.00 | |
| MEIOC | 3.66 | 0.64 | 5 | 4 | 6 | 1 | 34 | 929 | 99.68 | |
| RNASEL | 3.66 | 0.61 | 2 | 9 | 3 | 0 | 27 | 738 | 99.60 | |
| ENTHD1 | 3.64 | 0.59 | 2 | 3 | 5 | 2 | 22 | 604 | 99.67 | |
| CCDC175 | 3.64 | 0.61 | 6 | 6 | 3 | 0 | 28 | 770 | 97.10 | |
| DRC1 | 3.62 | 0.67 | 4 | 1 | 4 | 0 | 25 | 690 | 99.42 | |
| ABCC3 | 3.61 | 0.69 | 6 | 4 | 5 | 0 | 40 | 1109 | 84.53 | |
| FBF1 | 3.60 | 0.68 | 5 | 3 | 1 | 1 | 28 | 777 | 83.01 | Gorilla |
| KIAA0408 | 3.60 | 0.67 | 5 | 2 | 1 | 1 | 25 | 694 | 100.00 | Gorilla |
| SOX30 | 3.60 | 0.61 | 3 | 2 | 6 | 3 | 27 | 750 | 99.60 | |
| CCDC158 | 3.60 | 0.76 | 2 | 2 | 5 | 0 | 40 | 1112 | 100.00 | |
| JMJD4 | 3.59 | 0.63 | 4 | 1 | 1 | 0 | 16 | 446 | 99.78 | |
| PPP1R3A | 3.57 | 0.60 | 7 | 4 | 7 | 6 | 40 | 1120 | 100.00 | |
| SGO2 | 3.57 | 0.64 | 5 | 6 | 8 | 2 | 42 | 1176 | 99.32 | |
| PARP10 | 3.55 | 0.56 | 8 | 10 | 5 | 3 | 36 | 1014 | 98.93 | |
| CRYBG3 | 3.55 | 0.58 | 23 | 12 | 27 | 10 | 102 | 2873 | 98.22 | |
| MEGF6 | 3.54 | 0.57 | 15 | 5 | 5 | 3 | 41 | 1159 | 83.38 | |
| CCDC180 | 3.54 | 0.58 | 10 | 9 | 15 | 4 | 57 | 1612 | 96.93 | |
| ZNF519 | 3.53 | 0.61 | 4 | 1 | 1 | 3 | 19 | 539 | 100.00 | |
| CARD14 | 3.50 | 0.61 | 2 | 6 | 8 | 0 | 30 | 856 | 91.94 | |
| CHFR | 3.50 | 0.63 | 0 | 2 | 5 | 1 | 19 | 543 | 85.92 | Human |
| RTN3 | 3.50 | 0.64 | 4 | 5 | 7 | 1 | 36 | 1029 | 99.71 | |
| NLRP12 | 3.49 | 0.64 | 5 | 7 | 6 | 0 | 37 | 1059 | 99.91 | |
| WDR90 | 3.49 | 0.61 | 8 | 5 | 15 | 3 | 53 | 1518 | 94.64 | |
| NEK4 | 3.48 | 0.65 | 4 | 3 | 4 | 0 | 27 | 775 | 95.68 | |
| ERVMER34-1 | 3.48 | 0.65 | 1 | 4 | 2 | 0 | 19 | 546 | 100.00 | |
| TTC31 | 3.47 | 0.64 | 4 | 2 | 1 | 0 | 18 | 518 | 99.81 | |
| CIITA | 3.47 | 0.60 | 6 | 3 | 11 | 2 | 38 | 1096 | 96.91 | |
| OBSCN | 3.45 | 0.54 | 33 | 49 | 92 | 37 | 250 | 7241 | 89.73 | |
| CC2D2B | 3.45 | 0.62 | 2 | 6 | 5 | 6 | 36 | 1043 | 98.58 | Human |

| | | | | | | | | | | |
|---|---|---|---|---|---|---|---|---|---|---|
| ZNF205 | 3.44 | 0.61 | 3 | 2 | 4 | 0 | 19 | 552 | 100.00 | |
| RGSL1 | 3.44 | 0.61 | 6 | 5 | 7 | 3 | 37 | 1075 | 99.91 | |
| AKAP1 | 3.44 | 0.56 | 5 | 8 | 6 | 2 | 31 | 901 | 99.89 | |
| FHAD1 | 3.44 | 0.59 | 5 | 7 | 10 | 4 | 42 | 1221 | 91.05 | |
| PLB1 | 3.43 | 0.69 | 1 | 3 | 10 | 4 | 47 | 1369 | 96.54 | |
| TDRD12 | 3.41 | 0.60 | 5 | 7 | 4 | 5 | 36 | 1057 | 92.64 | |
| IFT81 | 3.40 | 0.73 | 0 | 1 | 4 | 0 | 23 | 676 | 100.00 | Human |
| TEX15 | 3.40 | 0.57 | 23 | 12 | 33 | 9 | 107 | 3147 | 99.37 | |
| ELAC2 | 3.39 | 0.65 | 5 | 2 | 4 | 0 | 27 | 797 | 98.03 | |
| DHX58 | 3.38 | 0.59 | 4 | 2 | 4 | 1 | 20 | 591 | 91.91 | |
| CCDC142 | 3.38 | 0.64 | 1 | 4 | 4 | 2 | 25 | 739 | 99.46 | |
| TDRD6 | 3.38 | 0.65 | 5 | 5 | 20 | 4 | 70 | 2071 | 98.90 | |
| ALPK3 | 3.37 | 0.60 | 9 | 7 | 15 | 2 | 53 | 1572 | 96.09 | |
| BTBD18 | 3.37 | 0.64 | 4 | 1 | 3 | 2 | 24 | 712 | 100.00 | |
| TDRD15 | 3.37 | 0.67 | 11 | 8 | 4 | 1 | 54 | 1604 | 99.75 | Gorilla |
| DCLRE1A | 3.37 | 0.58 | 5 | 7 | 9 | 1 | 35 | 1040 | 100.00 | |
| FSIP2 | 3.36 | 0.59 | 36 | 36 | 58 | 20 | 219 | 6513 | 97.79 | |
| GNAS | 3.35 | 0.68 | 2 | 2 | 6 | 2 | 33 | 984 | 99.90 | |
| SOWAHB | 3.35 | 0.61 | 4 | 3 | 5 | 0 | 24 | 716 | 95.34 | |
| SETX | 3.35 | 0.61 | 7 | 23 | 20 | 1 | 84 | 2511 | 97.48 | Human |
| TMEM8A | 3.34 | 0.64 | 2 | 2 | 4 | 1 | 22 | 658 | 96.06 | |
| KANK3 | 3.33 | 0.57 | 4 | 6 | 6 | 2 | 28 | 840 | 100.00 | |
| AKNA | 3.32 | 0.59 | 7 | 7 | 11 | 2 | 43 | 1295 | 92.83 | |
| KIAA1671 | 3.31 | 0.61 | 8 | 10 | 11 | 0 | 49 | 1479 | 91.86 | |
| BHMG1 | 3.31 | 0.61 | 2 | 4 | 4 | 0 | 21 | 634 | 99.37 | |
| PRR14L | 3.31 | 0.58 | 12 | 11 | 16 | 9 | 71 | 2144 | 99.72 | |
| CRYBG2 | 3.31 | 0.59 | 6 | 4 | 14 | 4 | 45 | 1360 | 99.85 | |
| ZNF541 | 3.31 | 0.68 | 4 | 4 | 5 | 1 | 36 | 1089 | 95.28 | |
| USPL1 | 3.30 | 0.68 | 9 | 3 | 2 | 0 | 36 | 1091 | 99.91 | Gorilla |
| BRDT | 3.28 | 0.59 | 4 | 6 | 8 | 1 | 31 | 944 | 99.89 | |
| VWA3A | 3.28 | 0.60 | 4 | 9 | 7 | 2 | 38 | 1159 | 97.89 | |
| TBCD | 3.28 | 0.73 | 6 | 1 | 2 | 0 | 34 | 1037 | 88.48 | |
| MIS18BP1 | 3.27 | 0.57 | 9 | 5 | 8 | 3 | 37 | 1132 | 100.00 | |
| ADGRG4 | 3.27 | 0.57 | 18 | 15 | 28 | 12 | 99 | 3031 | 99.38 | |
| EXPH5 | 3.27 | 0.59 | 8 | 15 | 13 | 5 | 64 | 1960 | 98.94 | |
| KIF24 | 3.26 | 0.70 | 5 | 3 | 8 | 0 | 44 | 1348 | 98.54 | Chimpanzee |
| ADAMTS13 | 3.25 | 0.56 | 6 | 7 | 8 | 8 | 41 | 1260 | 92.72 | |
| DLEC1 | 3.25 | 0.59 | 11 | 7 | 10 | 2 | 47 | 1447 | 94.14 | |
| MYCBPAP | 3.24 | 0.62 | 6 | 2 | 5 | 2 | 29 | 896 | 94.42 | |
| FASTKD1 | 3.21 | 0.66 | 2 | 1 | 4 | 3 | 26 | 810 | 100.00 | Chimpanzee |
| CDK5RAP2 | 3.21 | 0.65 | 7 | 7 | 13 | 1 | 58 | 1809 | 95.56 | |
| CEP89 | 3.20 | 0.60 | 4 | 3 | 5 | 0 | 23 | 718 | 99.86 | |
| WDR87 | 3.20 | 0.56 | 16 | 19 | 21 | 14 | 92 | 2879 | 99.14 | |
| TAS1R1 | 3.19 | 0.62 | 3 | 5 | 3 | 0 | 23 | 722 | 94.26 | |
| CMYA5 | 3.16 | 0.57 | 21 | 24 | 34 | 10 | 123 | 3892 | 97.74 | |
| TCOF1 | 3.15 | 0.61 | 6 | 12 | 5 | 0 | 41 | 1301 | 89.85 | |
| HAUS6 | 3.14 | 0.57 | 4 | 7 | 7 | 2 | 30 | 954 | 100.00 | |
| MIA3 | 3.14 | 0.63 | 11 | 13 | 6 | 2 | 59 | 1880 | 98.58 | Gorilla |
| ADGRG7 | 3.14 | 0.67 | 2 | 0 | 3 | 4 | 25 | 797 | 100.00 | Chimpanzee |
| SLC26A11 | 3.14 | 0.65 | 1 | 4 | 2 | 0 | 19 | 606 | 100.00 | |
| KCTD19 | 3.13 | 0.65 | 2 | 0 | 5 | 2 | 23 | 734 | 95.82 | Chimpanzee |
| PALB2 | 3.12 | 0.57 | 6 | 5 | 12 | 2 | 37 | 1186 | 100.00 | |
| ABCA13 | 3.11 | 0.55 | 40 | 33 | 40 | 10 | 152 | 4882 | 98.65 | |
| SHCBP1L | 3.11 | 0.68 | 6 | 0 | 0 | 0 | 20 | 643 | 100.00 | |
| HELZ2 | 3.11 | 0.63 | 11 | 13 | 13 | 7 | 79 | 2542 | 97.32 | |
| KNL1 | 3.10 | 0.58 | 12 | 18 | 16 | 3 | 71 | 2294 | 98.41 | |
| TMEM108 | 3.09 | 0.61 | 3 | 0 | 4 | 0 | 16 | 518 | 100.00 | Chimpanzee |
| LRGUK | 3.09 | 0.66 | 1 | 1 | 4 | 0 | 18 | 583 | 73.15 | |
| B4GALNT3 | 3.08 | 0.68 | 2 | 2 | 4 | 1 | 26 | 844 | 89.50 | |
| FANCM | 3.08 | 0.62 | 11 | 10 | 15 | 0 | 63 | 2046 | 99.90 | |
| SYNM | 3.07 | 0.60 | 7 | 12 | 7 | 1 | 45 | 1465 | 98.59 | |
| CCDC14 | 3.07 | 0.64 | 2 | 5 | 3 | 0 | 24 | 783 | 86.81 | |
| NSUN7 | 3.06 | 0.59 | 5 | 3 | 4 | 0 | 22 | 718 | 100.00 | |
| CCIN | 3.06 | 0.64 | 1 | 1 | 4 | 1 | 18 | 588 | 100.00 | |
| SEC16A | 3.06 | 0.57 | 18 | 13 | 15 | 1 | 65 | 2127 | 94.24 | |
| VWCE | 3.06 | 0.60 | 4 | 3 | 5 | 2 | 26 | 851 | 97.70 | |
| COL20A1 | 3.05 | 0.58 | 7 | 12 | 6 | 0 | 39 | 1277 | 99.22 | |
| SEL1L2 | 3.05 | 0.61 | 0 | 5 | 4 | 1 | 21 | 688 | 100.00 | Human |
| MYLK3 | 3.05 | 0.62 | 2 | 1 | 6 | 0 | 20 | 656 | 100.00 | |
| PCSK9 | 3.04 | 0.60 | 3 | 3 | 5 | 0 | 21 | 690 | 100.00 | |
| EHBP1L1 | 3.04 | 0.56 | 9 | 9 | 9 | 4 | 43 | 1414 | 95.54 | |
| POLQ | 3.02 | 0.61 | 10 | 3 | 18 | 9 | 67 | 2215 | 90.19 | Chimpanzee |
| SYTL2 | 3.02 | 0.58 | 12 | 13 | 13 | 8 | 67 | 2219 | 99.95 | |
| PKDREJ | 3.02 | 0.60 | 8 | 11 | 18 | 2 | 64 | 2120 | 98.15 | |
| QSOX1 | 3.02 | 0.61 | 6 | 3 | 2 | 0 | 22 | 729 | 98.65 | |
| RRP1B | 3.02 | 0.66 | 0 | 5 | 1 | 2 | 22 | 729 | 100.00 | |
| LCA5 | 3.01 | 0.65 | 4 | 1 | 2 | 1 | 21 | 697 | 100.00 | |
| CTDP1 | 3.01 | 0.62 | 3 | 3 | 6 | 2 | 28 | 930 | 97.79 | |

| | | | | | | | | | | |
|---|---|---|---|---|---|---|---|---|---|---|
| UNC13B | 3.01 | 0.56 | 20 | 19 | 38 | 19 | 123 | 4091 | 96.99 | |
| ESCO2 | 3.00 | 0.60 | 2 | 2 | 5 | 0 | 18 | 600 | 100.00 | |
| CCDC39 | 3.00 | 0.66 | 1 | 4 | 5 | 0 | 26 | 867 | 100.00 | Human |
| NEFH | 3.00 | 0.62 | 1 | 4 | 4 | 3 | 25 | 834 | 96.08 | Human |
| LAMC3 | 2.99 | 0.57 | 10 | 4 | 17 | 1 | 46 | 1536 | 99.10 | |
| AKAP12 | 2.99 | 0.60 | 8 | 9 | 13 | 3 | 53 | 1771 | 99.55 | |
| CSF3R | 2.99 | 0.58 | 6 | 3 | 6 | 0 | 25 | 836 | 100.00 | |
| TONSL | 2.98 | 0.61 | 5 | 5 | 11 | 2 | 40 | 1341 | 97.31 | |
| AOAH | 2.98 | 0.60 | 4 | 4 | 1 | 1 | 20 | 671 | 98.24 | Gorilla |
| ALPK1 | 2.98 | 0.56 | 7 | 9 | 7 | 3 | 36 | 1208 | 97.34 | |
| SIGLEC1 | 2.96 | 0.62 | 10 | 5 | 8 | 1 | 44 | 1485 | 92.75 | |
| EYS | 2.96 | 0.55 | 19 | 21 | 25 | 8 | 93 | 3142 | 99.94 | |
| CEP126 | 2.95 | 0.56 | 3 | 9 | 9 | 2 | 33 | 1117 | 100.00 | |
| FGD3 | 2.95 | 0.60 | 2 | 0 | 5 | 3 | 20 | 678 | 100.00 | Chimpanzee |
| WDPCP | 2.95 | 0.66 | 2 | 1 | 5 | 0 | 22 | 746 | 100.00 | |
| MYBBP1A | 2.95 | 0.62 | 4 | 5 | 5 | 2 | 31 | 1052 | 94.43 | |
| SPAG17 | 2.94 | 0.58 | 11 | 14 | 15 | 3 | 63 | 2141 | 99.35 | |
| STOX1 | 2.94 | 0.58 | 3 | 4 | 7 | 2 | 26 | 885 | 99.77 | |
| ANPEP | 2.94 | 0.59 | 5 | 2 | 7 | 0 | 25 | 851 | 99.77 | |
| THADA | 2.93 | 0.67 | 4 | 8 | 9 | 1 | 52 | 1772 | 94.56 | Human |
| COBLL1 | 2.93 | 0.66 | 5 | 5 | 4 | 1 | 36 | 1230 | 99.76 | |
| SEPT4 | 2.93 | 0.60 | 8 | 3 | 2 | 3 | 29 | 991 | 99.50 | Gorilla |
| BFSP1 | 2.92 | 0.63 | 1 | 0 | 5 | 2 | 19 | 650 | 99.54 | Chimpanzee |
| KIAA0586 | 2.91 | 0.57 | 5 | 7 | 13 | 6 | 44 | 1513 | 95.46 | |
| ADAMTS20 | 2.90 | 0.61 | 9 | 7 | 10 | 5 | 52 | 1793 | 98.09 | |
| COL11A2 | 2.90 | 0.76 | 1 | 2 | 4 | 0 | 34 | 1173 | 87.67 | |
| NLRC5 | 2.90 | 0.60 | 9 | 7 | 10 | 2 | 47 | 1622 | 90.97 | |
| ZKSCAN2 | 2.90 | 0.65 | 2 | 2 | 7 | 1 | 28 | 967 | 100.00 | |
| PLBD1 | 2.89 | 0.66 | 4 | 0 | 1 | 0 | 16 | 553 | 100.00 | Chimpanzee |
| CC2D1A | 2.89 | 0.60 | 6 | 2 | 5 | 0 | 24 | 830 | 92.02 | |
| CHTF18 | 2.89 | 0.64 | 3 | 1 | 4 | 0 | 20 | 692 | 74.89 | |
| PTPN22 | 2.89 | 0.62 | 2 | 1 | 4 | 2 | 20 | 693 | 100.00 | |
| CEP250 | 2.88 | 0.71 | 4 | 6 | 10 | 5 | 69 | 2395 | 98.89 | |
| ZGRF1 | 2.88 | 0.60 | 8 | 8 | 15 | 0 | 51 | 1772 | 96.67 | |
| HEG1 | 2.88 | 0.62 | 5 | 4 | 7 | 2 | 34 | 1182 | 98.99 | |
| C5orf42 | 2.87 | 0.56 | 13 | 17 | 27 | 10 | 87 | 3028 | 98.12 | |
| COL24A1 | 2.87 | 0.58 | 10 | 7 | 11 | 4 | 49 | 1706 | 99.82 | |
| ACSL5 | 2.84 | 0.60 | 4 | 2 | 4 | 1 | 21 | 739 | 100.00 | |
| TOPAZ1 | 2.83 | 0.60 | 9 | 7 | 12 | 0 | 47 | 1658 | 99.64 | |
| MMP17 | 2.83 | 0.60 | 2 | 2 | 4 | 0 | 17 | 600 | 99.50 | |
| ATF7IP2 | 2.83 | 0.65 | 0 | 3 | 4 | 0 | 19 | 671 | 99.70 | Human |
| EXO1 | 2.83 | 0.63 | 7 | 1 | 2 | 0 | 23 | 813 | 96.79 | Chimpanzee |
| EFHC1 | 2.81 | 0.64 | 2 | 0 | 4 | 1 | 18 | 641 | 100.00 | Chimpanzee |
| CCDC171 | 2.81 | 0.59 | 8 | 7 | 6 | 2 | 37 | 1318 | 99.40 | |
| DGKK | 2.80 | 0.66 | 2 | 2 | 6 | 3 | 31 | 1109 | 88.44 | |
| CORIN | 2.78 | 0.63 | 7 | 1 | 6 | 0 | 29 | 1042 | 100.00 | Chimpanzee |
| MMP9 | 2.78 | 0.65 | 1 | 1 | 5 | 0 | 19 | 683 | 98.70 | |
| HYDIN | 2.78 | 0.65 | 21 | 24 | 23 | 5 | 139 | 5005 | 98.48 | Gorilla |
| RIF1 | 2.77 | 0.62 | 9 | 9 | 11 | 5 | 59 | 2130 | 93.38 | |
| SMC1B | 2.76 | 0.70 | 1 | 6 | 4 | 0 | 33 | 1197 | 98.76 | Human |
| MTIF2 | 2.75 | 0.62 | 5 | 1 | 2 | 1 | 20 | 727 | 100.00 | |
| CDHR2 | 2.75 | 0.63 | 4 | 4 | 4 | 2 | 29 | 1056 | 99.91 | |
| FGD5 | 2.74 | 0.66 | 2 | 3 | 7 | 2 | 34 | 1241 | 96.20 | Human |
| PIEZO1 | 2.74 | 0.57 | 8 | 8 | 20 | 3 | 56 | 2045 | 87.69 | |
| NUB1 | 2.74 | 0.62 | 3 | 0 | 4 | 0 | 17 | 621 | 97.18 | Chimpanzee |
| AKAP13 | 2.73 | 0.57 | 13 | 8 | 23 | 9 | 74 | 2711 | 98.19 | Chimpanzee |
| EGF | 2.72 | 0.58 | 8 | 5 | 6 | 1 | 32 | 1176 | 99.41 | |
| PKHD1 | 2.72 | 0.57 | 22 | 15 | 34 | 6 | 105 | 3859 | 97.70 | |
| RNF17 | 2.72 | 0.61 | 5 | 5 | 10 | 5 | 44 | 1618 | 99.88 | |
| KIF26A | 2.71 | 0.57 | 9 | 9 | 7 | 5 | 43 | 1586 | 91.73 | |
| MOCOS | 2.71 | 0.63 | 4 | 1 | 5 | 1 | 24 | 886 | 99.77 | Chimpanzee |
| PLCH2 | 2.71 | 0.66 | 4 | 2 | 7 | 2 | 35 | 1293 | 95.07 | Chimpanzee |
| ANO7 | 2.71 | 0.59 | 2 | 4 | 3 | 3 | 22 | 813 | 96.10 | |
| USP45 | 2.70 | 0.61 | 5 | 3 | 2 | 1 | 22 | 814 | 100.00 | |
| NOL6 | 2.70 | 0.68 | 1 | 3 | 5 | 2 | 30 | 1111 | 96.95 | Human |
| ST14 | 2.70 | 0.63 | 1 | 2 | 6 | 1 | 23 | 852 | 99.65 | Human |
| TDRD1 | 2.70 | 0.62 | 2 | 4 | 7 | 3 | 31 | 1149 | 99.65 | Human |
| PDZD2 | 2.70 | 0.61 | 8 | 15 | 16 | 1 | 68 | 2523 | 97.11 | Human |
| DUSP27 | 2.68 | 0.58 | 6 | 9 | 4 | 0 | 31 | 1158 | 100.00 | |
| NCKAP5 | 2.67 | 0.56 | 12 | 9 | 11 | 4 | 49 | 1832 | 95.97 | |
| HR | 2.67 | 0.61 | 7 | 3 | 5 | 0 | 28 | 1047 | 93.07 | |
| PPL | 2.67 | 0.63 | 7 | 11 | 5 | 0 | 45 | 1683 | 98.13 | Gorilla |
| DTHD1 | 2.67 | 0.60 | 1 | 9 | 2 | 1 | 24 | 898 | 99.56 | |
| ZNF407 | 2.67 | 0.57 | 5 | 11 | 11 | 5 | 46 | 1722 | 99.54 | |
| NLRC4 | 2.67 | 0.69 | 2 | 1 | 4 | 0 | 23 | 861 | 99.88 | Chimpanzee |
| CRYBG1 | 2.67 | 0.59 | 8 | 14 | 11 | 1 | 53 | 1986 | 99.75 | |
| PLEKHG6 | 2.66 | 0.63 | 1 | 1 | 6 | 1 | 21 | 790 | 100.00 | |
| SPIDR | 2.65 | 0.58 | 2 | 1 | 8 | 3 | 24 | 904 | 99.12 | Chimpanzee |

| | | | | | | | | | | |
|---|---|---|---|---|---|---|---|---|---|---|
| CEP162 | 2.64 | 0.68 | 2 | 2 | 8 | 2 | 37 | 1400 | 100.00 | |
| ARHGAP11A | 2.64 | 0.67 | 4 | 2 | 3 | 1 | 27 | 1023 | 100.00 | Chimpanzee |
| ANKRD24 | 2.64 | 0.58 | 5 | 3 | 6 | 3 | 28 | 1062 | 94.74 | |
| MMRN2 | 2.63 | 0.61 | 4 | 3 | 3 | 1 | 22 | 838 | 100.00 | |
| BOD1L1 | 2.62 | 0.60 | 9 | 14 | 22 | 5 | 80 | 3048 | 100.00 | Human |
| CCDC18 | 2.62 | 0.60 | 3 | 6 | 11 | 2 | 38 | 1453 | 100.00 | Human |
| MAP4 | 2.61 | 0.59 | 3 | 4 | 3 | 4 | 25 | 957 | 92.46 | |
| LRIG1 | 2.61 | 0.66 | 2 | 3 | 5 | 1 | 28 | 1073 | 98.53 | Human |
| TANGO6 | 2.59 | 0.66 | 4 | 4 | 3 | 0 | 28 | 1082 | 98.90 | Gorilla |
| KIF20B | 2.59 | 0.57 | 7 | 7 | 11 | 8 | 47 | 1818 | 99.89 | |
| ELP1 | 2.57 | 0.61 | 5 | 8 | 6 | 0 | 34 | 1322 | 99.55 | |
| JCAD | 2.55 | 0.58 | 2 | 10 | 8 | 2 | 34 | 1332 | 98.38 | Human |
| CTC1 | 2.55 | 0.62 | 5 | 4 | 2 | 2 | 26 | 1019 | 87.92 | Gorilla |
| KIF14 | 2.55 | 0.66 | 7 | 5 | 4 | 2 | 42 | 1648 | 100.00 | Gorilla |
| SI | 2.54 | 0.71 | 1 | 6 | 6 | 2 | 46 | 1814 | 99.40 | |
| TEP1 | 2.52 | 0.61 | 13 | 16 | 7 | 2 | 63 | 2502 | 98.00 | Gorilla |
| URB1 | 2.51 | 0.60 | 9 | 6 | 12 | 7 | 56 | 2231 | 98.33 | Chimpanzee |
| MICAL1 | 2.49 | 0.59 | 7 | 2 | 7 | 0 | 27 | 1085 | 99.91 | Chimpanzee |
| NPAT | 2.48 | 0.61 | 3 | 4 | 10 | 2 | 34 | 1371 | 97.86 | Human |
| HMCN2 | 2.45 | 0.60 | 19 | 17 | 29 | 3 | 105 | 4290 | 93.98 | |
| FREM3 | 2.43 | 0.55 | 11 | 9 | 16 | 4 | 52 | 2137 | 99.95 | |
| BAHCC1 | 2.43 | 0.61 | 7 | 6 | 11 | 3 | 47 | 1932 | 84.92 | |
| CENPF | 2.43 | 0.53 | 18 | 17 | 24 | 3 | 74 | 3044 | 99.57 | |
| CTAGE5 | 2.41 | 0.60 | 5 | 2 | 12 | 1 | 34 | 1409 | 99.79 | Chimpanzee |
| FREM1 | 2.40 | 0.54 | 10 | 8 | 20 | 4 | 52 | 2167 | 99.59 | |
| FASN | 2.39 | 0.56 | 9 | 8 | 17 | 6 | 55 | 2304 | 95.32 | |
| LCOR | 2.38 | 0.66 | 6 | 1 | 7 | 2 | 37 | 1554 | 99.87 | Chimpanzee |
| TTC3 | 2.36 | 0.57 | 13 | 11 | 4 | 4 | 46 | 1946 | 97.79 | Gorilla |
| C2CD3 | 2.36 | 0.64 | 9 | 8 | 7 | 3 | 53 | 2243 | 97.61 | Gorilla |
| APOB | 2.36 | 0.51 | 27 | 28 | 37 | 9 | 107 | 4534 | 99.45 | |
| ICE1 | 2.35 | 0.53 | 13 | 7 | 17 | 6 | 52 | 2217 | 99.86 | |
| ZDBF2 | 2.34 | 0.56 | 14 | 11 | 12 | 3 | 55 | 2349 | 100.00 | |
| F5 | 2.32 | 0.58 | 8 | 7 | 11 | 7 | 50 | 2155 | 99.31 | |
| SYNE2 | 2.19 | 0.56 | 38 | 30 | 42 | 3 | 146 | 6678 | 98.31 | |

**S9 Table**: Higher than expected substitution rate on the #1# branch

| Gene | #1# % subs per site | #1# norm branch length | Human subs | Chimp subs | Gorilla subs | #1# subs | Gibbon subs | Align overlap | Align Sat | Lower in |
|---|---|---|---|---|---|---|---|---|---|---|
| SLC39A14 | 3.51 | 0.62 | 2 | 0 | 1 | 17 | 4 | 485 | 98.98 | |
| SUMO3 | 3.20 | 0.39 | 0 | 0 | 1 | 4 | 3 | 125 | 89.29 | |
| IL9 | 2.78 | 0.33 | 0 | 0 | 0 | 4 | 6 | 144 | 100.00 | |
| C16orf78 | 2.27 | 0.22 | 1 | 1 | 3 | 6 | 16 | 264 | 100.00 | |
| CCDC78 | 1.82 | 0.24 | 0 | 2 | 5 | 6 | 11 | 330 | 89.19 | |
| TMEM231 | 1.50 | 0.33 | 1 | 0 | 2 | 4 | 3 | 266 | 83.91 | |
| NDUFAF6 | 1.50 | 0.43 | 0 | 1 | 2 | 5 | 1 | 333 | 100.00 | Gibbon |
| ZNF778 | 1.34 | 0.20 | 5 | 5 | 3 | 8 | 18 | 598 | 98.36 | |

**S10 Table**: Evidence for rapidly diverging human genes

| Gene | Tissue-specific expression | Phenotype | Biological Process |
|---|---|---|---|
| ADCYAP1 | Biased expression in appendix and 12 other tissues | Schizophrenia | Ovarian follicle development, Behavioral fear response, and Inflammatory response |
| PVALEF | Low expression overall, highest in fat | - | - |
| PGLYRP1 | Restricted expression toward bone marrow | Blood protein levels | Immune response to bacterium |
| PSORS1C2 | Restricted expression toward skin | Autism spectrum disorder or schizophrenia | - |
| BTNL2 | Low expression | Sarcoidosis, Autism spectrum disorder or schizophrenia, Asthma, Blood pressure | Positive regulation of T cell proliferation and interleukin-2 secretion |
